# Regulation of somatic stem cell precursor fates and proliferation during *Drosophila melanogaster* pupal ovary development resembles the signaling framework for adult stem cell behavior

**DOI:** 10.1101/2025.05.08.652870

**Authors:** Rachel Misner, Amy Reilein, Helen V. Kogan, Daniel Kalderon

## Abstract

Follicle Stem Cells (FSCs) in the Drosophila melanogaster ovary are maintained through independent regulation of division and differentiation to become proliferative Follicle Cells (FCs) to their posterior or quiescent Escort Cells (ECs) to their anterior. These behaviors are guided by graded extracellular Hedgehog (Hh) and Wnt signals emanating from cells anterior to FSCs and an inverse gradient of JAK-STAT pathway activity. Here, we used lineage analyses to investigate the development of ECs, FSCs and FCs from a common set of precursors during pupation. Previous studies found that the most anterior precursors divide slowest, with quiescence spreading from the anterior over time to include all ECs, that FSCs are specified simply by their location at eclosion, and that the first FCs derive from cells that accumulate posterior to the developing germline over the first 48h of pupation. We now provide evidence that those accumulating cells derive from migration out of the developing germarium. We found that Wnt pathway activity favored conversion of precursors to more anterior adult derivatives (ECs rather than FCs), while JAK-STAT pathway activity favored posterior outcomes. Posterior bias and faster division, explored by altering Cyclin E activity, both favored a precursor becoming an FSC. Both JAK-STAT and Hh signaling could increase precursor division rate. All of these characteristics resemble regulation of adult FSC behavior. We suggest that similar signaling networks and division rate dependence during maintenance and development may be general features for stem cells that are specified in parallel with tissue development and that exhibit division-independent differentiation.

**Article Summary:** Drosophila ovarian Follicle Stem Cell (FSC) behavior and regulation by graded Hedgehog (Hh), Wnt and JAK-STAT signals is well understood, motivating us to study mechanisms guiding development of FSCs and the neighboring cell types they maintain. The marked progeny of single cells labeled at pupariation were compared in adults for lineages with genetic alterations affecting signal transduction. Wnt and JAK-STAT had strong opposing influences on the anterior-posterior location, and hence identity of derivatives, while JAK-STAT and Hh (via Yorkie) stimulated cell division. Conversion of precursors to FSCs was favored by higher division rates.

## Introduction

Many adult tissues are maintained by adult stem cells (Clevers and Watt 2018; Post and Clevers 2019; Kalderon 2022; Gao *et al*. 2023; Beumer and Clevers 2024). The two universal stem cell behaviors of cell division and initiating differentiation at appropriate, balanced frequencies are inevitably regulated by their physical and signaling environment, often known as the stem cell niche (Voog and Jones 2010; Biteau *et al*. 2011; Losick *et al*. 2011). Moreover, the further differentiation of immediate stem cell derivatives is often guided by additional cues from the mature adult tissue. Thus, stem cell function depends on the nature and spatial organization of the stem cells themselves, neighboring niche cells and more distant cells guiding full differentiation. The behavior and regulation of different types of adult stem cell is being studied productively for several paradigms. By contrast, the developmental mechanisms that place an appropriate number of stem cells in a suitable niche and larger tissue context are largely unknown. Understanding that process would have the potential benefit of facilitating the creation of self-sustaining organoids and exploring developmental abnormalities leading to impaired adult stem cell function or cancer susceptibility.

The Drosophila ovary is an excellent paradigm for studying the development of adult germline and somatic stem cells (Giedt and Tootle 2023; ST JOHNSTON 2023). Each adult ovary consists of 15-20 ovarioles, with an anterior germarium and a series of developing egg chambers of increasing maturity, with each ovariole producing a mature egg as often as every 12h (Fig. 1A) (Spradling *et al*. 1997; Duhart *et al*. 2017; Giedt and Tootle 2023). This formidable cell production is fueled by 2-3 Germline Stem Cells (GSCs) (Eliazer and Buszczak 2011) and about sixteen somatic Follicle Stem Cells (FSCs) (Reilein *et al*. 2017; Hayashi *et al*. 2020; Kalderon *et al*. 2021; Kalderon 2022). The known biology of these two types of stem cell includes numerous interesting contrasts. GSCs generally remain in place at the anterior of the germarium, are individually long-lived, maintained by single-cell asymmetry and depend on anti-differentiation signals from immediately adjacent niche cells (Cap cells) (Kahney *et al*. 2019; Zhang and Cai 2020; ST JOHNSTON 2023). The differentiation of the immediate “cystoblast” derivative of a GSC depends on interactions with non-dividing somatic Escort Cells (ECs) (Kirilly *et al*. 2011) during posterior migration and is manifest as a set of four synchronized divisions with incomplete cytokinesis through region 1 to form a sixteen-cell region 2 cyst.

**Figure 1:**
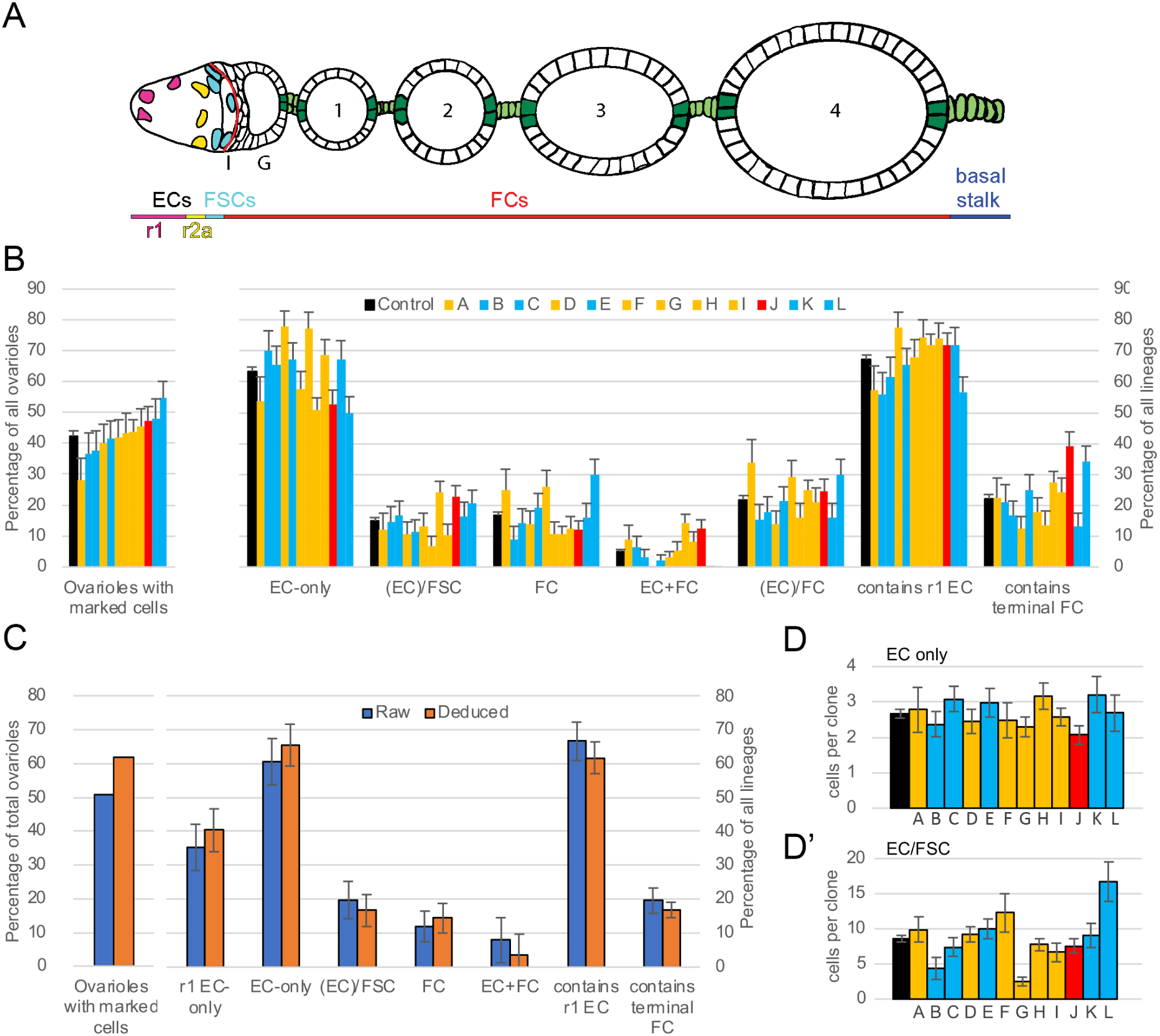
Estimation of single-cell lineage outcomes and cell numbers in control samples. **(A)** Illustration of somatic cells of an ovariole in a newly-eclosed fly. The germarium (anterior, left) produces budded egg chambers (posterior, right), four of which (numbered 1-4) are typically present in each ovariole in a newly-eclosed fly. Escort Cells (ECs) are divided according to their location, as region 1 (r1: magenta) and region 2a (r2a: yellow). FSCs (cyan) reside principally in two A/P rings, just anterior to the anterior border of strong Fas3 cell surface expression (“Fas3 border”: red line). FCs (mostly without a filled color), posterior to the Fas3 border, are classified by their location: Immediate (Region 2b, I), Germarium (Region 3, G), egg chamber 1, 2, 3, and 4. Specialized FC sub-types colored are polar cells (dark green), stalk cells (light green) and basal stalk cells (light green). **(B)** MARCM controls from 12 experiments, analyzed in newly-eclosed flies after 0h APF clone induction. Colors indicate the recombinant FRT site: 40A (yellow), 42D (blue), 82B (red). Black is the non-weighted average of all control values. The first cluster reports the percentage of all ovarioles with any marked cells. The number of marked ovarioles ranged from 36 to 121 (36, 41, 51, 54, 58, 56, 48, 121, 62, 98, 49, 65, respectively) and the total number of ovarioles from 102 to 277 (128, 112, 135, 135, 140, 134, 111, 277, 136, 207, 102, 119, respectively). Other clusters report the deduced frequency of the named lineage categories among marked lineages after estimating single-cell lineage frequencies (see methods). The number of FSC-containing lineages without ECs is generally very low and is therefore combined with lineages containing ECs and FSCs (written as (EC)/FSC); almost all of these lineages also contain marked FCs. Ovarioles with ECs and FCs but no FSCs are thought mostly to contain two lineages. The EC+FC category therefore gives an indication of the limitations of the method used to convert raw data to estimates of single-cell lineages. Here, the (EC+FC) and FC-only frequencies have also been added to give the (EC)/FC category. The final two clusters show the percentage of all lineages that includes at least one marked r1 EC or terminal FC. **(C)** An individual control from (**B**) (control C, third control) to show raw values (blue) compared to deduced single-cell lineage values (orange). **(D, D’)** Deduced average number of **(D)** cells per EC-only clone (total cell number: 54, 68, 103, 103, 116, 80, 85, 195, 110, 107, 106, 87, respectively) and **(D’)** EC plus FSC cells in EC/FSC clones (total EC + FSC cell number: 40, 24, 79, 56, 71, 116, 12, 266, 58, 194, 85, 281, respectively; the number of EC/FSC lineages for samples B and G are notably low) for all MARCM controls analyzed in newly-eclosed flies after 0h APF clone induction. (**B-D**) SEMs shown. For raw data and calculations please see supplementary spreadsheets titled 0h APF Graphs Fig1,3; 0h APF Controls Variation; Compilation of Numbers Figure 1 Controls.

FSC cell bodies reside principally in two adjacent rings along the anterior-posterior (A/P) axis, surrounding a central germline stage 2b cyst and bounded on the other side by the basement membrane lining the germarium (Fig. 1A) (Reilein *et al*. 2017; Kalderon 2022). FSCs can exchange A/P locations and move circumferentially. They are maintained by population asymmetry with independent cell division and differentiation, characterized by frequent and variable, apparently stochastic, loss and expansion of individual cell lineages (Reilein *et al*. 2017; Reilein *et al*. 2018; Kalderon 2022). This picture of FSC organization and behavior differs markedly from an earlier model as a result of more thorough investigation and discarding earlier expectations of single-cell asymmetry, which led to highlighting only a minority of all FSC lineages (Margolis and Spradling 1995; Kalderon 2022). FSCs are responsible for producing about six founder Follicle Cells (FCs) for each future egg chamber. A few of these FCs arrest division soon afterwards and form specialized polar cells at the anterior and posterior termini of each egg chamber, as well as adjacent stalk cells, which connect adjacent egg chambers and allow budding of an egg chamber from the germarium (Fig. 1A) (Grammont and Irvine 2001; ASSA-KUNIK *et al*. 2007; Dai *et al*. 2017). Other FCs continue division in growing egg chambers to form an expanding single-layer epithelium of about 650 cells by mid-oogenesis (stage 6). These FCs acquire different identities according to their location, guided by JAK-STAT signals produced in polar cells and EGFR-stimulating signals from the oocyte, initially at the posterior of the germline cyst, and then from the oocyte nucleus after migration to an anterior cortical location that defines the future dorsal aspect of a developing embryo (Mcgregor *et al*. 2002; Xi *et al*. 2003; Merkle *et al*. 2020; ST JOHNSTON 2023). FSCs also produce new ECs in adults, albeit at a much lower rate than FCs (Reilein *et al*. 2017; Melamed and Kalderon 2020). The A/P location and differentiation of FSCs are guided principally by long-range signals, emanating from the anterior (multiple Wnt ligands from Cap cells and ECs) and posterior (JAK-STAT ligand Unpaired from newly- differentiating polar cells) to produce inverse gradients of those pathway activities across the stem cell domain, effectively defining the A/P extent of the FSC domain (Reilein *et al*. 2017; Melamed and Kalderon 2020). JAK-STAT pathway activity also stimulates FSC division in a graded fashion (Melamed and Kalderon 2020; Melamed *et al*. 2023). Hedgehog pathway activity also stimulates FSC division but may not contribute to the spatial pattern of FSC division (Vied and Kalderon 2009; Huang and Kalderon 2014). Other factors supporting the much faster division of posterior FSCs remain to be identified. The division rate of posterior FSCs approximately matches the rate of their conversion to FCs, so the exchange of FSCs between anterior and posterior layers is roughly at equilibrium (Melamed *et al*. 2023).

During development, germline and somatic gonadal cells are specified separately in the embryo and coalesce after germ cell migration to form an embryonic gonad (Boyle and Dinardo 1995; Murray *et al*. 2010; JEMC 2011). During larval stages, the ovary develops from anterior to posterior (initially termed apical to basal), starting with the specification of somatic Terminal Filament (TF) cells, which resolve into separate intercalated stacks that seed each individual ovariole (Godt and Laski 1995; Gilboa 2015). A Notch signal from TF cells recruits adjacent cells from a pool of somatic cells (Intermingled Cells: ICs) intermingled with germ cells, to become quiescent Cap cells (Ward *et al*. 2006; Song *et al*. 2007; Panchal *et al*. 2017; Zamfirescu *et al*. 2022). Germ cells that contact Cap Cells become the future GSCs, with other germ cells, in more posterior locations, initiating differentiation soon afterwards (Zhu and Xie 2003; Asaoka and Lin 2004; Gancz *et al*. 2011; Gancz and Gilboa 2013). As Cap cells and GSCs form in third instar larvae, somatic Swarm Cells migrate basally (towards the future posterior) around the TF cells to separate sets of developing ICs and germ cells into individual units (Banisch *et al*. 2021). Then, during pupation, germ cells sequentially mature (towards the posterior) into sixteen-cell cysts, while somatic ICs amplify. These IC precursors eventually become ECs, FSCs and FCs, such that the newly-eclosed adult has a mature germarium structure, followed by four egg chambers of increasing size, allowing egg-laying to commence within about a day (Reilein *et al*. 2021).

The specification of different ovarian somatic cell types during pupation has been studied by lineage analyses and direct morphological observation, including live imaging, yielding three major conclusions. First, FSCs are not specified at a particular time during development; instead, single-cell lineages generally include ECs and FCs together with FSCs, with the latter defined simply by occupying the appropriate A/P locations at eclosion (Reilein *et al*. 2021). Thus, in this paradigm, the stem cells are not set aside early in development but derive from the same process that forms the whole adult tissue. Another study had suggested separate precursors for ECs and FSCs at the start of pupation, based on lineage analysis initiated by two GAL4 drivers (*Con-GAL4* and *bond-GAL4*) together with temperature sensitive GAL80 (Slaidina *et al*. 2020). That suggestion was, however, based on assuming that the additional presence of marked ECs in most adult ovarioles with marked FSCs was due to the presence of two or more lineages. Sparse induction of cell lineages showed this explanation to be incorrect; FSC-containing single-cell lineages almost always included ECs, revealing a common origin (Reilein *et al*. 2021). Moreover, using the same reagents, it was apparent that neither the frequency, nor the time of lineage initiation could be controlled satisfactorily using the temperature-sensitive GAL80 technique, contrasting with lineage initiation using a heat-shock to activate recombinase encoded by a *hs-flp* transgene (Reilein *et al*. 2021). Second, anterior somatic cell precursors divide more slowly than posterior precursors and terminate division during pupation. This feature appears to limit the number of the most anterior derivatives, Escort Cells (Reilein *et al*. 2021). Third, the formation of the first FCs and the first egg chamber is morphologically very different from the process in adults, where newly-formed FCs are adjacent to a germline cyst and that germline cyst is subsequently surrounded by a monolayer of FCs before budding from the posterior of the germarium. Instead, many somatic cells accumulate posterior to the germarium (harboring all germ cells) during the first half of pupation and are then invaded by a germline cyst released from the germarium and migrating a considerable distance towards the posterior (Reilein *et al*. 2021).

We would like to understand what orchestrates the movements and proliferation of somatic cells during pupation required to adopt appropriate adult identities and assemble a morphologically normal adult ovariole. As a first step, we have investigated cell-autonomous responses of somatic precursors to genetic alterations by using lineage analyses during pupation to measure final outcomes as ECs, FSCs and FCs. We began by investigating signals and cellular properties known to influence the behavior of adult FSCs. This includes examining signals instrumental to FSC maintenance and differentiation (Wnt and JAK-STAT), signals and effectors important for FSC division (JAK-STAT, Hedgehog (Hh) and Yorkie (Yki)), and the effect of cell division rate on cell fate (Zhang and Kalderon 2001; Song and Xie 2003; Vied and Kalderon 2009; Wang and Kalderon 2009; Vied *et al*. 2012; Wang *et al*. 2012; Huang and Kalderon 2014; Wang and Page-mccaw 2014; Reilein *et al*. 2017; Hayashi *et al*. 2020; Melamed and Kalderon 2020; Melamed *et al*. 2023). Among adult FSCs, there is competition for residence in the FSC domain, with conversion of FSCs to FCs or ECs balanced by FSC amplification through cell division. Hence, maintenance and amplification of an individual FSC lineage is cell-autonomously favored by genetic alterations that either increase the division rate or reduce the frequency of differentiation (Reilein *et al*. 2017; Reilein *et al*. 2018). We found that this principle and the roles of Wnt and JAK-STAT signaling apply also to the pupal development of FSCs, while responses to Hh and Yki are at least partly shared for stem cell development and adult maintenance. A key source of anterior signals (Wnt, Hh) is present at the start of pupal development. The original source of the ligand activating the JAK-STAT pathway, which guides FC formation from the start of pupation, is not defined but likely has a posterior bias; the major source, which instructs a robust gradient of pathway activity, emerges after the first egg chamber forms midway through pupation. These experimental findings and their rationalization suggest principles that may be shared generally among adult stem cells with division-independent differentiation that is guided by location: early establishment of niche signaling centers, continuous competition between division and differentiation for an individual cell to become or remain a stem cell, and regulation of division and cell location by the same signaling networks during adult stem cell development and maintenance.

## Materials and Methods

### Clone induction via MARCM

GFP-marked MARCM clones (Lee and Luo 2001) were induced on different chromosome arms (2L, 2R, and 3R) using flies of the genotypes listed below (“X” and “Y” identities are listed subsequently; “*>*” indicates an FRT element, “*tub*” refers to the *αtub84B* promoter (Lee and Luo 2001) and “*act*” to the *act5C* promoter (Pignoni and Zipursky 1997)). In all cases, recombination is initiated at *FRT* sites by heat-shock induction of FLP recombinase, leading to marked clones lacking a *Gal80* transgene and expressing *UAS-GFP* from *tub-GAL4* and *act>GAL4* drivers. Simultaneously, a recessive allele (“X”) becomes homozygous or a transgene (“*UAS-(Y)*”) initiates expression, or both, or neither (controls).

(2L) *hs-FLP UAS-nGFP tub-Gal4*; *(X) FRT40A/tub-Gal80 FRT 40A*; *UAS- (Y)/act > CD2 > GAL4*

(2R) *hs-FLP UAS-nGFP tub-Gal4*; *FRT42D (X) / FRT 42D tub-Gal80*; *UAS- (Y)/act > CD2 > GAL4*

(3R) *hs-FLP UAS-nGFP tub-Gal4*; *+ / act > CD2 > GAL4, UAS-GFP*; *FRT 82B (X) / FRT 82B tub-Gal80*.

Studies conducted at 29C utilized flies of the genotype below (where an additional *act-GAL80* element on 2R is necessary to silence GAL4):

(2R) *hs-FLP UAS-nGFP tub-Gal4*; *FRT42D (X) / FRT 42D act-Gal80 tub-Gal80*; *UAS-(Y)/act > CD2 > GAL4*.

For 2L chromosome studies, (X) included *NM* (control), *cycE^wx^*, *smo^2^*, *pka-C1^H2^*, *smo^2^ pka-C1^H2^*, *cutlet^4.5.43^*, and (Y) included *DIAP1*, *CycE*, *Yki^S168A^*; also, a *Su(fu)^LP^/ Su(fu)^LP^* background was used for *smo pka* clones.

For 2R chromosome studies, (X) included *sha* (control), *ptc^S2^*, *arr^2^*, *yki^B5^*, *ptc^S2^ yki^B5^*, *hpo^42-47^* and (Y) included *Hop* together with *UAS-Dap*, as well as a *tub-yki* transgene that does not include the *UAS* promoter.

For 3R chromosome studies, (X) includes *NM* (control), *apc1^1^ apc2^2^*, *axn^E77^*, *kibra^del^*, *wts^x1^*, and *stat^85C9^*.

### Lineage analysis by MARCM in newly-eclosed flies

From the time that pre-mated adults were put into a fresh vial (for 24h or less) to eclosion of the first several adult progeny, sufficient for analysis, was most commonly 11 days (Reilein *et al*. 2021). Timing of heat shocks to initiate lineages was determined with this 11-day course in mind. The larval-to- pupal transition, where third-instar larvae are crawling up the sides of the vial and beginning puparium formation, was generally 5 days before eclosion. Hence, adults collected 5d after heat-shock were considered to have lineages initiated at pupariation (“0h APF”). At the time of heat-shock (generally 6d after parents were first added to the vial), some animals were at pupariation; others were slightly more advanced or delayed. Vials were cleared of any eclosed adults the evening before collecting newly-eclosed adults over a period of no more than 6h centered on 120h after heat-shock. Similar procedures were used for collection of adults with lineages designated as initiated 2 days prior to pupariation (“-2d APF”: heat-shock 4d after introducing parents; collect adult progeny over 6h, centered on 168h after heat-shock) and 36h after pupariation (“36h APF”: heat-shock 7.5d after introducing parents; collect adult progeny over 6h, centered on 84h after heat-shock). Vials were heat shocked at 33C for 12 min (0h APF, 36h APF) or 20 min (-48h APF). These conditions were chosen as the mildest conditions consistently inducing a similar yield of marked lineages at 0h APF, so that many ovarioles would harbor a single marked lineage. To achieve a similar result, a longer period of heat-shock was used at -48h APF because fewer precursors are present. Afterwards, vials were incubated at 25C or 29C. Higher temperature increases GAL4 activity and was used for expression of *UAS-Hop* and controls for that test (analogous tests in adult ovaries showed that 29C is necessary to achieve strong activation of the JAK-STAT pathway (Melamed and Kalderon 2020)); all other flies were incubated at 25C or room temperature (23C-25C). Flies were dissected on the day of eclosion (0d).

### Adult ovary fixation and staining

Adults were dissected in PBS using forceps by separating the entire posterior region of the abdomen, revealing the ovaries within. Non-ovary material, such as fat bodies and other organs, was gently shaken or teased out with forceps, and the ovaries (still attached to the cuticle) were transferred using forceps to tubes containing 400 μl fixative. Ovaries were fixed in 4% paraformaldehyde in PBS for 10 min at room temperature, rinsed 3x in PBS with 0.1% Triton and 0.05% Tween-20 (PBST), and blocked in 10% normal goat serum (Jackson ImmunoResearch Laboratories) in PBST. Ovarioles were incubated in primary antibody for 1 hr. Ovarioles were rinsed in PBST 3x and incubated for 1 hr in secondary antibodies diluted 1:1000. Ovaries were rinsed 2x in PBST and 1x in PBS for 5 min each.

Ovaries were transferred using a p1000 with the narrow portion of the tip cut off to a well containing PBS. The cuticle was then separated from the ovaries, which were transferred using forceps to a slide containing a drop of PBS. Each slide accommodates two drops, and each drop contained the ovaries from one individual animal. Ovaries were gently teased apart with extra sharp forceps, separating into clusters of 1-4 ovarioles, taking care to ensure that entire ovarioles remained intact and did not separate, as tracing the clones throughout the entire ovariole is crucial. After separation was complete, 25 μl of Flouromount-DAPI was applied to a coverslip and carefully placed over the portion of the slide with each sample.

### Immunohistochemistry

Monoclonal antibodies against Fasciclin III (Fas3) and Vasa were obtained from the Developmental Studies Hybridoma Bank, created by the NICHD of the NIH and maintained at The University of Iowa, Department of Biology, Iowa City, Iowa 52242. 7G10 anti-Fasciclin III was deposited to the DSHB by C. Goodman, and was used at 1:250. Anti-Vasa (used at 1:10) was deposited by A. C. Spradling/D. Williams and used for staining adult ovaries. For pupal ovaries, the monoclonal rat anti-Vasa was not effective and instead rabbit anti-Vasa (gift from R. Lehmann, MIT) was used. Other primary antibodies used were anti-GFP (A6455, Molecular Probes) at 1:1,000; and goat FITC-anti-GFP (Abcam ab6662) at 1:400. Secondary antibodies were Alexa-488, Alexa-546, Alexa-594 or Alexa-647 from Molecular Probes. DAPI-Fluoromount-G (Southern Biotech) was used as mounting medium for all experiments. Images were collected with a Zeiss LSM700 or LSM800 laser scanning confocal microscope (Zeiss) using a 63x 1.4 N.A. lens.

### Scoring and Analysis of marked lineages in adult ovaries

Ovaries were stained using antibodies to Vasa, Fas3 and GFP. Vasa staining assists in classifying cysts of regions 1 (fewer than 16 cells) versus regions 2a and 2b (16-cell cysts). The anterior border of strong Fas3 expression corresponds to the border between FSCs and FCs, with cells immediately adjacent and anterior to this boundary, together with their immediate anterior neighbors (three rings of cells anterior to the boundary of strong Fas3 staining), classified as FSCs. Escort cells were scored as r1 or r2a based on measurements that were performed with wild-type ovaries to determine the lengths of these regions. We previously examined EdU incorporation in ovaries of newly -eclosed flies and measured the distance from cap cells to the end of region 1 as determined by the presence of cysts that incorporated EdU. Region 1 cysts are going through mitosis and Region 2a cysts have the complete complement of 16 germ cells. The length ratio of regions 1:2a is about 3:1 in newly-eclosed adults. To measure the percentage of each terminal egg chamber covered by marked FCs, the ratio of GFP-marked cells to unmarked cells (stained with DAPI) was estimated throughout all z-sections.

The total number of ovarioles containing any GFP-positive cells (“clones” or “lineages”) was recorded, along with the total number of ovarioles, for each coverslip, which has a pair of ovaries from one animal. Each ovariole was scored for GFP-positive cells by counting the number of r1 ECs, r2a ECs, and FSCs. FCs were scored for presence (yes or no) in the following regions: “Immediate Daughter” (region 2b), “Germarium” (region 3), “1”, “2”, “3”, or “4” (corresponding to the four egg chambers typically present in ovaries of newly-eclosed flies; sometimes “5” also), and “BS” (Basal Stalk; a cluster of about 10 cells around the polar cells at the posterior of the terminal, most mature, egg chamber). Notes were also added if labeled cells of a specific genotype were present at unusual frequencies in certain locations, including polar cells and stalk cells. Each scored ovariole was then labeled with its broad clone type (EC, EC/FSC, EC/FSC/FC, EC/FC, FSC, FSC/FC, or FC). This classification, together with the exact number of GFP-labeled r1 and r2a ECs, FSCs, and the presence or absence of FCs in the regions listed above, was recorded in each row of a spreadsheet for a given genotype in an experiment. For each genotype in an experiment, we aimed to score at least 50 marked ovarioles across at least three coverslips. The data from each such spreadsheet were aggregated in order to measure several parameters, as described in the next section.

### Estimating single-cell lineage outcomes underlying raw observations

Although we used a mild and brief heat-shock in order to induce only low activity of a *hs-flp* transgene, it is inevitable that some adult ovarioles will include lineages from more than one marked precursor. Previously, two methods were considered for estimating single-cell lineage outcomes from raw observations (Reilein *et al*. 2021). One assumed that every marking event is independent (zero-clone frequency method, below). Another used information about the frequency of ovarioles with marked ECs and FCs (but no marked FSCs) as a likely indicator of the presence of more than one lineage. For wild-type lineages, the latter method (of the two) indicated a higher frequency of ovarioles with more than one lineage for a given sample, and was the preferred method (because we believed it to be more accurate). Here, that method cannot be used because different genotypes may greatly alter EC and FC frequencies, as well as producing single lineages with only marked ECs and FCs (from selective loss of prospective FSCs). Past experience suggested that the “zero-clone” frequency method used here will underestimate the number of ovarioles with more than one lineage. An indicator of that potential error is the proportion of deduced single-cell lineages with only EC and FCs; the average frequency of that lineage category was 5% for controls. However, during the course of the studies reported here we learned that lineages with just ECs and FCs can arise from loss of FSCs from a single EC/FSC/FC lineage rather than representing simultaneous EC-only and FC-only lineages. Thus, the error in estimating single-cell lineage frequency and composition by the method described below is less than initially anticipated and overestimated by the EC/FC frequency.

### Zero-clone frequency method

We assume all marking events are independent and use binomial calculations to derive the expected frequency of ovarioles with 1 and 2 lineages by deriving the probability of a recombination event in each precursor cell.

Although we need to use an estimate of the total number of cells present at the time of recombination, the numerical outcomes are not very sensitive to this exact number.

From prior published work (Reilein *et al*. 2021), lineage data estimated 24 or 17 precursor cells at pupariation (different experiments and labeling techniques). The number of precursor cells directly observed was 38 at 12h APF and 51 at 24h APF. This leads to an estimated number at 0h APF (assuming similar expansion from 0-12h and 12-24h APF) of 38 x 38/51= 28 cells

Since MARCM (the same technique used here) was used to deduce 24 precursors, we use that number.

If the frequency of ovarioles with no marked cells = f and p is the probability of recombination in a single cell, f = (1-p)^24^

The proportion of ovarioles with a single clone will be 24p(1-p)^23^ = f(24p)/(1-p).

Hence, the frequency of single-cell clones among labeled ovarioles = f(24p)/((1-p)(1-f)).

The frequency of ovarioles with 3 or more clones (estimated by this method) will likely be <20% of remaining ovarioles. Hence, we approximate, for simplicity, that all of the remaining ovarioles (non-zero and non-single cell clones) contain two clones. We also round up to a whole number for the estimated percentages of ovarioles with zero, 1 or 2 lineages (adding to 100).

For lineages induced 48h before pupariation, we previously estimated there might be 10 precursor cells.

So, f = (1-p)^10^

and the frequency of single-cell clones among labeled ovarioles = f(10p)/((1-p)(1-f)).

### Estimating single-cell lineage type frequencies

After establishing the estimated number of ovarioles with one and two lineages for a given experimental sample, we deduce the total number of single-cell lineages present and then estimate the fraction of those of specific types. The major types are outlined first and then sub-divisions are brought in.

We start with raw data of major phenotypes: EC, EC/FSC, FSC, FC, EC+FC

Single-cell clone frequencies are named: EC(q), EC/FSC (r), FSC(s), FC (t), EC+FC(m)

Single lineage frequency = p1, two-lineage frequency = p2

Observed EC only = q.p1 + q^2^.p2 = q^2^.p2 + q.p1

Hence, we can calculate q directly

Observed FC only = t.p1 + t^2^.p2 = t^2^.p2 + t.p1

Hence, we can calculate t directly

Observed FSC only = s.p1 + s^2^.p2 + 2s.t.p2 = s^2^.p2 + s(p1+2.t.p2)

Observed EC+FC = m.p1 + m^2^.p2 + 2.p2.m(q+t) + 2.p2.q.t

= m^2^.p2 + m (p1+2.p2. (q+t)) + 2.p2.q.t Observed EC/FSC = 1- (q+s+t+m)

Then, for r1 (a) and terminal FCs (z)

Observed r1 only = a.p1 + a^2^.p2 = a^2^.p2 + a.p1

Observed “Terminal FC only” = z.p1 + z^2^.p2 = z^2^.p2 + z.p1

Note-here, “Terminal FC only” means there are marked cells in the terminal egg chamber and there are no marked cells other than FCs.

r1 ECs + FC (b) and EC + terminal FC only (y) can also be inferred.

Observed r1+FC = b.p1 + b^2^.p2 + 2.p2.b(q+t) + 2.p2.a.t

= b^2^.p2 + b (p1 + 2.p2 (q+t)) + 2.p2.a.t

Observed EC + terminal FC only = y.p1 + y^2^.p2 + 2.p2.y(q+t) + 2.p2.q.z

= y^2^.p2 + y (p1 + 2.p2 (q+t)) + 2.p2.q.z

The above two equations exclude EC + FC with no r1 only or terminal FCs but that is a very small error.

For “only Term FC”, x, meaning no marked cells (even FCs) anterior to the penultimate egg chamber, and a similar category of EC + only Terminal FC, w, there are analogous equations.

Observed “only Terminal FC” = x.p1 + x^2^.p2 = x^2^.p2 + x.p1

Observed EC+ only Terminal FC = w.p1 + w^2^.p2 + 2.p2.w(q+x) + 2.p2.q.x

= w^2^.p2 + w (p1 + 2.p2 (q+x)) + 2.p2.q.x

### Added categories

(a) **“Normalized” numbers to acknowledge that EC+FC category sometimes includes an EC-only and an FC-only lineage.**

We acknowledge that the zero-clone frequency method might underestimate multiple clone frequency. Hence, the EC+FC category may include some EC-only and FC-only clones.

However, it is (i) reasonable that some prospective FSCs might be lost for WT from EC/FSC/FC lineages to give EC+FC lineages, (ii) genotypes with reduced division rate may cause selective loss of FSCs (this work), converting prospective EC/FSC/FC to EC+FC lineages. This may also occur to a degree for all genotypes. So, taking EC+FC as possibly composite (2 lineages) should be avoided if there is a division defect and is likely an overestimate of composite clones in all cases.

Method: assume all EC+FC (fraction m) are really single EC and single FC clones. Hence, m is added to EC-only & FC-only proportions (same for EC+FC specifically with term FCs or r1 ECs) and the total clone number is multiplied by (1+m) to give “normalized” EC and FC values.

These values appear in supplementary spreadsheets but are not discussed in the text.

**(b)** Clones with any terminal FC or r1 EC representation.

This is useful for considering cell allocation to the most extreme A/P locations (independent of other issues).

After deriving single-cell lineage frequencies,

Observed raw # of ovarioles with marked terminal FCs = “ObsTerm” Raw ovarioles with clones = x

Deduced fraction of ovarioles with two lineages = f

Fraction of single-cell lineages with marked terminal FCs = T

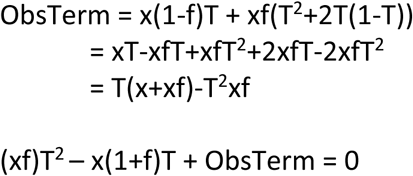

Quadratic solution = ((x(1+f)-SQRT[(x+xf)^2^-4xf.ObsTerm]))/(2xf)

Same for r1, using Obsr1 to calculate R

### Criticisms/shortcomings

If clone frequency varies more from fly to fly than ovariole to ovariole (within the same fly), then these corrections ought theoretically to be applied to each fly/coverslip. However, this is impractical (the methods cannot work well when n is low). To minimize this deficiency, it is best if all scored coverslips have roughly similar clone frequencies. We therefore instituted the criterion that coverslips (containing ovaries from a single fly) scored for clone phenotypes should have ovarioles with labeled cells at a frequency of 75% or lower (there were one or two experiments where we included coverslips with higher frequencies because there was no choice). We also instituted a minimum clone frequency of 15%. This was for two reasons. First, so that all clone frequencies were in a narrower range, so that errors due to single-cell lineage estimations would not vary widely among samples.

Second, we had observed previously that there is occasional cell marking in the absence of deliberate heat-shock. Such lineages arise at unknown times (if originating from recombination; most appear to be single EC marks of unknown origin). Such background is less than 5% and has minimal impact if deliberately-induced lineages are prevalent. For controls, the average frequency of marked ovarioles scored was 42% at pupariation and 63% 48h before pupariation.

Because any error in single-cell lineage estimations likely depends on clone frequency, we experimented with splitting each control sample into two of similar number, with the lowest lineage frequency flies segregated from the highest lineage frequency flies. We processed the data separately and then aggregated. The final outcomes were almost identical, so we treated all fly ovaries from one experiment as a single batch for single-cell lineage estimations.

### Terminal FC labeling with or without other FCs

“Only terminal FC” means labeled cells in the basal stalk and/or terminal egg chamber (which may extend to the penultimate egg chamber) but nowhere else in the ovariole.

“FC-only with terminal FCs” includes the category above, but also includes lineages with terminal FCs and additional FCs anterior to the penultimate egg chamber, but no ECs or FSCs.

Subtracting the category above from “FC-only” gives the FC-only lineages that do not include any labeled terminal FCs.

Thus, within the FC-only category, frequencies can be derived for only Terminal FCs, no terminal FCs, and a mix of terminal and non-terminal FCs.

These calculations were performed only for control lineages in order to determine if there were common precursors of FCs of terminal and non-terminal egg chambers.

### Cells per clone: EC-only lineages

We considered only EC-only, and not EC+FC, raw data because some of the latter may derive from EC/FSC/FC clones that have lost FSCs.

We calculated raw ECs per EC-only clone (and used the same principle for r1 ECs only) and multiplied by the ratio of raw EC-only clone frequency to calculate single-clone EC-only frequency. Because this is a derived number, we use raw observations of EC number in EC-only clones to estimate standard deviation (and then, standard error).

### Cells per clone: EC/FSC/(FC) lineages

Single-cell EC/FSC clones will all be within this raw data category. This is an inclusive category-most clones have FCs but those without FCs are also included. However, (a) such clones in raw data will sometimes also include an EC-only or an FSC-only clone and (b) the correct number of EC/FSC clones deduced by single-cell lineage estimations should be used.

Single-clone EC/FSC frequency = r

Observed EC/FSC = r.p1 + r^2^.p2 + 2.p2.r.(q+m) + 2.p2.r.s + 2.p2.(q+m).s

(a) Subtractions from total counted cells

Additional ECs come from EC only clones at (average cells per EC-only clone)(total number of single clones= actual clone number plus deduced number of double-clones)(2p2.(q+m).r)

Additional ECs come from EC only clones at (average cells per EC-only clone)(total number of single clones= actual clone number plus deduced number of double-clones)(2p2.s.r)

(b) Total number of single-cell EC/FSC clones in observed EC/FSC samples

= r. total number of single-cell clones in sample

= r. total clones in sample (1+p2)

Because the final estimates are derived numbers, we use raw observations of EC and FSC number in EC/FSC clones to estimate standard deviation (and then, standard error)

All of the above calculations are in the supplementary spreadsheet that includes all samples for lineages induced at different times: “Single-cell 0hAPF calculations” and “Single-cell -2dAPF calculations”.

### Evaluation of Controls Expected variability

If we assume that we are examining single-cell clones and that the chance of hitting each type (location) of cell in a given developing ovariole is independent (and given by the average frequency of that clone type (p) in a sample with n single clones), we expect to see a binomial distribution of the frequency of outcome p.

We therefore calculated the “expected” SEM for this process for each average frequency of lineage category over all 12 controls for lineages induced at pupariation. For binomial distributions the expected standard deviation= SQRT[np(1-p)]

We also calculated the observed SEM among the 12 individual values for each category.

The ratio of the two values (observed/expected) was mostly in the range of 1-2, consistent with the random sampling (by recombination events) of the starting A/P location of precursors being a major contributor to variation.

These data are presented in the supplementary spreadsheet “0h APF Controls Variation”.

### Significant Differences

The significance of differences for the frequency of a specific type of lineage for an experimental sample versus controls was estimated using Fisher’s exact two-tailed test. The data on the spreadsheets described above was copied (as values) on to separate spreadsheets to add some extra summary categories. Calculated p values (from an online calculator) were then added when these were either very low or of importance to acknowledge as not being low (i.e., values above 0.05 were generally not entered). Altogether, we tested the following types of condition as both reduced or increased by genetic manipulation- cell division rate, Wnt signaling, JAK-STAT signaling, Yki activity, and Hedgehog signaling, translating to 10 broad conditions. It may therefore be of special interest to know when p values are below 0.05 and below 0.005, so we designated those values by “*” and “**” in graphs. Supplementary spreadsheets with these p values are named “0h APF Significance Fisher” and “-2d APF Significance Fisher”.

### Inferring clone types, FSC and EC numbers in newly-eclosed adults from scoring 2d-old adults

In order to make quantitative conclusions about the developmental process up to eclosion from scoring clonal outcomes in 2d-old adults we made the following assumptions and adjustments. We assumed that EC numbers and locations did not change during this 2d interval because ECs generally appear to be quite long-lived. ECs are, however, produced from FSCs during adulthood, so it is likely that a small number of marked ECs scored at +2d were produced from FSCs after eclosion. The proportion of marked ECs arising from FSCs is very likely small based on known rates of FSC conversion to ECs and the observed high frequency of FSC clones with no ECs (44%) from control -2d samples. Individual marked FSCs can be lost, mainly by becoming FCs, or amplified at high rates during adulthood. If one or more marked FSCs were present at 0d but became FCs by +2d those marked FCs would reside around the two germarial cysts or in the two youngest egg chambers because egg chambers bud roughly every 12h, allowing four cycles of FC recruitment over 2d. We therefore scored an ovariole as containing an FSC clone (FSC-only or FSC/EC, according to EC content) if there were any marked FCs up to and including the second egg chamber, even if there were no marked FSCs at +2d. In those cases, we scored the number of marked FSCs as zero. On average, the number of marked FSCs should remain constant from 0d to +2d, so we should obtain a very good estimation of FSC numbers at 0d by scoring the numbers of marked FSCs at +2d, provided we include all examples where marked FSCs were lost altogether as containing zero FSCs. By using these guidelines, we could surmise the exact number of different types of lineage at 0d and the exact number of constituent ECs and FSCs from data collected by scoring clones in 2d-old adults.

### Dissection and staining of pupal ovaries

Pupae were removed from the vial wall by adding a drop of water and transferred with forceps into a well of a Corning PYREX glass spot plate (Corning, cat. no. 7220–85) containing either PBS or 4% paraformaldehyde solution with care taken not to damage or pierce the posterior end of the pupa. If the dissection was done in fixative solution, it could take no longer than 10 min (ovaries were rocked in clean fixative for a minimum of 15 min, but for no longer than 30 min). Ovaries from 72h APF and older had fewer tears in them when dissected directly into fixative. Using a dissecting microscope, pupae were held against the bottom of the well using forceps and the posterior tip was cut off using Vannas Spring Scissors (2.5mm straight edge, Fine Science Tools, Foster City, CA, item number 15000-08) about a third of the way in from the posterior end. The anterior part of the pupa was removed to a discard well. The posterior end was searched for ovaries (clear, spherical, striated structures) by gently dislodging fat tissue surrounding the ovaries so as not to cloud the well with debris. This was done by gently tearing apart the contents of the pupal case or swirling them in the well using two pairs of forceps. As soon as they were spotted, ovaries were transferred to a new well of fixative by the following process. A 20μl pipette tip with a pipettor set to 20μl was coated in 10% Normal Goat Serum (NGS) by pipetting up and down several times. The pipettor was set to 5μl to pipette up the ovary from the dissection well to transfer into the fixative well. Ovaries were fixed by rocking in the well for 15 min at room temperature. Fixative (and all liquid in subsequent steps) was removed in either of two ways: with a 1000μl pipette set to 270μl to slowly and carefully pipette up the liquid; or, using the same method to transfer ovaries above, with a 20μl pipette tip with a pipettor set to 20μl, which was coated in 10% Normal Goat Serum (NGS) by pipetting up and down several times. The pipettor was set to 5μl to pipette up the ovary from the fixative well to transfer into a fresh well with PBST. Ovaries were rinsed 3x in 2% PBST 0.5% Tween solution for 5, 10, and 45 min at 4C by rocking very slowly in a horizontal plane. The glass dissection dish was covered with a large pipette tip box. Ovaries were rinsed for 30-60 min in 10% NGS solution at 4C, and then rocked gently in primary antibody overnight. Ovaries were rinsed 3x with 0.5% PBST and then incubated for 1 hour (covered, cold room, rocker set on 2) in secondary antibody diluted 1:1000 with 0.5% Triton. Ovaries were rinsed 2x with 0.5% PBST and 1x with PBS. To mount ovaries, a 20μl pipette tip coated in 10% NGS solution, as described above, was set to 5μl and used to capture the ovary and transfer to a glass slide. 20-25μl of DAPI Fluoromount was added to a coverslip, and then placed on top of the slide with the ovaries. Care was taken to avoid pressing the coverslip and potentially disfiguring the mounted ovaries. In some studies of older pupal ovaries, specifically 72h APF and onwards, forceps were used to gently tear apart the ovarioles to separate germaria for clearer imaging. A sharpie was used to draw arrows to mark the location of the ovaries on the coverslip.

### Staging of pupae (Fz3-RFP experiments)

Third-instar larvae were sorted to select females, transferred to a vial with food and checked every hour to mark the time each individual developed into a puparium (white, but immobile, with small anterior spiracles). This time is 0 hours APF (after puparium formation). We found that keeping animals in the light at night and in the dark during the day on a 12h/12h cycle, allowed more of them to pupate during the daytime. We used a programmable outlet timer (Nearpow) together with a manual LED soft white nightlight (Energizer cat. 37099) installed in a dark incubator at 25C.

### Imaging and quantitation of Fz3-RFP profiles

Fz3-RFP flies (made by Ramanuj DasGupta, Genome Institute of Singapore and a gift from Erika Bach, NYU, USA) were dissected and stained for Fas3 and DAPI. Ovaries were imaged with a 63 × 1.4 N.A lens on a Zeiss LSM 800 confocal microscope (Carl Zeiss). Zeiss Zen Blue was used to acquire microscope images of 1024x1024 pixels at 16-bit depth with line averaging 2. Zeiss Zen was used to measure the intensity of RFP in cells along an edge of a germarium in a line starting adjacent to Cap cells and ending at the border of Fas3 expression. Measurements were taken in 5 µm distance increments within a z-stack range that did not exceed 8 µm. If there was more than one cell within the same 5 µm increment, both cells would be measured. If there was no suitable cell within the 8 µm z-stack range at a specific 5 µm increment, no measurement was collected. Fz3-RFP intensities were measured by circling the nucleus of a cell in the DAPI channel using the spline contour tool and obtaining the average intensity value in the 561nm channel for the enclosed area from the Z-stack with the highest RFP intensity. The 561nm laser intensity was set at 1.1% for 21-36h APF germaria and reduced to 0.3% beginning at 48h-APF through adult in order to keep RFP within the linear range. Because overall RFP levels varied from germarium to germarium, individual RFP profiles were constructed for each germarium; RFP intensities for each germarium were normalized to the brightest cell on the edge of that gemarium. Normalized intensities were then averaged for each time point. In order to reflect the relative RFP brightness from 27h to 36h-APF, the averaged, normalized values were divided by the laser intensity at which they were taken. The averaged normalized values for every distance along the 27h-APF and 36-APF curves were divided by 1.1. The averaged normalized values for every distance along the 48h-APF, 60h-APF, and adult-stage curves were divided by 0.3. The 27h-APF, 36h-APF, 48h-APF, 60h-APF, and adult were then normalized to the highest value in all of the graphs to reflect relative intensities and display the gradient profiles.

Because the 21h-36h germaria were necessarily acquired with a higher laser intensity, output levels in Photoshop were reduced from 255 to 100 to indicate the greatly reduced RFP intensity of those samples. The distances from Cap cells to the border of strong Fas3 expression on the edge of the germarium and to the posterior end of the germline in the center of the germarium were also measured for every germarium and averaged for each time-point.

## Results

### Cell lineage methods and analysis

The cell-autonomous effects of genetic alterations on cell fate can generally be studied by comparing outcomes for marked cells, with or without a genetic alteration, over the relevant developmental period. Here, we are interested in examining all somatic cells of an ovariole from pupariation (when mature third instar larvae cease movements) to adulthood. We therefore induced FRT-catalyzed recombination events to mark and modify any ovarian somatic cell at a chosen developmental stage by using a heat-shock inducible Flp recombinase transgene. Prior studies using this approach for wild-type cells suggested that the number and identity of adult somatic cells in a marked lineage depends principally on the A/P location of the marked precursor at pupariation (Reilein *et al*. 2021). Anterior precursors become ECs, posterior precursors become FCs, and precursors in intermediate locations produce mixed lineages that include ECs, FSCs and FCs (Fig. 1A). Moreover, posterior precursors divide faster and, unlike EC-only precursors, they do not cease division during pupation (Reilein *et al*. 2021). Here, we use equivalent lineage studies with specific genetic alterations to identify factors that guide patterned cell division, differentiation of FCs during pupation, and anterior-posterior (A/P) migration of somatic ovarian precursors, which dictates EC and FSC fates in the adult germarium. There are, however, some significant technical challenges and limitations.

First, the marked cells in a single adult ovariole ideally derive from a single precursor within a pool of about two dozen precursors at pupariation. We therefore used a very mild heat-shock protocol, carefully administered for uniformity, to induce mitotic recombination in a MARCM (Mosaic Analysis with a Repressible Marker) strategy (Lee and Luo 2001). In practice, this generally leads to about 30-50% of all ovarioles with at least one labeled cell (Fig. 1B). Some of these ovarioles will, however, include derivatives of two or more initially marked cells. We therefore estimated this fraction from the percentage of unlabeled ovarioles by assuming that all recombination events are independent, and we then inferred the frequency and composition of single-cell lineages (see Methods). An example of the results of this conversion of raw data to deduced single-cell lineage data is shown for a control sample (Fig. 1C). In all other presented data, only the deduced single-cell lineage data are shown (with sample numbers in figure legends referring to raw data; all raw data and conversion details are in supplementary data spreadsheets).

Second, although the chosen approach can simultaneously report on the behavior of all precursors in a variety of AP locations, it necessarily includes a sampling error. For example, if we are examining the fate of the roughly 20% of precursors that would normally become FCs, the sample of such cells will not always be exactly 20%. We have performed many sets of experiments, each including a control (“wild-type”) genotype. Those multiple controls allow experimental evaluation of sampling variability. They also provide a cumulatively large control sample to produce reliable measures for each parameter deduced from lineage studies. For MARCM clones induced at pupariation, the standard deviation among controls for the frequency of a given lineage type (Fig. 1B) was generally similar to that expected for random sampling of individual cells from the estimated number of somatic precursors present at pupariation (see Methods and cited supplementary spreadsheet).

Third, clone sizes provide only limited insight into the spatial regulation of division rates. First, the deduced average number of marked ECs per EC-only clone was consistently between 2 and 3 for controls, consistent with limited division of the most anterior precursors and providing little opportunity to measure significant quantitative deficits (Fig. 1D). Second, almost all lineages with FSCs also include ECs and FCs. The number of ECs and FSCs in such lineages (average of about 8 for controls) (Fig. 1D’) provides some indication of the amplification of a subset of the precursors, but this excludes the large and variable numbers of FCs generally present in such lineages. It is not possible to measure the number of marked FCs that first associate with a germline cyst (which would be the key number to measure divisions in and around the prospective FSC domain) because each such cell generally divides many further times as the egg chamber forms and grows. Thus, the sum of ECs and FSCs in an FSC-containing lineage only provides limited information about division rates in regions that give rise to ECs, FSCs and FCs. The fraction of cysts with marked FCs in FSC-containing lineages provides a second measure. The number of marked cells in the most posterior egg chamber can report on proliferation of the most posterior precursors. Additional information comes from measuring the frequency with which such lineages extend into the penultimate egg chamber.

Fourth, the MARCM method has the key virtue of clear positive marking, so that every lineage (even a single cell) and every marked cell within a lineage can be identified readily. A well-known disadvantage is, however, that GAL80 gene products must first be diluted and degraded before GAL4- driven *UAS-X* transgene products can accumulate. Strong accumulation of GFP from *UAS-GFP* in somatic ovarian cells takes at least 3d using our genetic reagents, and a similar delay is therefore expected before any *UAS-*driven transgene begins to exert maximal effect. Thus, both the onset and plateau of phenotypic responses to activation of a *UAS*-driven transgene in a MARCM clone can only be estimated and will generally be later than for the generation of a loss-of-function homozygous mutation. These and other relevant methodological considerations are discussed at greater length in the context of specific results obtained for variant genotypes.

### Terminal FC source is not distinct from other EC, FSC and FC precursors

We are interested in understanding how somatic cells are instructed to occupy various positions in the adult germarium to function subsequently as FSCs and ECs. We are also especially interested in how the first FCs are specified during pupation. This is an intriguing question because FC formation in adults relies on signals from previously produced FCs. We therefore first sought to clarify the origins of the FCs on the first-formed egg chamber.

At the start of pupation, about 20-30 somatic cells are dispersed among germline cells to form a primitive germarium (Fig. 2A). These intermingled cells (ICs) express the transcription factor, Traffic Jam (TJ) (Li *et al*. 2003). Numerous additional TJ-negative somatic cells, referred to as basal cells (Swarm cells after migration; not shown in Fig. 2A and not the same as basal stalk cells), are present in a more posterior location, as a seemingly contiguous group, rather than associated with individual developing ovarioles. Over the next 48h, TJ-positive somatic cells accumulate posterior to the developing germarium (beyond the most posterior germline cyst) to form a structure termed the Extra-Germarial Crown (EGC) (Reilein *et al*. 2021). In the most posterior portion of the EGC, somatic cells are intercalated to form a “basal stalk”, which also includes some TJ-negative cells at the posterior (King *et al*. 1968; Godt and Laski 1995; Vlachos *et al*. 2015; Reilein *et al*. 2021). Thereafter (at about 56h APF on average), the most posterior germline cyst moves away from the germarium, into the EGC and basal stalk, as cells in that extra-germarial territory, including some originally TJ-negative cells, move anteriorly along the cyst to form a monolayer epithelium, leaving a short residue of basal stalk at the posterior end that persists until eclosion (Reilein *et al*. 2021). Most of the mid-pupal (around 48h APF) EGC and basal stalk cells become FCs (or adult basal stalk cells) on the first budded (terminal) egg chamber but some contribute to the next (penultimate) egg chamber, alongside FCs derived from IC precursors present within the germarium around 48h APF (Fig. 2A). These deductions derive from fixed images at successive stages, live imaging cell tracing and lineage analyses initiated by a heat-shock at 36h APF (Reilein *et al*. 2021). Many such 36h APF lineages contained only marked FCs or basal stalk cells on the terminal egg chamber (purple lineage in Fig. 2A), while other lineages had marked FCs only in more anterior locations (cyan and yellow lineages in Fig. 2A). Each type of FC-only lineage sometimes included marked FCs in the penultimate egg chamber. Thus, at 36-48h APF, only precursors in the EGC or basal stalk contribute terminal FCs and only IC precursors yield FCs anterior to the two terminal egg chambers.

**Figure 2:**
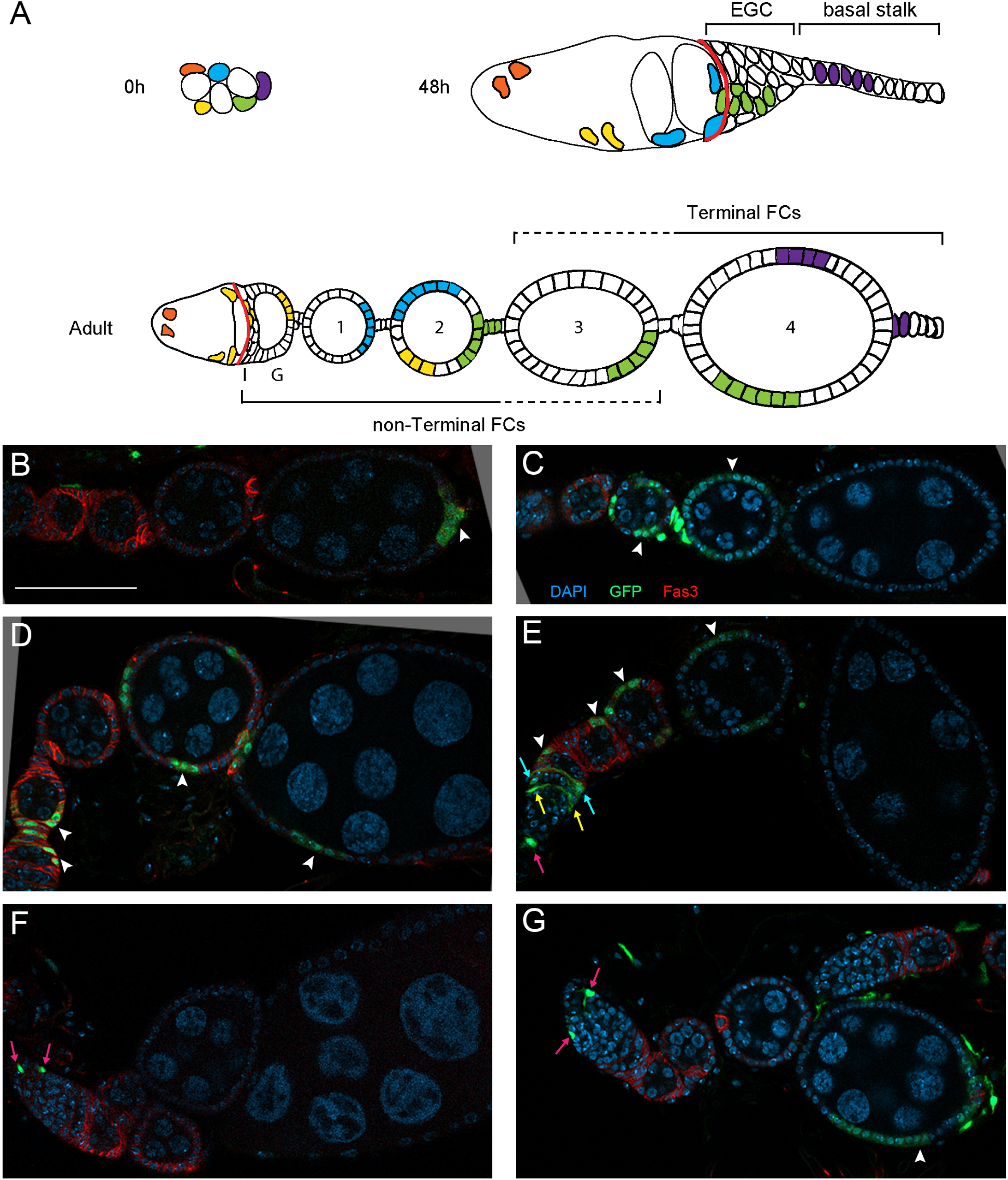
Terminal and non-terminal Follicle Cells derive from germarial ICs at pupariation. **(A)** Illustration of selected individual precursors at pupariation (0h APF) and their progression over time. At 0h APF, the somatic cells are interspersed with single germline cells and 2-cell cysts. By 48h APF, the most mature germline cysts have 16 cells and a single such cyst occupies the most posterior position in the germarium. At this time, more posterior somatic cells express Fas3 (anterior border of expression indicated by the red line). The relationships between precursors at 48h APF and adult derivatives have previously been established. Specifically, precursors within the Extra-Germarial Crown (EGC) and Basal Stalk, which contain no germline cells and are therefore considered extra-germarial, give rise to FCs in the terminal egg chamber or adult basal stalk (with some derivatives extending into the penultimate egg chamber but no further anterior). Precursors within the germarium at 48h APF give rise to adult ECs only (orange), ECs, FSCs and non-terminal FCs (yellow) or non-terminal FCs only (blue). It was not known whether cells in the EGC and basal stalk migrated out of the germarium between 0 and 48h APF (as indicated by the colors in the diagram) or if they derived from precursors in another location. The existence of precursors at pupariation producing adult FCs in terminal and non-terminal locations (as indicated for green cells) would support a germarial origin of terminal FCs. It would still be expected that some precursors at 0h APF would give rise only to terminal FCs (as shown for purple cells). The outcomes shown for green and purple cells were found, as illustrated in (b) and (D). **(B-G)** Representative images of lineages induced at 0h APF in ovarioles of newly-eclosed flies. **(B)** Terminal FC-only; **(C)** non-Terminal FCs only (egg chambers 2 and 3); **(D)** non-Terminal and Terminal FCs, showing that all FCs derive from germarial cells at pupariation; **(E)** EC/FSC/FC (designated as EC/FSC in graphs), **(F)** EC-only, **(G)** EC plus Terminal FC (presumed to be two lineages in the same ovariole because lineages derived from a single cell generally occupy largely contiguous territory). Scale Bar 50 µm for all images. White arrowheads indicate marked Follicle Cells; magenta arrows, r1 ECs; yellow arrows, r2a ECs; cyan arrows, FSCs. DAPI, blue; GFP, green; Fas3, red.

The remaining question is whether the precursors of the first FCs (EGC and basal stalk cells) emerge from germarial ICs over the first 48h of pupation. This hypothetical migration out of the developing germarium has not been visualized directly. There is extensive division outside the germarium, so there may be only a modest ongoing exodus from the germarium that escaped detection during live imaging. An alternative possibility is that the EGC and basal stalk cells seen at 48h APF have a separate origin from IC cells, conceivably deriving from extra-germarial basal cells present at pupariation (King *et al*. 1968) and acquiring TJ expression over the following days. These possibilities can be distinguished by lineage analysis. An FC-only lineage (induced prior to 36h APF) that includes FCs in both the terminal egg chamber (“terminal FCs”) and at least one location anterior to the penultimate egg chamber (“non-terminal” FCs) would provide evidence of a common precursor of ICs and EGC/basal stalk cells present before 36h APF.

From 12 separate control experiments for lineages initiated at pupariation, the average fraction of FC-only lineages with terminal FCs (n=101) that also included marked FCs anterior to the penultimate egg chamber was (0.253/0.685) 0.37. This argues strongly for a single cell often acting as a precursor to both terminal and non-terminal FCs. Such precursors, like other non-terminal FC precursors, are therefore within or contiguous with the IC population, meaning that EGC and basal stalk cells derive from posterior migration of ICs out of a primitive germarium. Thus, it appears that some precursors in the germarium at pupariation produce only terminal FCs (Fig. 2B; purple cell in Fig. 2A), others produce only non-terminal FCs (Fig. 2C; blue cell in Fig. 2A), while several produce both terminal and non-terminal FCs (Fig. 2D; green cell in Fig. 2A). More anterior precursors produce EC/FSC/FC lineages (Fig. 2E; yellow cell in Fig. 2A) or only ECs (Fig. 2F; orange cell in Fig. 2A).

It is possible that our method for estimating single-cell lineage composition from raw observations underestimates the frequency of “double-clones” in a single ovariole (see Methods). To estimate the potential impact of an underestimation of this type we looked at the average frequency of lineages with terminal FCs that also include ECs (Fig. 2G). This incidence (0.039/(0.102 + 0.039)= 0.26) was previously considered mostly due to the presence of two lineages. To estimate the frequency of a second lineage including non-terminal FCs rather than ECs (which are much more frequently marked) due to “double-clones”, we multiply by the relative frequency of the latter two lineages ((0.154-0.064)/0.644) = 0.14). This gives an expected incidence of (0.26 x 0.14) = 0.036, which is far below the observed incidence of 0.37. In fact, further studies reported here suggest that some lineages containing only marked ECs and FCs are potential EC/FSC/FC lineages that have lost FSCs rather than indicating two superimposed lineages. Hence, the incidence of “double-clones” in our deduced single-cell lineages is likely even lower than estimated above. We can therefore be confident that ICs and extra-germarial somatic cells at 36-48h APF have a common origin at pupariation, with the terminal FC precursors migrating out of the nascent germarium. This means that we can use altered genotypes in lineage analyses to look for differences in the production of terminal FCs in order to identify factors that regulate posterior migration out of the developing germarium over the first 48h of pupation. The lineage studies provide evidence that the migration must occur prior to 48h APF but do not define when the migration occurs. There are some Traffic-Jam-positive cells posterior to all germ cells even by 12h APF (Reilein *et al*. 2021), so it is likely that some migration is early.

### Consequences of altered precursor division rate

In an adult ovariole, the division rate of a marked FSC over a period of several days cannot simply be assessed by counting the number of marked derivatives, principally because FC derivatives are initially highly proliferative, so that the number of FCs directly produced from marked FSCs cannot be counted. The analogous limitation applies to any pupal precursors that give rise to a mixture of FCs and other cell types during pupation. The number of divisions for EC-only lineages can be measured, but the average size of control EC-only clones is only 2.7 (Figs. 1D and 3I’), providing only a small range for detecting reduced division rates. The most posterior somatic precursors at pupariation contribute principally to FCs of the first-formed egg chamber (purple cell in Fig. 2A). The division frequency of a posterior precursor can therefore theoretically be estimated as the proportion of FCs in the terminal egg chamber that are marked (ignoring lineages that only mark polar cell and stalk cells, which cease division soon after specification).

**Figure 3:**
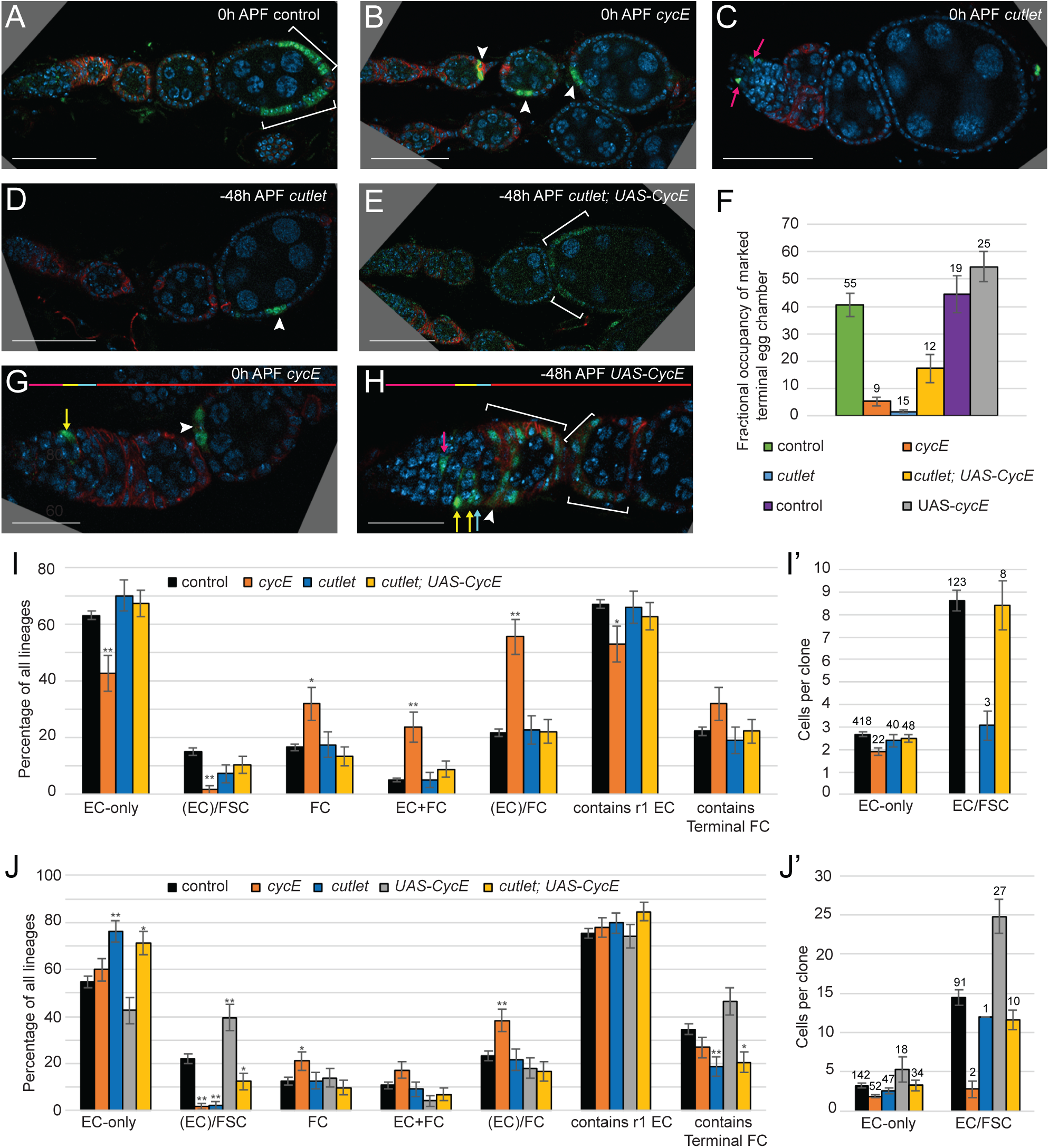
Effects of altered division frequency on lineage outcomes. **(A-F)** Fractional occupancy of terminal egg chamber by marked FCs, when present, as an approximate measure of division frequency for posterior precursors. **(A-C)** show lineages induced at pupariation and **(D-F)** are lineages induced 48h prior to pupariation for the indicated genotypes. **(F)** Percentage of marked cells out of total cells in terminal egg chamber for lineages of the indicated genotypes induced 48h before pupariation. Direct controls precede other genotypes for each experiment. **(G, H)** Ovarioles with **(G)** a *cycE* EC/FC lineage induced at 0h APF, illustrating the increased frequency of such lineages, potentially due to selective loss of FSCs from EC/FSC/FC lineages and **(H)** a *UAS-CycE* EC/FSC/FC lineage, present at elevated frequency, induced 48h before pupariation. White arrowheads indicate Follicle Cells; magenta arrows, r1 ECs; yellow arrows, r2a ECs; cyan arrows, FSCs. DAPI, blue; GFP, green; Fas3, red. Scale Bar **(A-E)** 50 µm and **(G-H)** 20 µm). **(I, J)** Deduced frequency of the indicated types of single-cell lineage as a percentage of all lineages and **(I’, J’)** average number of cells per EC-only or EC/FSC lineage (ECs and FSCs only) induced at **(I)** 0h APF and **(J)** -48h APF. Error bars show SEM. One asterisk (*) indicates p<0.05; two asterisks (**) indicates p<0.005 (Fisher’s two-tailed exact test). **(I)** Number of marked ovarioles = 739 (control), 50 (*cycE*), 60 (*cutlet*), 78 (*cutlet; UAS-CycE*). **(J)** Number of marked ovarioles = 319 (control), 93 (*cycE*), 119 (*cutlet*), 85 (*UAS-CycE*), 75 (*cutlet; UAS-CycE*). **(I,’ J’)** Number of lineages indicated above bars. For raw data and calculations please see supplementary spreadsheets titled 0h APF Graphs Fig1,3; -2d APF Graphs; FC occupancy aggregate; Compilation of Numbers Figure 3 Division -5d; Compilation of Numbers Figure 3 Division -7d.

We wanted first to establish whether changes in division rate can be measured in pupal ovary precursors in various locations and then to determine whether altering the rate of cell division has a systematic effect on developmental outcomes. We have previously shown that a specific hypomorphic *cycE* allele (*cycE^WX^*) reduces the division rate of adult FSCs without affecting differentiation significantly, and that this leads to loss of marked FSCs over time (Wang and Kalderon 2009; Reilein *et al*. 2018). The reduced competitiveness of slowly dividing FSCs is an important characteristic that results from division and differentiation being independent processes for these stem cells (as in some other population asymmetry paradigms) (Vermeulen and Snippert 2014; Reilein *et al*. 2018). Essentially, this is because the marked *cycE* mutant stem cells drain from the stem cell domain at the normal rate through differentiation but are replenished by division at a lower rate.

We found that the average number of marked ECs per EC-only clone for *cycE* induced at 0h APF was 1.9, compared to 2.7 for controls (Fig. 3I’). This is consistent with reduced division of the most anterior somatic ovarian precursors but likely does not report the magnitude of the effect well because wild-type *cycE* products likely perdure for the short duration of pre-EC divisions. The most posterior somatic precursors at pupariation contribute principally to FCs of the first-formed egg chamber (purple cell in Fig. 2A). The division frequency of a posterior precursor can therefore theoretically be estimated as the proportion of FCs in the terminal egg chamber that are marked. The average fraction of marked FCs among terminal FCs with main-body FC labeling was 3.0% for *cycE*, compared to nearly 50% for the direct control (Fig. 3A, B). This measure might be distorted if the altered genotype affects the likelihood of precursors becoming stalk or polar cells, which proliferate less than other FCs. However, marked stalk or polar cells were no more prevalent for *cycE* (2/14 samples) than for controls (5/10 samples). Thus, it appears that reduced CycE activity associated with the *cycE^WX^* allele greatly impaired cell division rates for posterior ovarian precursors.

Reduced precursor division frequency might be anticipated to have different consequences on lineage outcomes depending on precursor location. First, we might hypothesize that slowly-dividing *cycE* cells in regions giving rise to future FSCs will be outcompeted by normal precursors, depleting marked FSCs, exactly as seen for adult-induced FSC lineages. The logic here is that since there is a certain A/P domain where precursors reside to become FSCs, the survival of a lineage in this domain requires cell division to balance posterior loss. When induced at pupariation and analyzed in newly-eclosed adult ovaries, there was a large reduction in the frequency of *cycE* clones that included one or more FSCs (2% vs 15% for controls; p<0.005) (Fig. 3I). Out of all marked ECs and FSCs, only 1% of *cycE* cells were FSCs vs 23% of control cells, confirming the substantial deficiency in FSC production. The frequency of ovarioles harboring marked cells of any type was not significantly altered (47% *cycE*, 42% in the direct, side-by-side control), suggesting that the *cycE* mutation does not impact cell survival and is therefore unlikely to be depleting FSCs through selective cell death. The results therefore support the hypothesis that severely reduced cell division substantially reduces the probability of a cell emerging from pupal development as an FSC. Selective loss of FSCs from prospective EC/FSC/FC lineages would be expected to result in an unusually high frequency of lineages containing only marked ECs and FCs. This was observed for *cycE* lineages (Fig. 3G, I; p<0.005).

A second hypothesis is that reduced division rate may result in reduced posterior movement of precursors. The somatic cell population grows and expands posteriorly over the first 48h of pupation to occupy a wider A/P territory prior to the budding of the first egg chamber. While it is more likely that posterior expansion is fueled by pressure from growth and division of germline cysts and somatic cells that lie anterior, it is possible that the division of a cell has some cell-autonomous contribution to its posterior movement. If such an effect were significant, we might expect that *cycE* mutant cells would less frequently contribute to FCs of the terminal egg chamber and would more frequently become ECs, especially region 1 ECs. Contrary to that hypothesis, the fraction of lineages including terminal FCs was higher (32% vs 22%; p= 0.09) and the fraction with r1 ECs was lower (53% vs 67%; p<0.05) for *cycE* lineages compared to controls (Fig. 3I). The frequency of EC-only lineages was also reduced for *cycE* (43% vs 63%; p<0.005), while the frequency of FC-only lineages was increased (32% vs 17%; p<0.05) (Fig. 3I). These results therefore suggest an unexpected cell- autonomous posterior movement of *cycE* mutant cells relative to surrounding precursors over the whole A/P range.

To test whether the posterior movement of *cycE* mutant precursors was due to reduced division or another action of CycE we tested an allele of the replication factor gene *cutlet*, which reduced division rate in adult FSCs (Wang *et al*. 2012). The mean number of marked ECs per EC-only was 2.4 compared to 2.7 for controls (Fig. 3I’), while the average fraction of marked FCs among terminal FCs was 7.4%, compared to 24.3% for the direct control (Fig. 3C). These results are consistent with reduced cell division rates for *cutlet* mutant cells, albeit to a lesser degree than observed for the *cycE^WX^* allele. The fraction of lineages with marked FSCs was 7%, compared to 15% for controls (Fig. 3I). EC-only clone frequency increased slightly relative to controls (70% vs 63%) but the frequencies of FC-only lineages (17% vs 17%), labeling of r1 ECs (66% vs 67%) and of terminal FCs (19% vs 22%) were largely unchanged (Fig. 3I). Thus, there was selective loss of *cutlet* mutant FSCs without any significant A/P bias of precursor fates. Loss of FSCs for both *cycE* (15% to 2%) and *cutlet* (15% to 7%) is consistent with the conclusion that reduced cell division rates lower FSC production. By contrast, a large deficit of r1 EC-only lineages, an increase in FC-only lineages far exceeding that expected just from FSC loss, and an increase in terminal FCs were observed only for *cycE,* suggesting that these measures of net posterior movement of precursors do not result from a reduced division rate.

We then assessed the effects of overexpression of CycE, using *UAS-CycE* expression with *tub- GAL4* plus *act-GAL4* as drivers of expression (our standard MARCM condition) in marked lineages induced 48h before pupariation. The timing was chosen so that increased CycE levels might start to accrue roughly coincident with pupariation. In adult FSC lineage studies, *UAS-CycE* increased the division rate of FSCs but not ECs, which are quiescent in adulthood (Melamed and Kalderon 2020; Melamed *et al*. 2023). EC precursors are still dividing early in pupal development, so they might be stimulated to divide more frequently if excess CycE is provided at the start of pupation. Excess CycE would certainly be expected to increase the division rate of more posterior somatic cell precursors. Hence, by the converse of the logic explained earlier for reduced CycE, faster division might increase the representation of marked FSCs. The number of marked cells per EC-only clone was higher for *UAS-CycE* (5.3) than for controls (3.2) (Fig. 3J’). The average fraction of marked FCs among terminal FCs was also increased, from 45% to 55% (Fig. 3C). Among EC/FSC/FC lineages (where FC production cannot be counted, limiting the accuracy of estimating division rates), the average number of marked germarial cells (ECs and FSCs) rose from 14.5 (including 9.5 ECs) for controls to 24.8 (including 17.4 ECs) for *UAS-CycE* (Fig. 3J’). Thus, *UAS-CycE* increased the division rate of precursors in all locations. *UAS-CycE* produced a slightly higher clone rate (frequency of ovarioles with marked cells) than the side-by-side control (65% vs. 56%), demonstrating that excess CycE does not reduce cell survival.

Importantly, *UAS-CycE* lineages contained FSCs (40%) more frequently than controls (23% for direct control; 22% for average controls; p<0.005) (Fig. 3H, J). These results support the hypothesis that increased division rate during pupal stages can increase FSC allocation, just as faster division increases FSC representation in adult FSC lineages.

The observed increased representation of EC/FSC/FC lineages is expected to be because potential FSCs are less frequently lost to yield lineages with just ECs and FCs. Indeed, the frequency of (EC+FC) lineages was lower for UAS-CycE (4% vs 11% controls for lineages induced 48h prior to pupariation) and higher for reduced *cycE* activity induced 48h prior to pupation (17% vs 11%) or at pupariation (24% vs 6%). These results suggest that a wild-type precursor in a location that can yield ECs, FSCs and FCs because of modest A/P spread during pupal development will occasionally contain only adult ECs and FCs because cells in prospective FSC locations were not replenished fast enough to counter continuous drain to more posterior FC locations. That, in turn, suggests that a significant fraction of (EC+FC) lineages result from single precursors, rather than the super-position of an EC-only and an FC-only lineage, as previously thought (Reilein *et al*. 2021) and that our estimation of single- cell lineages numbers derived from raw data is better than we had believed on the basis of the deduced (EC+FC) lineage frequency.

The frequencies of both EC-only (43% vs 55%; p=0.1) and r1-only lineages (24% vs 43%; p<0.005) were decreased for *UAS-CycE*. This could simply be because the increased number of cells per lineage inevitably results in several clones crossing posterior boundaries, into r2a EC and FSC territory, respectively. The frequency of FC-only lineages was unaltered (14% vs 13%), suggesting there is no large systematic displacement of all marked cells towards the posterior. Rather, the principal response was that the representation of a central population, which become FSCs, was increased.

Since *UAS-CycE* was found to restore survival of adult FSC lineages with *cutlet* mutations (albeit assessed prior to more definitive measures of FSC competition introduced in 2017) (Wang *et al*. 2012), we tested whether excess CycE might ameliorate the key phenotypes of *cutlet* mutant lineages induced 48h prior to pupariation (deficiency of FSCs and indications of reduced division rate). *UAS-CycE* increased the average number of number of cells in EC-only lineages from 2.5 to the control value of 3.2 (Fig. 3J’). The average fraction of marked FCs among terminal FCs was greatly increased, from 2% to 17%, indicating substantial rescue of division rate towards control values (Fig. 3C). The frequency of lineages with FSCs was also partially restored (from 2% to 12%) towards the control value (22%) (Fig. 3D, E, J). In the EC/FSC lineages there were now similar numbers of ECs (8 vs 9.5) and FSCs (3.5 vs 5.0) to controls (Fig. 3J’, sum shown). The FC productivity of FSC-containing lineages can be roughly estimated by counting the proportion of egg chambers, region 2b and region 3a germline cysts with associated marked FCs; this measure would be increased by recruitment of more future FSCs or a higher division rate of those cells. The low proportion of cysts with marked FCs for *cutle*t (1/11; 9%) was increased by *UAS-CycE* (33/54; 61%) towards the control value for this experiment (128/156; 82%). These results are all consistent with substantial correlated restoration of cell division rate and FSC production. A smaller restoration of FSC-containing lineages (from 7% to 10%) towards control values (15%) was seen for the addition of *UAS-CycE* to *cutlet* loss of function in lineages initiated at pupariation (Fig. 3I). The proportion of cysts with surrounding marked FCs in FSC-containing lineages was also increased from 14% (3/21) for *cutlet* to 46% (25/54) by the addition of *UAS-CycE*, towards the control value of 65% (26/40). The delayed expression of excess CycE in lineages initiated at pupariation likely accounts for the lesser rescue.

Reduced CycE activity was also tested in lineages induced 2d prior to pupariation (Fig. 3J). The major phenotype was a large loss of lineages with FSCs, similar to *cutlet*. There was also an increase in FC-only and EC/FC lineages combined (“(EC)/FC”) for *cycE* (38%) relative to *cutlet* (22%) and controls (23%), similar to results for lineages induced at pupariation. However, there was neither an increase for terminal FC representation (27% vs control 35%), nor a large reduction in r1 EC representation (80% vs 75%) seen for lineages initiated at pupariation. Thus, the strong phenotype of FSC loss was seen for both stages of lineage induction, while the unexpected posterior movement of precursors due to loss of *cycE* was prominent only for lineages induced at pupariation.

This set of studies suggests that the primary consequence of cell-autonomously reducing the rate of precursor division within a competitive environment of normal cells during pupation is a large reduction in the frequency of lineages containing FSCs in newly-eclosed adults. This was observed for both loss of *cycE* and loss of *cutlet*, initiating lineages at pupariation and 2d earlier. The converse was seen in response to increasing the division rate. No other consequence was consistently seen for both *cycE* and *cutlet* loss of function mutations, while excess CycE also appeared only to affect FSC representation.

### Wnt signaling magnitude strongly influences precursor fates across the whole A/P domain

We wished to investigate the roles of Wnt and JAK-STAT signaling on somatic ovarian precursor development because these are the two major pathways regulating adult FSC behavior (Vied *et al*. 2012; Reilein *et al*. 2017; Melamed and Kalderon 2020). Canonical Wnt signaling is initiated by binding of ligand to Frizzled and LRP5/6 (Arrow in Drosophila) family co-receptors, leading to release of β-catenin from a destruction complex scaffolded by Axin (Axn) and its subsequent association with nuclear TCF/LEF (TCF in Drosophila) to convert TCF from a repressor to an activator of transcription (Wiese *et al*. 2018).

We used the strong *axn^E77^* allele (which truncates the protein after Q406 to eliminate DIX, GSK3, β-catenin and protein phosphatase 2A binding domains to behave like a null in several assays) to create marked homozygous *axn* mutant precursors at pupariation and measure the spectrum of lineages obtained consequent to increasing Wnt pathway activity cell autonomously during pupal development. Almost all lineages consisted of only marked ECs (91%, compared to 63% in controls) (Fig. 4A, D, F), with an elevated yield of lineages confined to r1 ECs (61% vs 49% control) and of lineages containing at least one marked r1 EC (81% vs 67% control). Only 2% of lineages contained marked FCs without FSCs (22% for controls) and only 6% contained any marked FSCs (15% for controls) (Fig. 4F). Thus, it appears that all marked precursors have preferentially moved anteriorly during pupal development in a cell-autonomous response to increased Wnt pathway activity, leading to a near-absence of marked posterior derivatives, FSCs and FCs. In adult FSCs, loss of *axn* leads to strong net anterior movement, producing a high frequency of marked ECs over time but it also severely reduces cell division (Reilein *et al*. 2017; Melamed and Kalderon 2020). Based on *cycE* and *cutlet* mutant phenotypes over pupal development, reduced division would be expected to reduce the yield of FSCs but some of the “lost” FSCs would become FCs. The drastic loss of both FSCs and FCs for *axn* mutant precursors argues for a strongly altered behavior (anterior migration) other than reduced division. The average number of cells per EC clone was slightly lower for *axn* (2.5) than the control value (2.7). However, this is not a sensitive test and the *axn* mutant ECs, on average, also occupy a slightly more anterior set of EC positions than control derivatives, which may lead to earlier cessation of division. Since almost all *axn* precursors become ECs, there were too few FSC and FC *axn* derivatives to assess cell division rates of more posterior precursors. Thus, it is not clear whether increased Wnt signaling directly affects cell division during pupation but there is a strong phenotype of anterior bias for adult derivatives of precursors in all locations.

**Figure 4:**
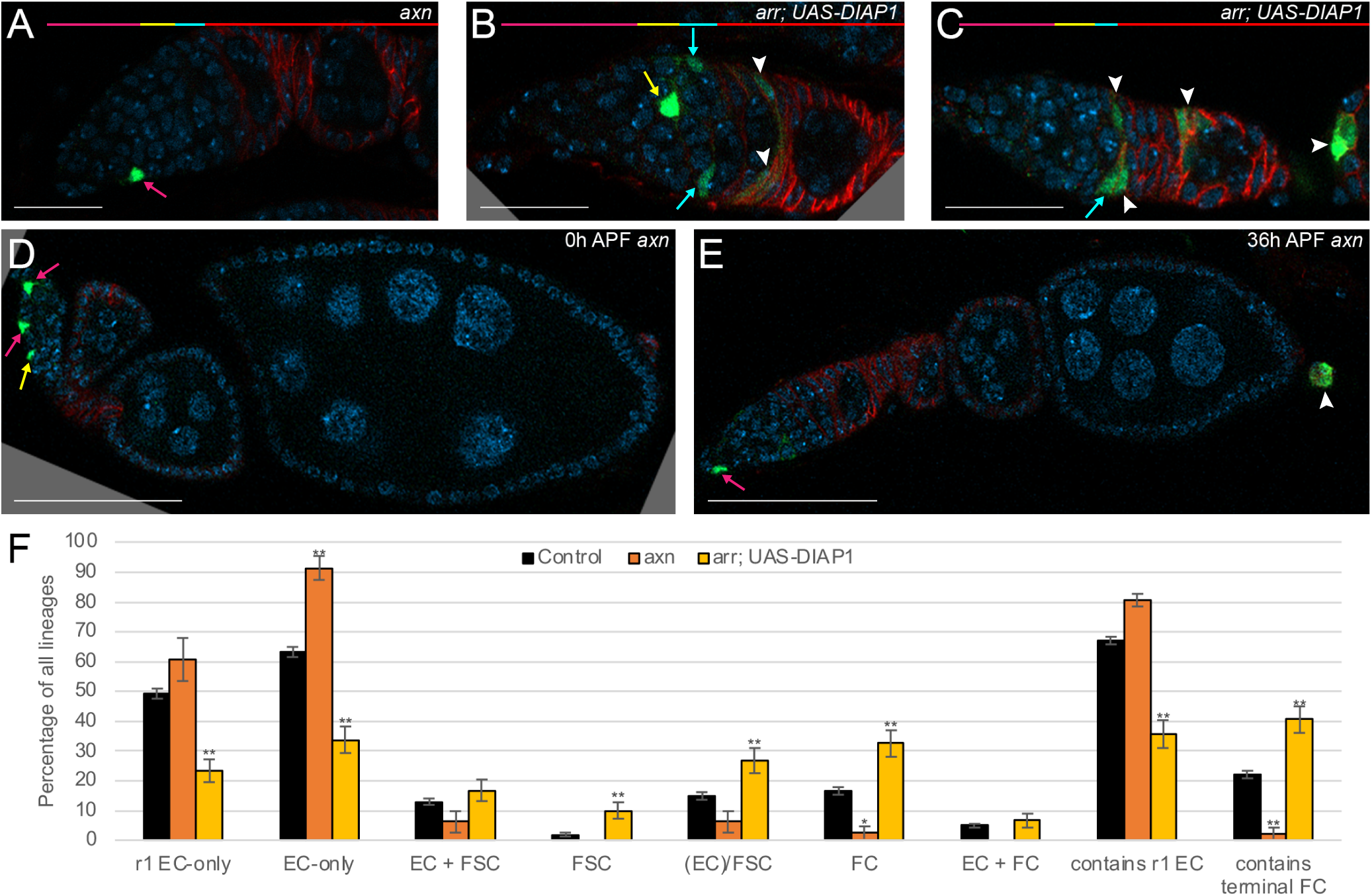
Higher Wnt pathway activity promotes anterior precursor movement during pupation. **(A-E)** Representative lineages induced at (**A-D**) 0h APF or (**E**) 36h APF for the indicated genotypes. White arrowheads indicate Follicle Cells; magenta arrows, r1 ECs; yellow arrows, r2a ECs; cyan arrows, FSCs. DAPI, blue; GFP, green; Fas3, red. (**A-C**) The r1 EC, r2a EC, FSC and FC domains are shown in corresponding colors in the horizontal bars above. Characteristic phenotypes with elevated frequency included **(A)** an *axn* r1 EC-only lineage and **(B, C)** *arr; UAS-DIAP1* lineages containing FSCs and FCs, (**B**) with or (**C**) without ECs (the latter is rarely observed for control lineages). **(D, E)** Ovarioles with an *axn* lineage induced at **(D)** 0h APF, with only ECs, and at **(E)** 36h APF, with marked terminal FCs in the basal stalk (arrowhead). Scale Bar 20 µm for (**A-C**) and 50 µm for (**D, E**). **(F)** Deduced frequency of the indicated types of single-cell lineage as a percentage of all lineages induced at 0h APF. Error bars show SEM. One asterisk (*) indicates p<0.05; two asterisks (**) indicates p<0.005 (Fisher’s two-tailed exact test). Number of marked ovarioles = 739 (control), 44 (*axn*), 95 (*arr; UAS-DIAP1*). For raw data and calculations please see supplementary spreadsheets titled 0h APF Graphs Fig4-10; Compilation of Numbers Figure 4 Wnt.

To investigate the effects of loss of Wnt pathway activity we used a strong or null *arrow* allele (*arr^2^*) that introduces a premature stop codon. We also added *UAS-DIAP1*, which was found in adult FSC derivatives to increase the yield of labeled *arr* mutant ECs (Melamed and Kalderon 2020). Other studies have found that adult ECs, especially those at the far anterior, are subject to apoptosis when Wnt pathway activity is severely reduced (Wang *et al*. 2015; WANG AND PAGE-MCCAW 2018). Relative to controls, *arr* mutant lineages induced at pupariation (with *UAS-DIAP1*) had a strong deficit of lineages with only r1 ECs (23% vs 49% control), only ECs (34% vs 63% control) or any r1 EC (36% vs 67% control) (Fig. 4B, C, F). By contrast, there was an increased fraction of lineages with marked FSCs (27% vs 15% control), only FCs (33% vs 17%) or containing terminal FCs (41% vs 22% control). The average number of marked germarial cells in each EC-only (2.1 vs 2.7) or EC/FSC/FC lineage (7.4 vs 8.6) was largely unchanged. Thus, as in adult FSC lineages, loss of Wnt pathway activity appears to have no major effect on the rate of cell division but causes a strong posterior shift in location, leading to a deficit of ECs and an excess of FCs. The increase in lineages with FSCs can be rationalized as due to a greater numerical shift of would-be ECs (which are the most numerous) to FSCs than of would-be FSCs to FCs. These conclusions explain the unexpected observation long ago that loss of *arr* increased FSC clonal yield if induced at larval stages and assayed 10 days later, despite reducing FSC longevity in adult FSC lineages because of a posterior shift increasing conversion of FSCs to FCs (Vied *et al*. 2012). In other words, the same primary consequence of preferential posterior movement due to autonomous loss of Wnt signaling reduces FSC yield in adults because marked FSCs cannot be replenished from marked ECs, but increases FSC yield during pupation because marked EC precursors can move into FSC locations; in both situations there is faster depletion of marked FSCs posteriorly to become FCs.

The effects of altered Wnt pathway activity on the conversion of precursors to adult ECs and FSCs can also be evaluated by counting the total number of these derivatives over all examined lineages (FC numbers cannot be evaluated in this way). This analysis has the virtue that it does not rely on whether marked cells in an ovariole derive from one or more precursors. In line with the analysis of lineage types above, the proportion of derivatives that became FSCs (control 20%) was greatly reduced for *axn* clones (6%) and markedly increased for *arr* clones (35%), while the proportion that became r1 ECs (control 54%) was similar for *axn* clones (50%) and markedly decreased for *arr* clones (31%). The unchanged proportion of r1 ECs for *axn* is somewhat deceptive because FCs are not included in this analysis. Control samples include many FCs whereas *axn* lineages do not. Hence, r1 EC frequency as a proportion of all marked cells (including FCs) is actually lower for controls than for *axn*.

### Wnt pathway activity limits cell migration out of the germarium during the first half of pupation

We have shown that EGC and basal stalk cells, the progenitors of terminal egg chamber FCs, derive from precursors that migrate posteriorly out of the germarium over the first 48h of pupation, and that increased Wnt signaling throughout pupation strongly reduces the acquisition of terminal FC identity cell-autonomously (Fig. 4F). To refine the key period of action, we induced control and *axn* lineages at 36h APF, estimating that genetically increased Wnt signaling would be delayed by a further few hours to coincide roughly with the completion of EGC/basal stalk formation and the migration of the most posterior germline cyst out of the germarium. At 36h APF there are many more somatic cell precursors per ovariole than at pupariation, so even the mildest heat-shock conditions produce MARCM clones in almost all ovarioles. Consequently, most ovarioles will contain more than one lineage and it is not possible to derive single-cell lineage frequencies accurately; raw observations are therefore presented for that time-point. The clone frequency for *axn* (97%) was higher than for the direct control (58%), so that multiple lineages per ovariole are likely more frequent. Nevertheless, there was clearly no deficit in the fraction of marked ovarioles with labeled terminal FCs (46% vs 31% control) (Fig. 4E; Supplementary spreadsheet “-3.5d axn August 2023 Numbers”). This is markedly different to the result for lineages initiated at pupariation, indicating that the near absence of terminal FCs for those samples is caused by failure of precursors to migrate posteriorly out of the germarium over the first 48h of pupation, with no subsequent deficit in surviving as terminal FCs.

The frequency of lineages that included terminal FCs was much higher for *arr* (41%) than for control lineages (22%) when induced at pupariation, indicating that Wnt signaling normally limits terminal FC formation (Fig. 4F). When lineages were induced 24h APF there was a similar clone frequency for control (84%) and *arr* plus *UAS-DIAP1* (73%) and a similar proportion of ovarioles that included marked terminal FCs (29% control vs 30%; Supplementary spreadsheet “-4d arr DIAP Numbers”). Thus, the increased representation of terminal FCs seen for lineages induced at pupariation is absent 24h later. We deduce that normal Wnt signaling restrains posterior movement of precursors out of the germarium at the start of pupation. These consequences of elevated and reduced Wnt signaling on generation of the first FCs parallel the effects seen in adults for conversion of FSCs to FCs, in which a precise level of Wnt signal is required for normal FSC production and optimal FSC maintenance, even though FC formation in adults and for the first egg chamber in pupae are morphologically very different.

### Wnt activity is influential throughout pupation

The above results mostly showed a strong effect of Wnt signaling favoring more anterior precursor locations when changes in Wnt pathway activity were applied throughout pupal development. Additionally, strong effects on terminal FC formation were seen for genetic changes initiated at pupariation but not if initiated at 24h APF (*arr*) or 36h APF (*axn*), showing that Wnt signaling levels are critical very early in pupariation for migration of precursors out of the germarium to form the first FCs. In theory, further insight into critical times of action can be gained by initiating lineage tracing and a change in Wnt signaling at a full range of developmental times. However, a limitation of using MARCM clones is that a strong GFP signal only develops after 3d because of perdurance of GAL80 products. We therefore induced clones at 6d, 5d, 4d, 3d and 2d before eclosion and examined lineages in adults two days after eclosion. A drawback of this approach is that the first egg chambers produced during pupal development have matured into deposited eggs and are no longer present in 2d-old adults, so marked FCs cannot be scored comprehensively. Hence, our analysis focused only on ECs and FSCs. Also, FSCs present at eclosion are not scored directly but must be inferred (see Methods). The experiment was performed for *axn* and also for loss of a different β-catenin destruction complex member (using a double mutation, *apc1 apc2*) with similar results, for *arr*, and for three control genotypes. Results, expressed as the percentage of all marked ECs/FSCs (Supplementary Fig. 1), and were summed for all controls, two *arr* tests and the two genotypes (*axn* and *apc1 apc2*) producing excess Wnt pathway activity. A small deficit of FSCs was apparent for *axn*/*apc* in lineages initiated 2d before eclosion (48% vs 55% control) and a larger deficit was seen at 5d and 6d prior to eclosion (14% vs 33% control in each case), suggesting that increased Wnt pathway activity provokes movement of precursors away from FSC-producing AP locations throughout pupal development (Supplementary Fig. 1). Loss of *arr* activity produced a strong r1 EC deficit (5% vs 16% control) and a substantial FSC surfeit (84% vs 55%) even from 2d prior to eclosion, showing that normal Wnt signaling influences A/P movement of precursors in the last two days of pupal development. The excess frequency of FSCs among ECs and FSCs for *arr* relative to controls averaged over -2d and -3d clone initiation times was 32%; it was 64% averaged over -6d and -5d initiation points. (Supplementary Fig. 1 and supporting information spreadsheet). These data suggest that loss of Wnt activity, and hence normal Wnt signaling, has an effect on cell location within EC/FSC precursor territory, as well as on terminal FC precursors, early in pupal development. The above deductions have the limitation that they are based on FSC or r1 EC representation among non-FC precursors rather than all precursors. Both *axn* and *arr* alter the proportion of precursors that become FCs and the magnitude of this effect may vary during pupation, contributing to changes in the measured representation of FSCs and r1 ECs over time.

### Pattern of Wnt pathway activity during pupal ovary development; an A/P gradient

Wnt pathway activity in adult germaria, measured with a Fz3-RFP reporter that responds to genetic pathway manipulations in that tissue (Wang and Page-mccaw 2014; Dai *et al*. 2017; Reilein *et al*. 2017), is uniformly high in ECs and declines over the FSC domain to zero in the earliest FCs (Fig. 5L). The sources of Wnt in the adult germarium include high levels of Wg and Wnt6 production in Cap cells and lower levels of Wnt 2 and Wnt4 in ECs (Sahai-Hernandez and Nystul 2013; Wang and Page-mccaw 2014; Wang and Page-mccaw 2018).

**Figure 5:**
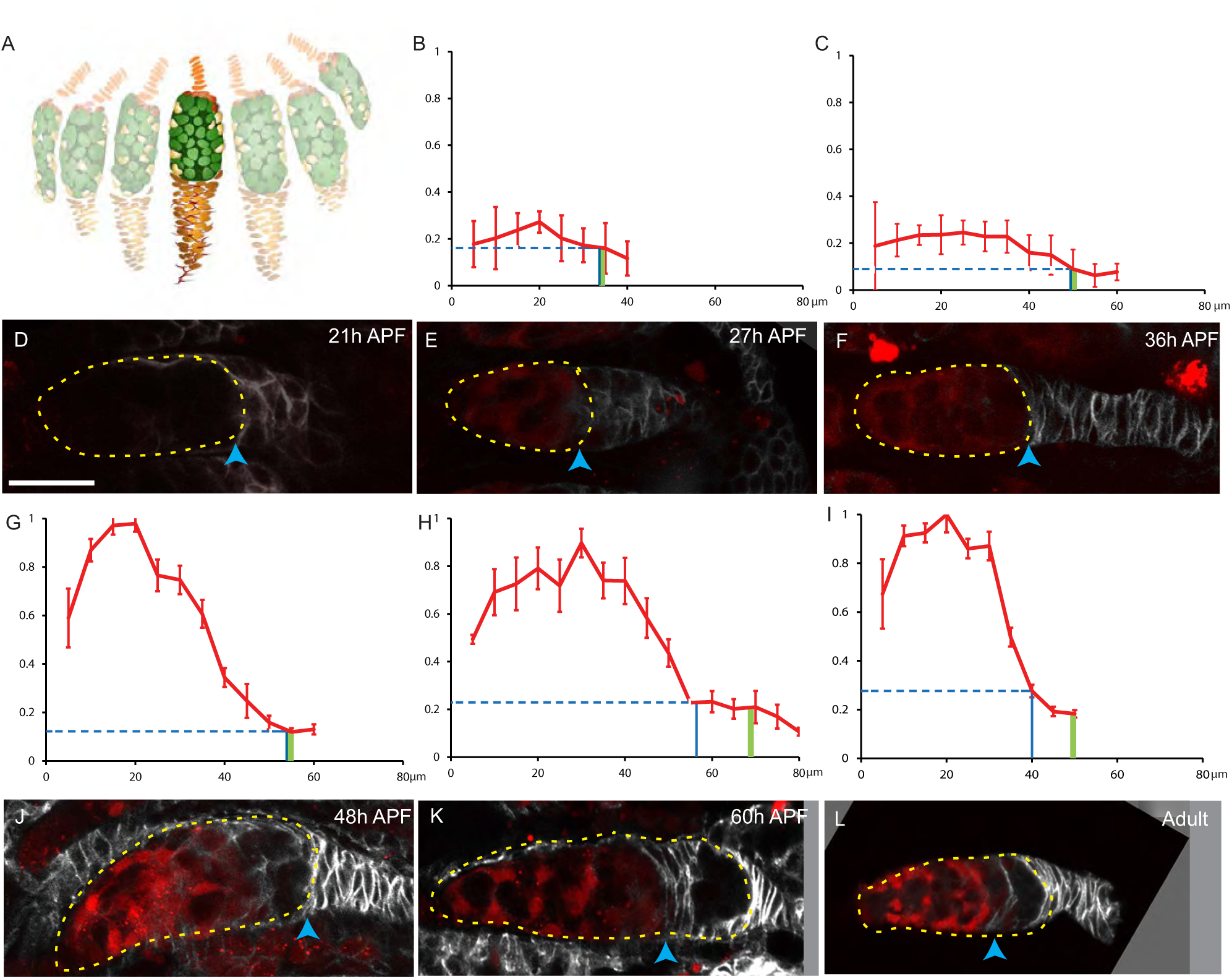
Development of a Wnt signaling gradient in pupal germaria. **(A-L)** Results from pupae expressing Fz3-RFP as a reporter for Wnt pathway activity. **(A)** Diagram of a 30h APF ovary. An individual germarium is highlighted with anterior at the top. **(D-F, J-L)**. Individual germaria at the noted stages, oriented anterior to the left, with the germarial region (containing germline cells) outlined in yellow and the anterior border of strong Fas3 staining indicated by a blue arrowhead. Samples were stained for Fas3 (white) and pupal samples were imaged intact, without separating developing ovarioles. **(B, C, G-I)** RFP profiles were constructed by measuring RFP intensities (y-axes) at 5 µm intervals along the edge of the germarium where most somatic cell bodies are located, starting adjacent to Cap Cells (0 on x-axes) and moving posteriorly. Individual germaria varied in RFP brightness and therefore the profile for each germarium was normalized to the brightest cell in that germarium. Normalized curves were averaged and SEM calculated to show the average Fz3-RFP profiles derived from germaria at **(B)** 27h APF (n=50 cells in 7 germaria), **(C)** 36h APF (n=108 cells in 14 germaria), **(G)** 48h APF (n=138 cells in 15 germaria), **(H)** 60h APF (n=178 cells in 14 germaria) and **(I)** adults (n=178 cells in 14 germaria). The average location of the border of strong Fas3 expression (blue vertical line), Fz3-RFP levels at that location (horizontal dashed line) and the location of the average posterior end of the germline cells in the germarium (green vertical line; measured from the Cap Cells to the central point at the posterior of the germline) are indicated. **(D-F, J-L)** Multiple z-sections were combined in a maximum intensity projection to show Fz3-RFP in cells along the length of germaria. **(D)** Fz3-RFP was not detected at 21h APF, **(E)** was present at low levels with an anterior bias at 27h (4 µm-thick z projection) and **(F)** at 36h APF (5 µm-thick z projection) before **(J)** accumulating to high anterior levels with a steep decline ending close to the strong Fas3 border at 48h APF (4 µm-thick z projection) and **(K)** 60h-APF (5 µm-thick z projection), similar to **(L)** the profile in newly-eclosed adults (10 µm-thick z projection). Scale bar of 20 µm applies to all images. For raw data please see supplementary spreadsheet titled Fig5 Fz3RFP Graph.

In pupal germaria, Fz3-RFP was first detected 27h APF and showed a marked increase in intensity between 36h and 48h (Fig. 5A-G, J). At all stages, Fz3-RFP was higher in anterior than posterior regions, consistent with the possibility that Cap Cells, which are specified at the anterior of the germarium very close to the start of pupation, and even developing EC precursors, may be major sources of Wnt ligand, as in the adult. The delay in reaching high levels of pathway activity might be due to maturation of Cap Cell properties but is perhaps more likely due to functional differentiation of ECs from precursors, most of which cease division, starting from the anterior, by 48h of pupation (Reilein *et al*. 2021). At 48h APF the spatial decline in Fz3-RFP was spread over the posterior two- thirds of the germarium (Fig. 5G, J). By 60h APF, the domain of consistently high Fz3-RFP expression occupied a larger anterior territory, extending roughly halfway to the region of strong Fas3 expression, followed by a steeper decline to background levels (Fig. 5H, K). At all stages, the anterior border of strong Fas3 expression coincided with very low levels of Wnt pathway reporter activity. In adult germaria and in pupal ovarioles prior to release of the first germline cyst, cells at, and posterior to, the strong Fas3 border have committed to becoming FCs. The persistent coincidence of this Fas3 border with a decline of Wnt pathway activity to near zero suggests that low Wnt signaling may be a pre-requisite for FC formation at all stages. This hypothesis is supported by the near-complete block of FC formation during pupal development in *axn* mutant precursors and the increased propensity of *arr* mutant precursors to form FCs throughout pupal development. These responses to altered signaling appear to be mediated by net A/P migration throughout the precursor domain. Thus, *arr* mutant cells are not depleted immediately anterior to potential FC-producing zones because there is also enhanced migration to those positions from the EC precursor domain; in fact, adult FSC production from *arr* precursor cells was increased overall. Conversely, *axn* mutant cells that do not become FCs do not accumulate extensively in the neighborhood of FSCs; there was, instead, a net loss of FSC fates in favor of EC production.

### JAK-STAT activity promotes FC formation and favors more posterior ovarian precursor fates

In Drosophila, Unpaired ligands activate a receptor to promote cross-phosphorylation and activation of the receptor-associated Janus kinase (JAK), Hopscotch (Hop), followed by phosphorylation and activation of the STAT transcription factor (Zeidler and Bausek 2013). The effect of altered JAK-STAT signaling was explored using a strong loss of function *stat* mutation and overexpression of Hop, as used in studies of adult FSC behavior (Vied *et al*. 2012; Melamed and Kalderon 2020). The spectrum of adult derivatives of *stat* mutant precursors induced at pupariation was very similar to that of *axn* mutant cells. There was a drastic loss of lineages with any marked terminal FCs (2% vs 22%), of FC-only lineages (4% vs 17%) and of lineages with FSCs (5% vs 15%), while EC-only lineages (90% vs 63%) increased greatly in frequency (Fig. 6A, B, G). However, in contrast to the behavior of *axn* mutant cells, the frequency of lineages containing only r1 ECs (46% vs 49% control) or at least one r1 EC (64% vs 67% control) was not increased for *stat* mutant cells (Fig. 6G). The frequency of labeled ovarioles was similar for *stat* mutant cells (43%) and the direct control (47%) in the same experiment, suggesting that marked cells had not been lost extensively. Thus, we deduce that all precursors posterior to EC-producing territory have preferentially moved anteriorly during pupal development in response to loss of JAK-STAT pathway activity, leading to a near-absence of posterior derivatives, FSCs and FCs.

**Figure 6:**
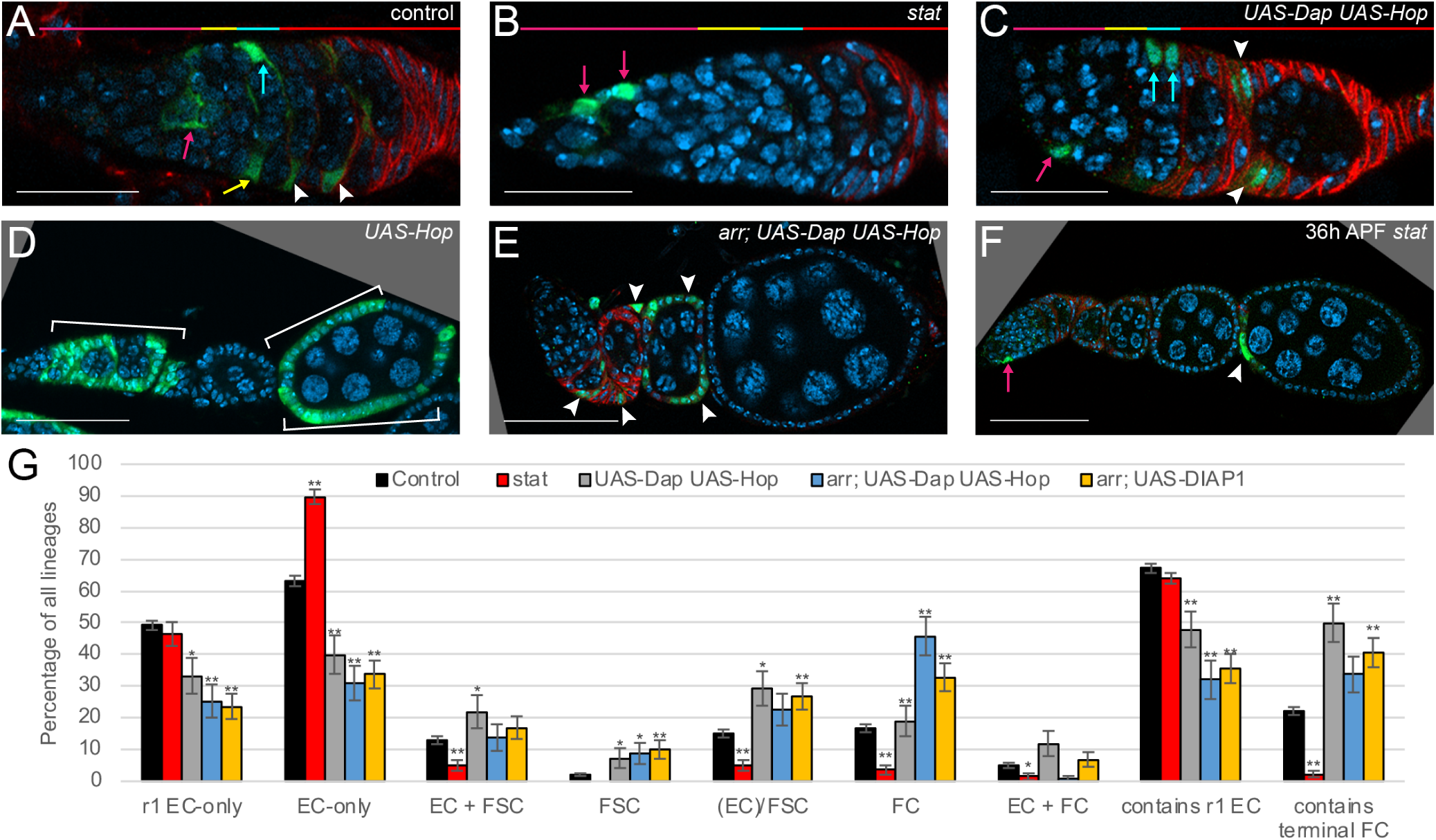
JAK-STAT pathway promotes posterior migration of precursors during pupation. **(A-F)** Representative lineages induced at (**A-E**) 0h APF or (**F**) 36h APF for the indicated genotypes. White arrowheads indicate Follicle Cells; magenta arrows, r1 ECs; yellow arrows, r2a ECs; cyan arrows, FSCs. DAPI, blue; GFP, green; Fas3, red. (**A-C**) The r1 EC, r2a EC, FSC and FC domains are shown in corresponding colors in the horizontal bars above. **(A)** control EC/FSC lineage that includes r1 ECs, **(B)** *stat* EC-only lineage, **(C)** *UAS-Dap UAS-Hop* EC/FSC/FC lineage, **(D)** *UAS-Hop* lineage, illustrating an exceptionally high number of marked cells, likely due to increased cell division, and **(E)** *arr; UAS-Dap UAS-Hop* lineage, illustrating the common FC-only lineage outcome. **(F)** A *stat* lineage induced at 36h APF illustrating terminal FC marking, which is virtually absent for *stat* lineages induced at 0h APF. Scale Bar 20 µm for **(A-C)** and 50 µm for **(D-F)**. **(G)** Deduced frequency of the indicated types of single-cell lineage as a percentage of all lineages induced at 0h APF. Error bars show SEM. One asterisk (*) indicates p<0.05; two asterisks (**) indicates p<0.005 (Fisher’s two-tailed exact test). Number of marked ovarioles = 739 (control), 142 (*stat*), 50 (*UAS-Dap UAS-Hop*), 57 (*arr; UAS-Dap UAS-Hop*), 95 (*arr; UAS-DIAP1*). For raw data and calculations please see supplementary spreadsheets titled 0h APF Graphs Fig4-10; Compilation of Numbers Figure 6 JAK-STAT.

We also examined *stat* mutant lineages induced at 36h APF. As for *axn*, marked ovarioles with labeled terminal FCs (38%) were now as frequent as for controls (31%) (Fig. 6F; Supplementary spreadsheet “-3.5d stat November 2021 Numbers**”**), consistent with a conclusion that JAK-STAT signaling is critical in the first 48h of pupation for promoting posterior migration of precursors out of the germarium into the EGC and basal stalk. Marked ovarioles with labeled FSCs were very rare (3% vs control 13%) and the fraction of all labeled ECs and FSCs that were FSCs was much reduced (3% vs control 21%), as for lineages induced at pupariation. This is consistent with the possibility that JAK-STAT signaling is required for FSC formation during the second half of pupation, perhaps by promoting cell division.

Expression of a *UAS-Hop* gene at 29C (or two copies at 25C) in adult FSCs leads to increased JAK-STAT activity and a variety of strong phenotypes, including increased FSC division and differentiation to FCs (Vied *et al*. 2012; Melamed and Kalderon 2020). In adults, the effects of increased JAK-STAT activity on cell location and differentiation could be studied in the absence of greatly altered cell division by co-expressing the CycE-Cdk2 inhibitor Dacapo (Dap) (Lane *et al*. 1996; Melamed and Kalderon 2020). Adult ovaries included very large numbers of labeled cells in *UAS-Hop* lineages induced at pupariation (Fig. 6D), indicating greatly increased cell division. The highly populated lineages included anterior, EC regions, which normally only include 2-4 marked cells. In adult ovaries, excess JAK-STAT can stimulate significant division in otherwise quiescent cells in the EC domain (Vied *et al*. 2012; Melamed and Kalderon 2020). The abundance of marked cells in pupal lineages precluded useful classification into clone types, other than noting that almost all lineages included ECs, FSCs and FCs.

We therefore added *UAS-Dap*. MARCM clones induced at pupariation to initiate *UAS-Hop* (and *UAS-Dap*) expression might be expected to increase JAK-STAT activity only after a delay. However, we found that the fraction of lineages that included terminal FCs was increased significantly (50% vs 22%, Fig. 6G), suggesting that increased JAK-STAT promotes terminal FC production and that the transgene markedly alters JAK-STAT activity within the first 48h after clone induction. Precursors with increased JAK-STAT activity also produced an increased proportion of lineages containing FSCs (29% vs 15%) (Fig. 6C), while greatly reducing the frequency of EC-only lineages (40% vs 63%), r1 only lineages (33% vs 49%) and lineages containing any r1 ECs (48% vs 68%) (Fig. 6G). The average number of cells in EC- only *UAS-Hop UAS-Dap* lineages was lower than for controls (1.1 vs 2.7). This is likely partly because many of the more posterior precursors that would normally produce only ECs shifted posteriorly to also produce FSCs (and FCs), so that EC-only lineages derived from only the most anterior precursors, which are the first to terminate division. The average number of ECs and FSCs in EC/FSC/FC lineages was similar to controls for *UAS-Hop UAS-Dap* lineages (8.0 vs 7.9 cells). Altogether, precursor division rates were likely similar to controls for this genotype, suggesting that the increase in FSC representation was due to direct effects on cell migration and not an increase in cell division rate, as observed for excess CycE expression.

Thus, increased JAK-STAT activity has the potential to promote posterior movement of all precursors, even from the most anterior r1 EC-producing locations. The restriction of the opposite phenotype for *stat* loss of function to cells posterior to the EC-producing domain (i.e., no increase in r1 EC-containing lineages) suggests that anterior regions of the developing ovariole may not normally experience significant levels of JAK-STAT signaling.

We examined JAK-STAT pathway activity using reporters containing either ten tandem STAT binding sites upstream of GFP (Bach *et al*. 2007) or STAT-responsive promoter sequences from the Socs36E gene upstream of RFP (He *et al*. 2019). Both reporters showed similar patterns in adult ovaries (Melamed and Kalderon 2020) and in pupal ovaries. There was very strong fluorescence surrounding each developing ovariole in both cases but this did not prevent signal detection in individual confocal sections of the developing ovarioles themselves (Fig. 7 and Supplementary Fig. 2). Activity was first detected clearly at about 30h APF, strongest around and anterior to the border between Fas3-positive and Fas3-negative somatic cells (Fig. 7A). Thereafter, reporter activity increased and spread posteriorly into Fas3-positive cells. The signal was lower in the anterior half of the developing germarium than in more posterior Fas3-negative somatic cells (Fig. 7B-D). These results are not sufficient to point to a specific cellular source of ligand. However, Fas3-positive cells are among the candidates. They may themselves initially be less responsive to ligand than Fas3- negative cells but reporter activity was as strong in Fas3-positive cells as in their anterior neighbors by 36h APF and thereafter (Fig. 7B-D; Supplementary Fig. 2). The higher pathway activity observed in somatic cells around, and posterior to, the three or four most mature germline cysts is consistent with the possibility that pathway activity in this region is necessary for a cell to maintain its position or to promote more posterior migration in order to seed FCs of the first budded egg chamber. We did not observe a clear gradient within this region (Fig. 7A-D).

**Figure 7:**
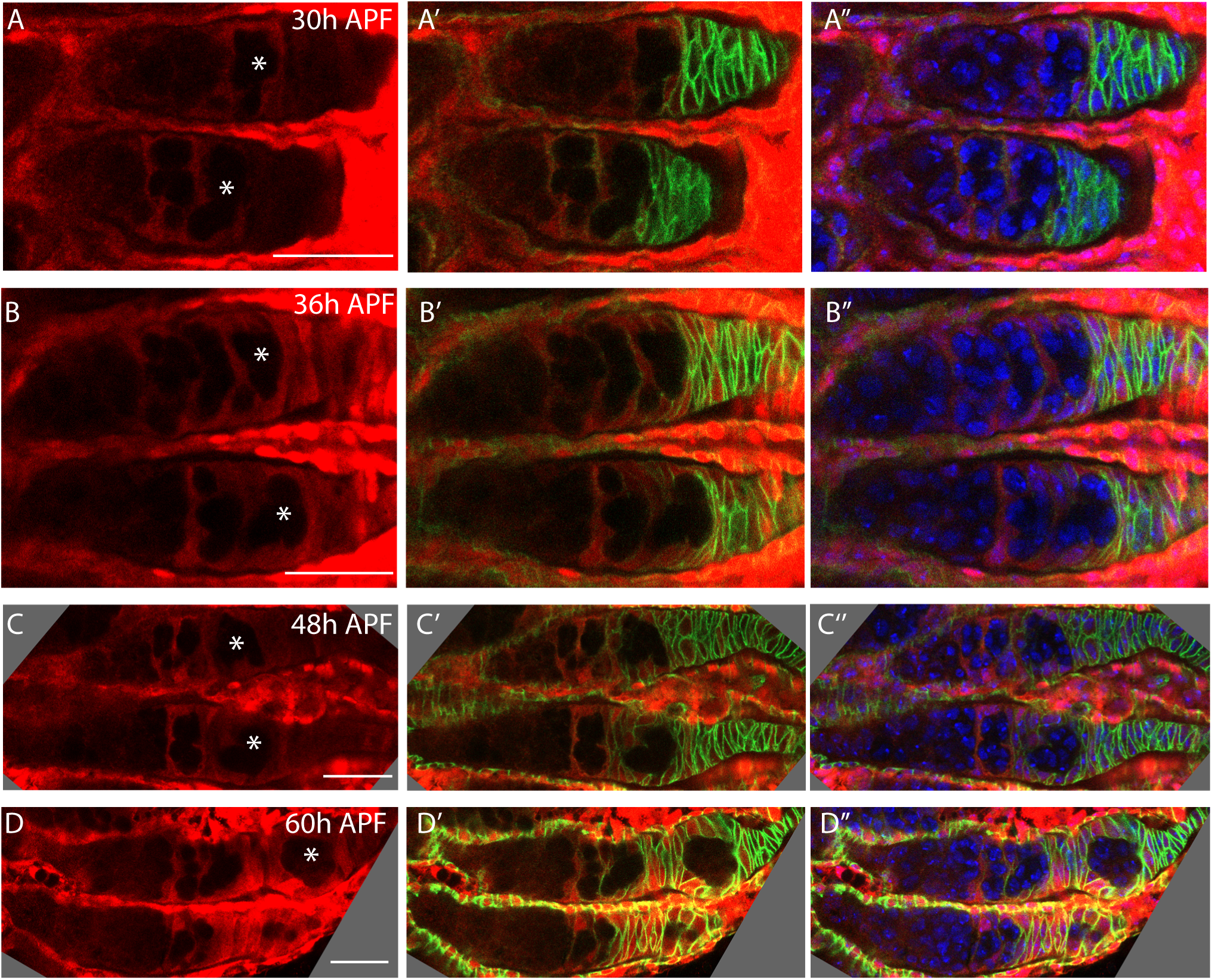
JAK/STAT signaling in pupal ovaries. **(A-D)** Pupal ovaries expressing STAT-RFP were stained for RFP (red), Fasciclin 3 (green), and with DAPI (blue). STAT-RFP was strongly expressed in epithelial sheath cells surrounding each developing germarium and the entire ovary. STAT-RFP was first detectable in developing germaria at 30h APF, and increased in intensity over time. **(A)** At 30h APF, STAT-RFP was detected at low levels around the most mature cysts in the germarium and in the most anterior Fasciclin III-expressing cells posterior to the germarium (in the Extra-Germarial Crown (EGC)). **(B-D)** From 36h APF to 60h APF, STAT-RFP expression was seen around the most mature egg chambers and throughout the EGCs and basal stalks. **(D)** At 60h APF, STAT-RFP was also seen in follicle cells of the budding egg chamber. The most mature cyst in each germarium is marked with an asterisk. Scale Bars, 20 µm.

Shortly after the most posterior germline cyst had left the germarium to form the first egg chamber (around 60h APF), somatic cells anterior to that cyst had particularly strong reporter activity, similar to nascent stalk cells during adult oogenesis (Supplementary Fig. 2). By that time, both Castor staining and higher Fas3 staining than in surrounding cells indicate the presence of polar cells in the first egg chamber (Reilein *et al*. 2021). Polar cells were similarly evident in the second egg chamber by about 84h APF, with strong STAT-GFP reporter activity in nearby somatic cells (Supplementary Fig. 2). The STAT-GFP signal extending into the germarium at 84h APF was significantly stronger than at earlier times and strongly resembled the adult pattern, with very little activity anterior to region 2a cysts (Supplementary Fig. 2). It is possible that the higher STAT-GFP signal is due to polar cells contributing JAK-STAT ligand. However, the overall spatial pattern of STAT-GFP was similar earlier and it is possible that the technical necessity of maintaining an intact ovary during dissection and staining of early pupal ovaries (rather than releasing individual ovarioles) limited GFP detection by antibody prior to 84h APF. Thus, there clearly is detectable JAK-STAT pathway activity prior to any polar cell formation in a pattern that largely excludes the anterior half of the developing germarium. It also appears that this pattern is significantly enhanced once polar cells have formed in the second budded egg chamber. The minimal levels of JAK-STAT pathway activity in anterior regions throughout the pupal period are consistent with the observation that marked r1 EC frequency is not altered by the absence of STAT activity, even though ectopic JAK-STAT can convert their precursors to adult cells in more posterior locations.

In adults, altering JAK-STAT activity in either direction greatly affects the division rate of FSCs, while excess JAK-STAT can promote significant division of otherwise quiescent ECs (Melamed and Kalderon 2020; Melamed *et al*. 2023). During pupation, loss of STAT activity barely altered the average number of cells in EC-only lineages (2.5 vs 2.7). This is likely partly because some of the precursors were originally in more posterior locations and set to divide faster. It is also consistent with the lack of net migration of r2a EC precursors to become r1 ECs and the observation of only low JAK-STAT activity in the normal EC precursor domain. There were insufficient *stat* mutant lineages in terminal egg chambers to measure average FC occupancy and insufficient FSC-containing lineages to measure the average number of marked ECs and FSCs. Thus, the strong anterior movement of *stat* mutant derivatives precluded direct measurement of the normal role of JAK-STAT signaling in supporting the division of posterior precursors through pupation.

Since loss of Wnt and increased JAK-STAT signaling have similar outcomes of promoting posterior precursor movement, it might be expected that combining these changes would produce still stronger phenotypes, as seen for adult FSCs. Adding *UAS-Hop UAS-Dap* to loss of *arr* produced very little change in the various measured parameters, other than an increase in FC-only lineages (46% vs 33%, Fig. 6E, G). Comparison of the phenotype of *arr UAS-Hop UAS-Dap* with *UAS-Hop UAS- Dap* is less straightforward for technical reasons: the clone frequency for the latter genotype was significantly higher, likely leading to incomplete resolution of double clones (suggested also by the comparatively high EC+FC frequency of 12% vs the exceptionally low value of 1% for the former genotype). Nevertheless, there are some clear changes imposed by loss of Wnt signaling. The frequency of r1 EC-only clones (25% vs 33%) and lineages containing any r1 ECs (32% vs 48%) was reduced, while the frequency of FC-only lineages was increased (46% vs 19%; or 47% vs 31% if EC+FC lineages are included), as was the fraction of lineages containing FCs but no FSC (46% vs 28%). These indicate greater posterior shifts for precursors in the normal EC and FSC producing domains. The effect on terminal FC production is less clear, likely because of the difference in clone frequencies and consequent limitations in deducing single lineages. Overall, the formation of terminal FCs, already increased by raising JAK-STAT activity, does not appear to be enhanced by additional elimination of Wnt pathway activity, contrasting with the effects on precursors in more anterior locations.

Altogether, the loss of Wnt signaling was more influential in anterior regions, while excess JAK-STAT had a greater influence in more posterior regions. A similar conclusion was made regarding the much narrower spatial domain of FSCs in adults (Melamed and Kalderon 2020).

### Yki activity primarily affects ovarian precursor survival and cell division

We were interested in examining the roles of Hedgehog (Hh) signaling and the Hippo/Yorkie pathway for two reasons. First, both have major inputs into adult FSC division rates (Hartman *et al*. 2013; Huang and Kalderon 2014; Hsu *et al*. 2017) and we hoped to learn more about the regulation of somatic ovarian precursor proliferation. In adults, Hh stimulates FSC cell division through Yorkie (Yki) (Huang and Kalderon 2014). Second, Hh ligand is known to emanate from anterior locations in pupal ovaries (Lai *et al*. 2017), so that it might influence precursor cell migration, even though neither Hh nor Yki are known to have a significant effect on the migration or differentiation of adult FSCs.

Yki is a transcriptional co-activator frequently implicated in the regulation of cell and tissue growth. Yki activity is generally regulated, principally through inactivating phosphorylation, by upstream members of the Hippo (Hpo) pathway, including Hpo and Warts (Wts) protein kinases and a variable assortment of more upstream proteins (including Kibra) capable of sensing mechanical stimuli, including tension (Misra and Irvine 2018). However, in adult ovaries, Yki is also transcriptionally regulated by the Hh signaling pathway (Huang and Kalderon 2014). In adult FSC lineages, loss of *yki* resulted in reduced division, increased apoptosis and rapid loss of FSCs. FSC maintenance was substantially improved by expression of excess CycE and excess DIAP1, both known transcriptional targets of Yki, to counter those deficiencies (Huang and Kalderon 2014).

When *yki* mutant lineages were induced at pupariation, the proportion of ovarioles with labeled cells was lower (19%) than for the direct side-by-side control (37%) but this deficit was largely restored by addition of *UAS-DIAP1* (32%) to inhibit apoptosis. Both the reduction relative to controls for *yki* (13% vs 59%) and restoration by *UAS-DIAP1* (to 60%) were observed in a second experiment. Although variation in the frequency of clone induction is potentially quite high because of the brief and moderate heat-shock protocol, these results suggest that some *yki* mutant lineages are lost to apoptosis during pupal development.

For each experiment, FSCs were rarely present in either *yki* (0% and 6%) or *yki; UAS-DIAP1* lineages (4% and 3%) relative to control averages (15%) (Fig. 8A, H). For *yki; UAS-DIAP1*, there was no significant difference in the fraction of lineages with either r1 ECs or terminal FCs relative to controls. This suggests that there is no strong overall A/P shift in precursor outcomes, contrasting with manipulations of Wnt or JAK-STAT pathways, and that FSC loss is therefore most likely to result from reduced division rates, as observed for *cycE* and *cutlet* mutations.

**Figure 8:**
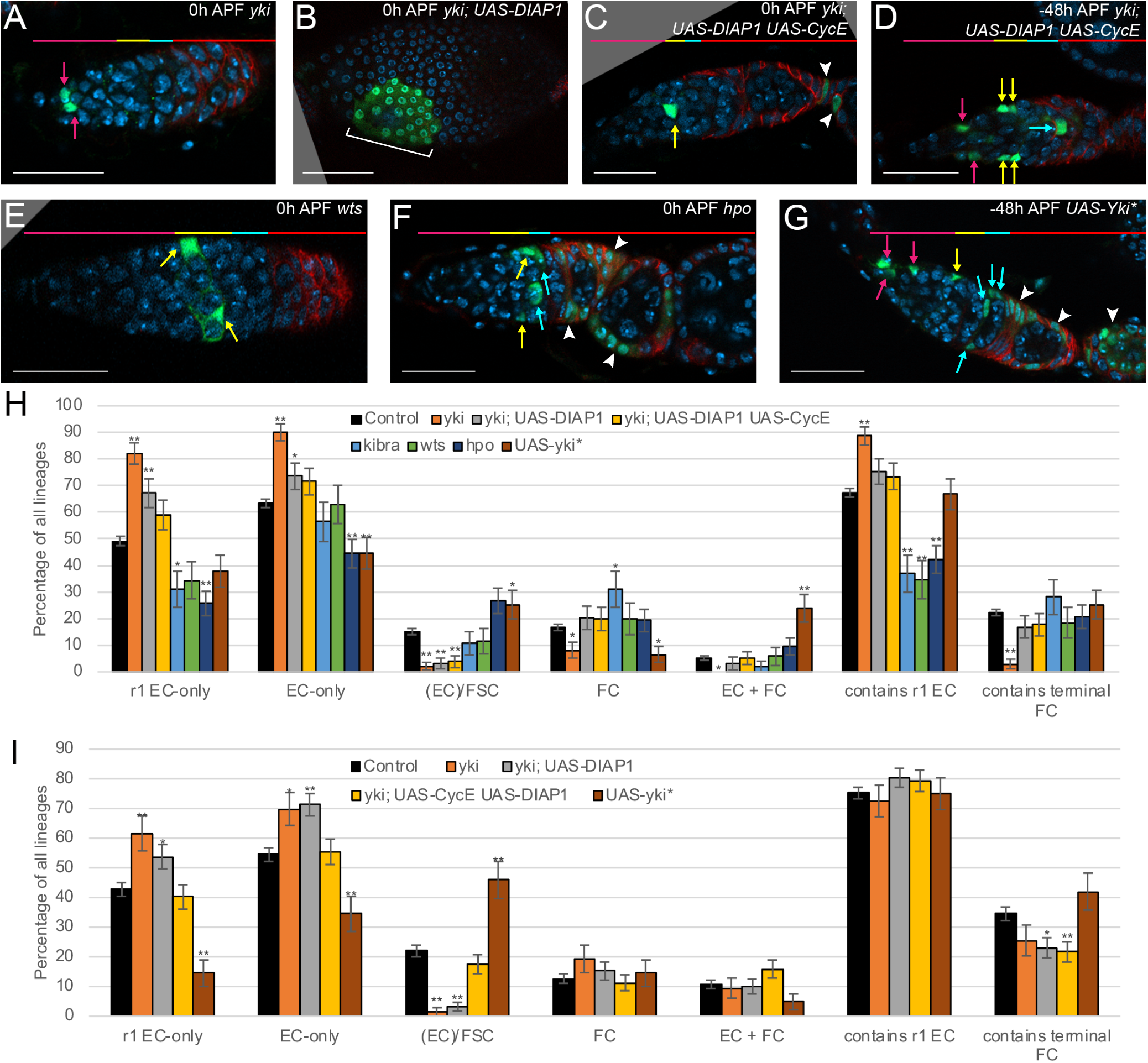
Yki activity counters posterior derivative cell apoptosis, and promotes cell division rate to increase FSC representation. **(A-G)** Representative lineages induced at (**A-C, E, F**) 0h APF or (**D, G**) 48h before pupariation for the indicated genotypes. White arrowheads indicate Follicle Cells; magenta arrows, r1 ECs; yellow arrows, r2a ECs; cyan arrows, FSCs. DAPI, blue; GFP, green; Fas3, red. The r1 EC, r2a EC, FSC and FC domains are shown in corresponding colors in the horizontal bars above images other than **(B)**. (A) *yki* commonly produced EC-only lineages, whereas **(B)** the addition of *UAS-DIAP1* greatly increased the representation of FCs, including terminal FCs, as shown here. **(C-D)** The further addition of *UAS-CycE* restored FSC-containing lineages for clones induced at **(D)** 48h before pupariation, as shown here, but not at **(C)** 0h APF; here showing an EC+FC lineage, characteristic of FSC loss. When *yki* mutant cells became FCs (generally requiring the presence of *UAS-DIAP1*)), they were frequently polar cells (arrowheads in **(C)**) but sometimes produced small patches of main-body FCs, as in **(B)**. **(E-G)** show lineages with increased Yki activity. **(E)** a *wts* lineage, illustrating the common r2a EC- only lineage, owing to a reduction in r1 EC fates, and **(F)** a *hpo* lineage, illustrating the increased frequency of EC/FSC/FC lineages, again in the absence of r1 ECs. **(G)** A UAS-Yki* (S168A) EC/FSC/FC lineage induced 48h before pupariation, illustrating the increase in such lineages and a large number of labeled cells, indicative of increased division rates. Scale Bar 20 µm for **(A, C-G)** and 50 µm for **(B)**. **(H, I)** Deduced frequency of the indicated types of single-cell lineage as a percentage of all lineages induced at **(H)** 0h APF and **(I)** -48h APF. Error bars show SEM. One asterisk (*) indicates p<0.05; two asterisks (**) indicates p<0.005 (Fisher’s two-tailed exact test). (**H**) Number of marked ovarioles = 739 (control), 84 (*yki*), 127 (*yki; UAS-DIAP1*), 67 (*yki; UAS-DIAP1 UAS-CycE*), 44 (*kibra*), 42 (*wt*s), 82 (*hpo)*, 49 (*UAS-Yki\**). (**I**) Number of marked ovarioles = 319 (control), 57 (*yki*), 101 (*yki; UAS-DIAP1*), 94 (*yki; UAS-CycE UAS-DIAP1*), 85 (*UAS-yki\**). For raw data and calculations please see supplementary spreadsheets titled 0h APF Graphs Fig4-10; -2d APF Graphs; Compilation of Numbers Figure 8 yki hpo -5d; Compilation of Numbers Figure 8 yki hpo -5d.

Other changes in lineage outcomes suggest that loss of Yki does have some effect on final A/P cell locations, either by affecting precursor migration or spatially selective apoptosis. FC-only lineages were seen at lower frequency for *yki* (6% and 10%) relative to controls (average 17%), with no increase of EC/FC lineages (1% and 0% vs 5% for controls); correspondingly, there was a large increase in EC-only clones (95% and 86% vs 63% for controls). For *yki; UAS-DIAP1*, however, the average frequency of FC-only (20% vs 17%) and EC/FC lineages (3% vs 5%) was similar to controls.

Addition of *UAS-CycE* together with *UAS-DIAP1* also resulted in frequencies of FC-only (20%) and EC/FC lineages (5%) similar to controls. Thus, the addition of *UAS-DIAP1* restored FC representation, including terminal FCs to control levels in two separate genetic tests, suggesting that posterior (would-be FCs) derivatives preferentially die in the absence of Yki. A caveat to this conclusion is that the restoration of FC lineages by *UAS-DIAP1* was seen only in one of two experiments for *yki; UAS- DIAP1* (5% and 35% for FC-only; 4% and 2% for EC/FC) (Fig. 8B, H). Moreover, the FC deficit seen for *yki* (with no *UAS-DIAP1*) in both experiments was not reproduced for lineages initiated 2d before pupariation (see below). A plausible explanation is that both a posterior bias of apoptosis induced by loss of Yki and elimination of that bias by addition of DIAP1 are variable (perhaps poised close to a threshold). In that case, the modest increase of EC-only lineage proportion and failure of lost FSC lineages to be converted to FC lineages for *yki; UAS-DIAP1*, as expected for reduced division, may be due to some residual preferentially posterior apoptosis, with no underlying direct effect of loss of Yki on A/P cell migration.

To examine further whether loss of lineages with FSCs was due to lower division rates, we tested the effect of adding *UAS-CycE* to *yki; UAS-DIAP1* lineages. This did not substantially restore FSCs (present in only 4% of lineages, Fig. 8C) for lineages induced at pupariation. However, the MARCM system introduces a delay in the accumulation of *UAS-CycE* product because perduring GAL80 gene products must first be diluted or degraded. When lineages were induced 2d prior to pupariation the frequency of FSC-containing lineages was increased from 1% (*yki*) and 3% (*yki; UAS- DIAP1*) to 18% (Fig. 8D), closer to the control average of 22% (Fig. 8I), suggesting that reduced division is indeed the primary reason for the deficit of FSCs in *yki* mutant lineages.

Other results from lineages induced 2d before pupariation showed some similarities and some differences to results from lineages induced at pupariation. The frequency of marked ovarioles for *yki; UAS-DIAP1* (74%) and *yki; UAS-CycE UAS-DIAP1* (73%) were once again higher than for *yki* (43%), suggesting some apoptotic loss of lineages for loss of Yki activity. The frequency of lineages with FCs but no FSCs (FC-only plus EC/FC) was similar to controls (23%) for the two types of *yki* lineage expressing DIAP1 (25% and 27%), as noted for clones induced at pupariation. However, *yki* lineages produced a similar result (28%) (Fig. 8I), rather than a reduction of these FC lineages, as observed for clones induced at pupariation (Fig. 8H). The fraction of lineages with labeled terminal FCs was also similar among all three *yki* genotypes (25%, 23% and 22%). Interestingly, these three values were all somewhat lower than the control average of 35%. This difference is particularly notable for the *yki; UAS-CycE UAS-DIAP1* genotype because all other lineage characteristics are similar to controls. Thus, loss of Yki results in reduced division, leading to a severe loss of FSCs, apoptosis with variable posterior cell (FC) selectivity, and a small decline in lineages with terminal FCs.

In all cases, FCs that lacked *yki* function shared some characteristics. Labeled cells were preferentially found only in polar cells (most commonly), stalk cells or the basal stalk (Fig. 8C, data not shown). Labeled FCs in other locations were approximately equally split between two categories: either a pair of cells with strongly elevated Fas3, resembling ectopic polar cells, or a relatively normal FC patch (Fig. 8B). Adult-induced lineages were previously described as frequently including polar cells or ectopic polar cells (Chen *et al*. 2011). Our results suggest that loss of *yki* greatly increase the chances that a FC derivative acquires the position or characteristics of a precursor of stalk and polar cells; cells that nevertheless become main-body FCs can develop further in a relatively normal fashion despite the absence of *yki* activity.

The consequences of increasing Yki activity were examined using strong loss of function alleles for upstream regulators that can limit Yki activity. *kibra*, wts and *hpo* lineages, induced at pupariation, shared the phenotype of a reduced fraction of both r1 EC-only lineages (31% *kibra*, 34% *wts* and 26% *hpo* vs 49% control) and lineages containing any r1 ECs (37%, 35% and 42% vs 67%, respectively) (Fig. 8H). These manipulations did not reproducibly change the proportion of lineages with marked terminal FCs (28%, 18% and 21% vs 22% control), while only specific mutations increased the proportion of lineages with FSCs (*hpo*; 27% vs 15%; others were 11%) (Fig. 8E, F, H). Neither the number of cells in EC-only lineages (*kibra* 1.8, *wts* 1.8 and *hpo* 2.2 vs control 2.7), nor the average sum of ECs and FSCs in EC/FSC/FC lineages were increased by these mutations (7.3, 9.8, 6.8 vs control 8.6). Thus, it appears that cell division rates were not significantly increased.

We also tested the effects of expressing a Yki derivative (Yki-S168A [*UAS-Yki\**]) partially resistant to down-regulation by upstream pathway components (Chen *et al*. 2011; Huang and Kalderon 2014). This resulted in an increase in lineages with FSCs (25% vs 15% control), as for *hpo* (but not the other Hpo pathway mutations), consistent with an expected consequence of an increased division rate. UAS-Yki* also produced a slightly reduced fraction of r1 EC-only lineages induced at pupariation (38% vs 49% control) but a normal fraction of lineages with any r1 ECs (67% vs 67%) (Fig. 8H). The former decrease might simply be a consequence of increased division expanding some of the most anterior clones into region 2a.

In *UAS-Yki\** lineages initiated 2d before pupariation, the frequency of r1 EC-only (15% vs 43%) and EC-only lineages (35% vs 55%) was greatly reduced while lineages with marked FSCs were greatly increased (46% vs 22%) (Fig. 8G, I). Lineages with any marked r1 ECs (75% vs 75%) or terminal FCs (42% vs 35% control) were not greatly changed in frequency. The average size of EC-only clones (3.5 vs 3.2) and the average number of ECs and FSCs in EC/FSC/FC clones (17.9 vs 14.5) were slightly increased. These results suggest that *UAS-Yki\** is more effective when induced prior to pupariation, allowing an increase in cell division rate, that greatly increases FSC production; the increase was slightly greater than achieved by excess CycE expression overv the same time period. The depletion of r1 only and EC-only lineages without affecting the frequency of lineages containing an r1 EC or an EC may result from increased clone sizes, with some spreading into more posterior domains, rather than indicating an anterior to posterior shift of all derivatives. By contrast, there was a depletion of r1 ECs amongst all clones for *kibra*, *wts* and *hpo* lineages induced at pupariation and the former two genotypes did not appear to affect cell division rate. Thus, Hippo pathway components primarily affected A/P movement of the most anterior precursors, whereas excess Yki primarily affected division rate and, hence, FSC representation.

### Hedgehog signaling promotes cell division, counters posterior derivative cell apoptosis and can influence A/P migration

The normal effect of Hh signaling on somatic ovarian precursors is most simply investigated by generating lineages homozygous for a strong *smoothened* (*smo^2^*) loss of function mutation, preventing all responses to Hh (Ingham 2022). When induced at pupariation, the proportion of ovarioles with labeled cells was consistently lower than for controls over three experiments (10% vs 28%, 12% vs 40%, 17% vs 34%). The yield of labeled ovarioles remained lower for clones of genotype *smo; UAS-DIAP1* (47% vs 73%, 19% and 17% vs 34%). Given the strong restoration of *yki* mutant clone frequency by *UAS-DIAP1* under analogous conditions, the results suggest that some *smo* mutant lineages are lost for reasons other than apoptosis or that the apoptotic signal is stronger than induced by *yki* and is not efficiently countered by *UAS-DIAP1*. The nature of recovered lineages was nevertheless influenced by the presence of *UAS-DIAP1*, as described below, suggesting some selective loss of cells by apoptosis. Expression of *UAS-DIAP1* alone did not alter any control phenotype significantly (Fig. 9D, E).

**Figure 9.**
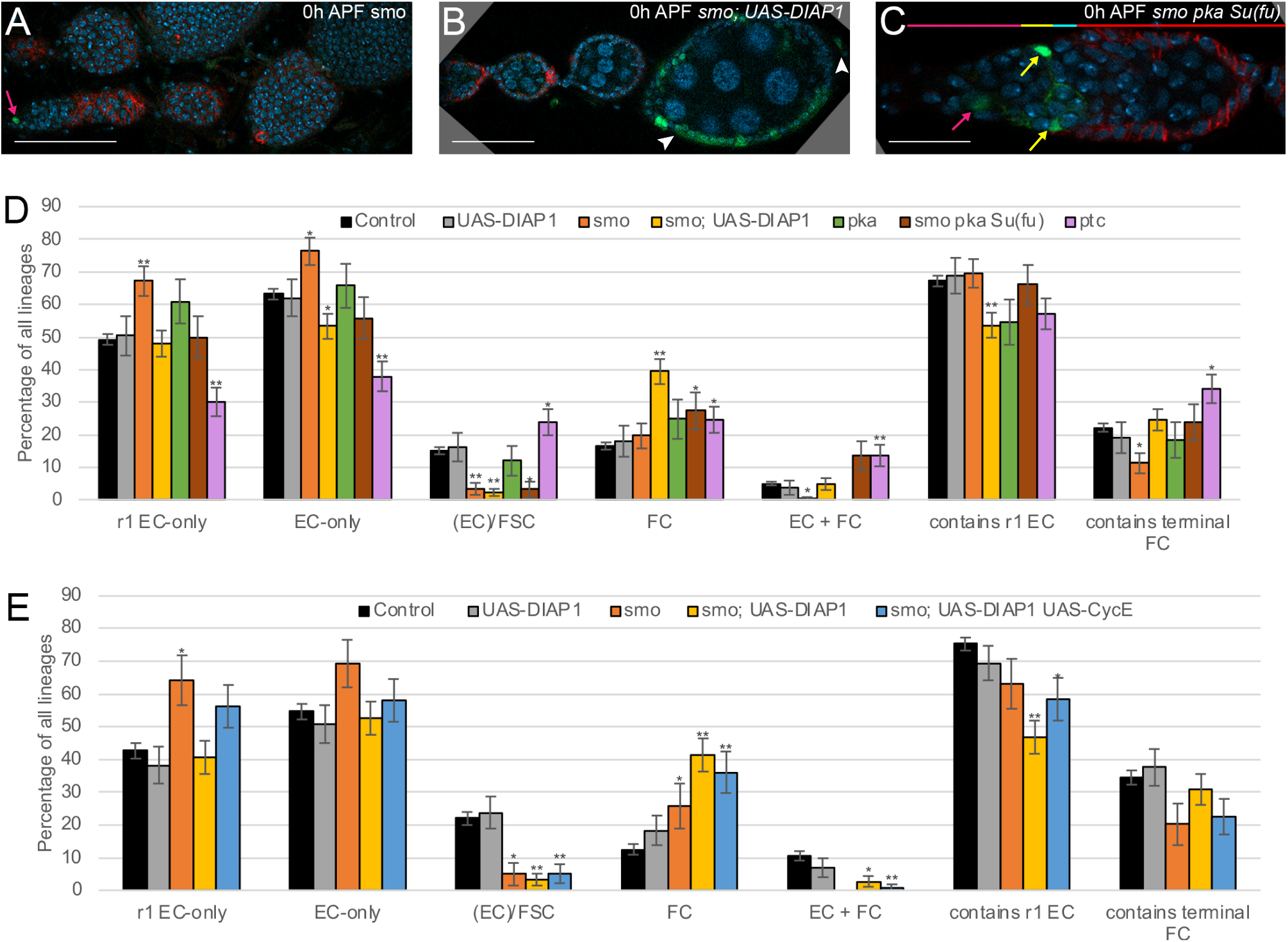
Hedgehog signaling promotes cell division and FSC representation, counters posterior derivative cell apoptosis and can influence A/P migration. **(A-C)** Representative lineages induced at 0h APF for the indicated genotypes. White arrowheads indicate Follicle Cells; magenta arrows, r1 ECs; yellow arrows, r2a ECs; cyan arrows, FSCs. DAPI, blue; GFP, green; Fas3, red. **(A)** A *smo* EC-only lineage and **(B)** a *smo; UAS-DIAP1* lineage with terminal FC marking, rarely seen in the absence of *UAS-DIAP1*. **(C)** A *smo pka Su(fu)* lineage, illustrating the common EC-only outcome. FSCs were rarely observed for all three illustrated genotypes. Scale Bar 20 µm for (**C**) and 50 µm for (**A, B**). **(D, E)** Deduced frequency of the indicated types of single-cell lineage as a percentage of all lineages induced at **(D)** 0h APF and **(E)** -48h APF. Error bars show SEM. One asterisk (*) indicates p<0.05; two asterisks (**) indicates p<0.005 (Fisher’s two-tailed exact test). (**D**) Number of marked ovarioles = 739 (control), 59 (*UAS-DIAP1*), 98 (*smo*), 142 (*smo; UAS-DIAP1*), 38 (*pka*), 51 (*smo pka Su(fu)*), 85 (*ptc*). (**E**) Number of marked ovarioles = 319 (control), 58 (*UAS-DIAP1*), 37 (*smo*), 88 (*smo; UAS-DIAP1*), 55 (*smo; UAS-DIAP1 UAS-CycE*). For raw data and calculations please see supplementary spreadsheets titled 0h APF Graphs Fig4-10; -2d APF Graphs; Compilation of Numbers Figure 9 Hh -5d; Compilation of Numbers Figure 9 Hh -5d.

The phenotype shared by *smo* and *smo; UAS-DIAP1* (each cited as the average of three tests) was a reduction of lineages that include FSCs (3% each vs 15% control) (Fig. 9A, B, D). The average number of cells per EC-only clone (1.4 and 1.4 vs 2.7) and per EC/FSC clone (1.2 and 3.1 (both from fewer than five clones) vs 8.6) were reduced (Fig. 10H), as was the average size of a terminal FC patch (*smo* 15% vs 26% control, *smo; UAS-DIAP1* 15% vs 20% control; Fig. 10J). All of these data are consistent with a reduced division rate leading to loss of FSCs. That deficit alone might be expected to produce an increase in FC-only or EC/FC lineages. The frequency of FC-only plus EC/FC lineages was indeed significantly increased for *smo; UAS-DIAP1* lineages (37% + 7% = 44% vs 17% + 6% = 23% for control) but not for *smo* lineages (19% + 1% = 20%). There was also a reduction in lineages that included terminal FCs for *smo* (11% vs 22% control) but not for *smo; UAS-DIAP1* (24%) (Fig. 9A, B, D). These results suggest that some *smo* mutant FCs, including some of those contributing to the terminal egg chamber, are lost to apoptosis, reminiscent of the *yki* mutant genotype. For *smo; UAS- DIAP1* cells, there was a small reduction in the fraction of lineages including r1 ECs (56% vs 67%). The increase in FC-only lineages (37% vs 17%) noted above was also greater than expected to result only from loss of FSCs. Hence, loss of Hh signaling in the presence of *UAS-DIAP1* appears also to result in a posterior shift in precursor derivatives. This does not appear to extend to the most posterior precursors, since the proportion of lineages with labeled terminal FCs remained almost unchanged (24% vs 22%).

**Figure 10:**
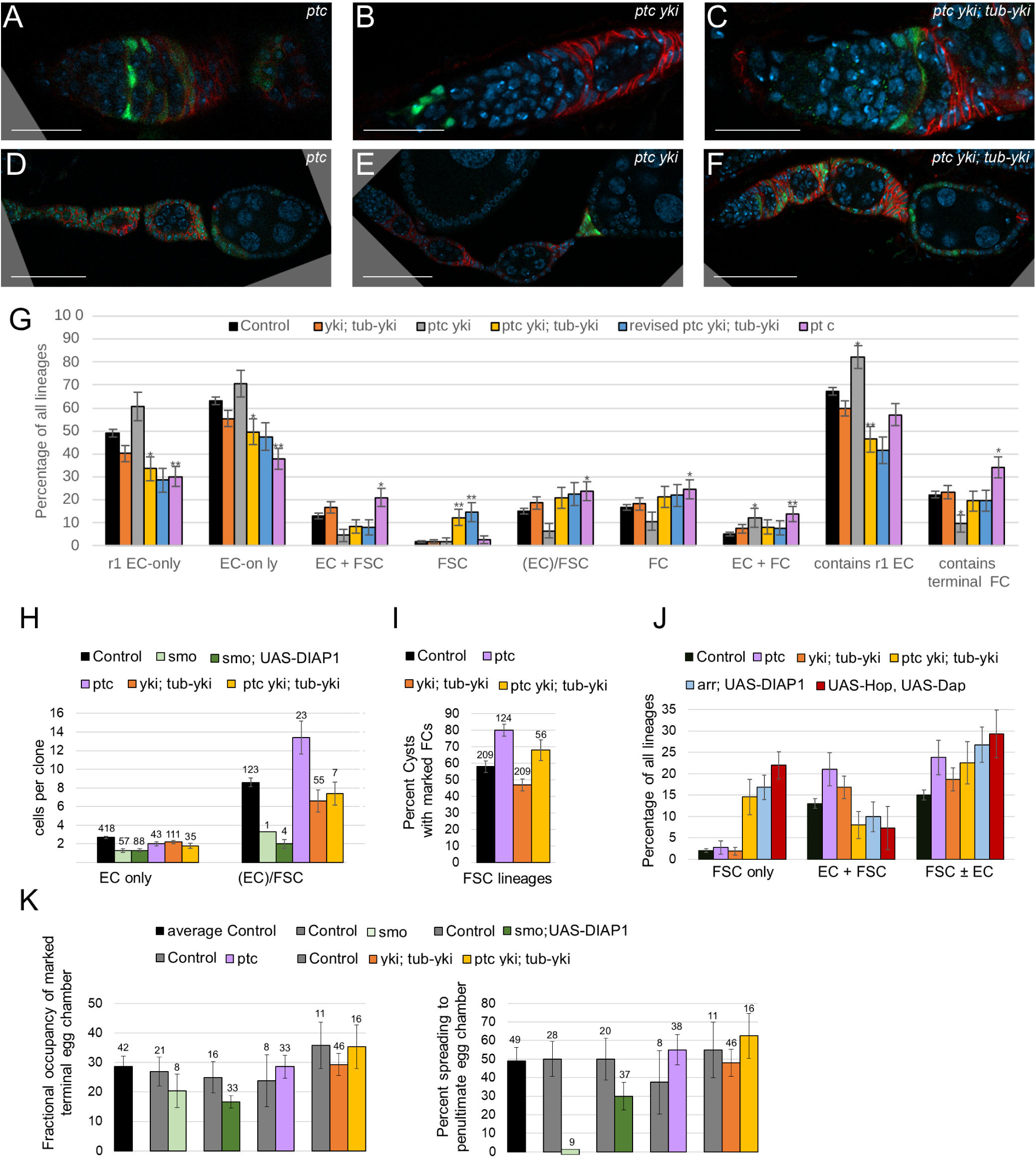
Dependence of *ptc* mutant phenotypes on transcriptional modulation of *yki*. **(A-F)** Representative lineages induced at 0h APF for the indicated genotypes. White arrowheads indicate Follicle Cells; magenta arrows, r1 ECs; yellow arrows, r2a ECs; cyan arrows, FSCs. DAPI, blue; GFP, green; Fas3, red. The r1 EC, r2a EC, FSC and FC domains are shown in corresponding colors in the horizontal bars for **(A-C). (A)** A *ptc* EC/FSC/FC lineage showing a large number of marked cells, likely due to increased division. **(B)** A *ptc yki* EC-only lineage resembling a common *yki* phenotype (Fig. 6A), and **(C)** a *ptc yki; tub-yki* EC/FSC/FC lineage with characteristically fewer cells than for *ptc*. **(D-F)** Ovarioles with *ptc* mutations induced at 0h APF produced ovarioles with multilayered FCs (paired white arrowheads) for **(D)** *ptc*, **(E)** *ptc yki* (which has relatively few marked FCs) and **(F)** *ptc yki; tub-yki*. Scale Bar 20 µm for **(A-C)** and 50 µm for **(D-F)**. **(G)** Deduced frequency of the indicated types of single-cell lineage as a percentage of all lineages induced at 0h APF. Error bars show SEM. One asterisk (*) indicates p<0.05; two asterisks (**) indicates p<0.005 (Fisher’s two-tailed exact test). Animals occasionally lose the *tub-yki* transgene. Hence, data are reported for all scored ovaries for the *ptc yki; tub-yki* genotype and also as “revised” after removing samples from animals thought likely to have lost the *tub-yki* transgene based on phenotypes (low clone frequency, low frequency of FSCs and FCs) of all examined ovarioles. Number of marked ovarioles = 739 (control), 178 (*yki; tub-yki*), 59 (*ptc yki*), 76 (*ptc yki; tub-yki)*, 65 (revised *ptc yki; tub- yki*), 85 (*ptc*). **(H-K)** Measures of division rate and composition of FSC-containing lineages. **(H)** Average number of ECs in EC-only lineages, and the average sum of EC and FSC number in FSC-containing lineages. SEMs are based on raw data (prior to correction for single-cell lineages). n values (number of lineages scored) are above columns. **(I)** Percentage of cysts (includes region 2b, region 3 and all egg chambers) with adjacent marked FCs among FSC-containing lineages. n values (number of cysts scored) are above columns. **(J)** Percentage of all lineages with marked FSC but no EC, marked EC and FSC, marked FSC with or without EC. Number of marked ovarioles as in (G) for first four genotypes; 95 for *arr; UAS- DIAP1* and 50 for *UAS-Hop UAS-DIAP1*. **(K)** Left: average percentage of all FCs around the terminal egg chamber that are marked among lineages that include terminal FCs (but excluding those with only polar cells, stalk cells or basal stalk cells). Right: percentage of lineages with marked terminal FCs (no exclusions) with marked FCs on the penultimate egg chamber. n values (number of terminal egg chambers scored) are above columns. For raw data and calculations please see supplementary spreadsheets titled 0h APF Graphs Fig4-10; FC occupancy aggregate; Division Measures Hh/Yki; Compilation of Numbers Figure 10 Hh Hpo.

The phenotypes of *smo* and *smo; UAS-DIAP1* lineages were reproduced when induced 2d prior to pupariation. Clone frequency was reduced (17% and 21%, respectively versus 53% for the control, which expressed *UAS-DIAP1*) and lineages with FSCs were greatly reduced in both cases (1% and 1% vs 22% control average, Fig. 9E). The fraction of lineages with marked FCs in the absence of FSCs (FC- only plus EC/FC) was significantly increased only in the presence of DIAP1 (*smo* 26% and *smo; UAS- DIAP1* 44% vs 23% control), with a marked decrease in representation of terminal FCs amongst all lineages only in the absence of *UAS-DIAP1* (*smo* 20% and *smo; UAS-DIAP1* 31% vs 35% controls, Fig. 9E). Again, there was a decrease of lineages containing r1 ECs when DIAP1 was present (47% vs 75% control). These results are consistent with a posterior shift of all but the most posterior FC precursors in the absence of Hh signaling, together with some loss of FCs through apoptosis. This posterior shift would by itself increase FSC representation, based on the *arr* mutant phenotype, so the observed loss of FSCs is almost certainly due to reduced division rates. The addition of *UAS-CycE* to the *smo; UAS- DIAP1* genotype did not, however, increase FSC representation (Fig. 9E), contrasting with the analogous rescue observed for *yki* mutant lineages. One possibility is that the reduction in cell division due to loss of *smo*, which may be more profound than from loss of *yki* (consistent with a greater FSC deficit), is not rescued effectively or in a timely fashion by *UAS-CycE*.

We also investigated the effects of partially compromising Hh signaling. In *Drosophila* wing discs, the loss of Protein Kinase A (PKA) compromises Smo activation by Hh but also blocks proteolytic processing of the full-length latent transcriptional activator Ci to a repressor form (Jia *et al*. 2003; Apionishev *et al*. 2005; Smelkinson *et al*. 2007; Little *et al*. 2020). It therefore produces an intermediate level of Hh pathway activity (high levels of full-length Ci but limited activation of Ci-155 by Smo-activated Fused kinase). Similarly, *smo pka: Su(fu)* mutant cells allow no activation of Fused but partially alleviate the requirement for Fu kinase activity because of the absence of Su(fu) (Suppressor of fused) restraint on full-length Ci activity (Ohlmeyer and Kalderon 1998). In wing discs, both *pka* and *smo pka; Su(fu)* genotypes produce relatively high levels of pathway activity but lower than maximal Hh signaling (manifest as strong transcriptional *patched* (*ptc*) induction but no induction of anterior Engrailed). In developing ovarioles, there was no significant reduction in clone frequency for these lineages induced at pupariation. Both genotypes led to a reduced fraction of lineages containing FSCs (12% and 3% vs 15% control, Fig. 9D). The effect was strongest for *smo pka Su(fu)* (Fig. 9C), suggesting that precursors in locations destined to produce FSCs normally receive high levels of Hh during some or all of pupal development. The FSC deficit was accompanied by an increase in lineages with FCs but no FSCs for *pka* and *smo pka; Su(fu)* (25% and 41% vs 22% for controls), without large changes in the fraction of lineages with terminal FCs (18% and 24% vs 22% control) or r1 ECs (54% and 66% vs 67%) (Fig. 9D). Thus, even intermediate reductions in Hh pathway activity reduce FSC production, likely through reduced cell division rates, while complete loss of pathway activity also results in a posterior shift of all but terminal FC precursors, accompanied by some loss of posterior precursor lineages through apoptosis.

The maximal physiological level of Hh pathway activity is generally reproduced by loss of *ptc* function (Ingham 2022). The phenotype of *ptc* mutant lineages induced at pupariation was consistent with an elevated division rate. The fraction of lineages that included an FSC was increased (24% vs 15% control, Fig. 9D, Fig. 10A), as was the average size of an EC/FSC lineage (13.4 vs 8.6 control EC plus FSC cells; Fig. 10H). The fraction of all marked ECs and FSCs that were FSCs was also higher (28% vs 19% control average). The average size of a terminal FC patch was slightly higher (29% vs 24%; this measure excludes clones exclusively composed of polar cells, stalk cells or basal stalk cells; Fig. 10K). Since *ptc* marked FCs were generally very abundant (Fig. 10D) we wondered if these measures fully represented the division rate of *ptc* mutant cells. We therefore also measured indicators of the spread of clones as a further indicator of division rate. A precursor at pupariation that is destined to contribute to the terminal egg chamber also sometimes contributes to the penultimate egg chamber. As noted in the first section of Results, the lineage sometimes also contributes FCs occupying more anterior positions. A lineage that produces more cells is more likely to spread into each of these locations. Control lineages contributing to the terminal FC also populated the penultimate egg chamber with a frequency of 38% and more anterior egg chambers with a frequency of 25%. Those values were increased to 55% and 66%, respectively for *ptc* lineages (Fig. 10L). This measure does not correct for single-cell lineages and in other experiments the control values for penultimate egg chamber occupancy were higher (to give an average of 49%) (Fig. 10L). Thus, our cumulative evidence suggests that loss of *ptc* function may not increase the division rate of the most posterior precursors. The spread of FSC-containing lineages was estimated by counting the fraction of cysts with adjacent marked FCs. This value was much higher for *ptc* (80%) than for control FSC-containing lineages (Fig. 10I), supporting the deduction from marked EC and FSC numbers in such lineages (Fig. 10H) that *ptc* increases division rate for precursors in the regions that can produce FSCs. The average size of an EC- only clone was not increased for *ptc* lineages (2.0 vs 2.7 control cells; Fig. 10H), suggesting that division is not increased in the most anterior regions, closest to the source of Hh.

There also appears to be a posterior bias in the distribution of *ptc* mutant lineages, with fewer lineages including r1 ECs (57% vs 67% control) and more lineages including terminal FCs (34% vs 22%), reflected also in an increased fraction of lineages containing FCs but no FSC (38% vs 22%) (Fig. 9D). A posterior shift can also contribute to an increase of lineages with FSCs, as observed for *arr* lineages. Nevertheless, the *ptc* phenotype shows a lesser posterior shift than *arr*, a greater increase in EC/FSC lineages and some direct evidence of increased division rates, affirming a significant contribution of division rate to FSC representation. The major effect of Hh signaling on cell division rate and FSC production was manifest by opposite *smo* and *ptc* phenotypes. Effects on cell location appear to be more complex: increased Hh signaling caused posterior displacement throughout the precursor domain, while loss of Hh signaling produced the same result (not the converse, as might be expected) for all but the most posterior precursors.

### Relationship between Hh signaling and Yki activity

In adult FSC lineages, *ptc* increased *yki* RNA levels, *yki* was fully epistatic to *ptc* and substituting *yki* with a *tub-yki* transgene, preventing normal transcriptional regulation of *yki*, suppressed all *ptc* phenotypes (to produce roughly wild-type FSC properties) (Huang and Kalderon 2014). It was therefore concluded that *ptc* increased FSC representation (through enhanced division) via transcriptional induction of *yki*. Another report suggested a post-transcriptional regulation of Yki activity through the Hh pathway (Li *et al*. 2015), while the study of adult ECs deficient for *yki* or *smo* activity led to a conclusion of independent (albeit similar) Hh and Yki effects, each leading to EC dysfunction and consequently delayed or deficient germ cell differentiation (Huang *et al*. 2017).

We explored the relationship between Hh signaling and Yki activity in ovarian precursors by manipulating both simultaneously. *ptc yki* lineages initiated at pupariation had phenotypes similar to *yki* phenotypes (Fig. 8A, Fig. 10B). Lineages with FSCs were less frequent than for controls (6% vs 15%; *yki* 2%), as were FC-only lineages (11% vs 17%; *yki* 8%) and the subset with only terminal FCs (5% vs 10%; *yki* 3%), while EC-only lineages were more frequent (71% vs 63%; *yki* 90%), as were r1 EC-only lineages (61% vs 49%; *yki* 82%) (Fig. 10G). Correspondingly, lineages that included labeled terminal FCs were less frequent (10% vs 22%; *yki* 3%) and those with r1 ECs were more abundant (82% vs 67%; *yki* 89%). All of these deviations from wild-type behavior are similar to those of *yki* lineages, though smaller in magnitude, and opposite to changes characteristic of *ptc* alone (Figs. 8 and 10). Thus, loss of *yki* is largely epistatic to loss of *ptc*, regarding the distribution of lineage types. All terminal FC samples for *ptc yki*, like *yki*, were mostly composed of polar cells or basal stalk cells, so a reliable measure of division rate for terminal epithelial FC precursors is not available. Similarly, there were too few FSC-containing lineages (just three) to provide a reliable estimate of division rate of EC/FSC precursors. Thus, the loss of FSC-containing *ptc yki* lineages is plausibly due to a reduced rate of precursor division, as deduced for *yki* lineages through rescue with *UAS-CycE*, but there is no direct supporting evidence.

The phenotype of marked *ptc yki* FCs commonly included the characteristic *yki* phenotype of occupying polar cell locations or producing ectopic polar-like cells with high Fas3, but also included multi-layering (Fig. 10E), with the marked cells in an outer layer, or the presence of multiple excess labeled cells between egg chambers. Those phenotypes were previously noted for *ptc* mutant FCs induced in adults (Zhang and Kalderon 2000) and were evident also in *ptc* mutant pupal lineages (Fig. 10D), where the FC patches were generally much larger than for *ptc yki* (supporting the hypothesis of a significantly lower division rate). We deduce that the multi-layering phenotype, indicative of a different affinity of *ptc* mutant FCs for germline cells and wild-type FCs, is not dependent on Yki activity.

Substitution of *yki* function with *tub-yki* resulted in lineages with phenotypes similar to controls (Fig. 10G). This result suggests that the levels of *yki* provided by the transgene are roughly normal (consistent with maintenance of healthy stocks with *tub-yki* replacing endogenous *yki* activity) and that *yki* need not be regulated transcriptionally for roughly normal ovarian somatic cell precursor behavior.

We noticed that the *tub-yki* transgene, marked by *w^+^*, is sometimes spontaneously excised in stocks. This is, unfortunately not directly visible in stocks carrying the *yki^B3^* allele, which itself has strong *w^+^* expression. We noted, however, that in our initial tests, many lineages expected to be of genotype *yki; tub-yki* showed characteristic *yki* mutant FC phenotypes (a high proportion of polar and stalk cells or elevated Fas3 expression). We therefore obtained the lineage results cited above by constructing a stock that is homozygous for *yki* together with one copy of *tub-yki*, so that the active transgene must be maintained for viability. The unstable *tub-yki* transgene also was lost over time from *ptc yki; tub-yki* stocks, as revealed by crosses to *yki* stocks to test viability, but here there was no solution to enforce *tub-yki* retention. We therefore measured *ptc yki; tub-yki* lineage phenotypes for two recently-derived stocks. In two flies derived from each stock, lineages resembled those seen for *ptc yki* (we score ovaries from individual flies separately). Results are therefore presented without any adjustment, likely representing a mix of *ptc yki; tub-yki* and *ptc yki* genotypes, and also after discarding the four sets of ovarioles very likely lacking *tub-yki*. Results were not greatly altered by the latter step, but we discuss here the second set of results (“revised *ptc yki; tub-yki*"), which likely represents the correct genotype.

The addition of *ptc* (to *yki; tub-yki*) increased the fraction of lineages with FSCs (Fig. 10C, F) from 18% to 23%, both higher than controls (15%) but not significantly (p>0.05) (Fig. 10D, J). The average number of cells per EC/FSC clone (7.4) was similar to the *yki; tub-yki* genotype (6.6) and to controls (8.6) but much lower than for *ptc* (13.4) (Fig. 10H). To probe division rate further in the neighborhood of precursors that can yield FSCs we also counted the frequency of marked FC patches in FSC-containing lineages. The very high proportion of germline cysts contacting marked FCs for *ptc* lineages (99/124; 80%), was not matched by *ptc yki; tub-yki* (38/56; 68%), yielding a value similar to the direct control for that experiment (35/51; 69%), slightly higher than the average of six such controls (121/209; 58%), and higher than for *yki;tub-yki* (98/209; 47%) (Fig. 10I). Thus, marked cells in FSC-containing lineages were less abundant in *ptc yki; tub-yki* ovarioles than for *pt*c when assessed both by EC/FSC numbers and a measure of FC production, suggesting that the faster division promoted by *ptc* is largely suppressed by *yki* substitution with *tub-yki*. That is consistent with the possibility that *ptc* increases division of precursors in intermediate FSC-producing locations largely through transcriptional induction of *yki*, as in adult FSC lineages. The average terminal FC clone size (35%) was very similar for *ptc yki; tub-yki* and the direct control (36%) but higher than for *yki; tub-yki* (29%) in the same experiment (Fig. 10K). The percentage of terminal FC clones that spread to the penultimate egg chamber was higher for *ptc yki; tub-yki* (63%) than both the control (55%) and *yki; tub-yki* (48%) (Fig. 10K). Thus, there may be a small loss of division of posterior precursors for *yki; tub-yki* and a small increase due to loss of *ptc* that does not rely on *yki* transcriptional induction. The phenotype of marked *ptc yki; tub-yki* FCs included multilayering and the accumulation of cells between egg chambers (Fig. 10F), confirming that these characteristics are caused by Hh signaling without using *yki* as an intermediate.

The fractions of lineages including r1 ECs (47% vs 67%) and of EC-only lineages (48% vs 63%) were lower than controls, while the frequency of FC-only lineages was increased (22% vs 17%) for *ptc yki; tub-yki*, as observed for *ptc* alone (Fig. 10G). However, terminal FC labeling among all lineages was similar to controls (20% vs 22%), unlike *ptc* (34%). These results suggest that the posterior bias produced by excessive Hh signaling is not dependent on *yki* induction for precursors other than the most posterior, potential terminal FC precursors. It is interesting that loss of *yki* can lead to reduced terminal FC representation, while excess Yki increased representation (Fig. 8I). Those results are consistent with the possibility of *ptc* increasing terminal FC representation by inducing *yki*.

Conversely, it is possible that reduced *yki* induction in *smo* mutant cells reduces terminal FC representation. This might balance a *yki*-independent consequence of *smo* that produces net posterior movement to result in the observed *smo* phenotype of posterior displacement of all but the most posterior clones. If correct, *smo* and *ptc* would have opposite *yki*-dependent changes in terminal FC formation, with Yki acting positively to promote terminal FC identity. For more anterior precursors, *smo* and *ptc* would both promote posterior movement in a Yki-independent manner.

Increased posterior movement in the EC/FSC precursor domain, together with a roughly normal rate of cell division might account for the unusually high proportion of *ptc yki; tub-yki* lineages with labeled FSCs but no labeled ECs (15% vs 2% for controls, Fig. 10G, J). The other genotypes with a similar phenotype were *arr* (loss of Wnt signaling) and *UAS-DAP UAS-Hop* (increased JAK-STAT with an inhibitor of increased division), which causes posterior movement of all precursors with no apparent effect on cell division (Fig. 10J). The phenotype is not shared by *ptc*, presumably because increased division additionally replenishes EC precursors in EC/FSC lineages to balance posterior loss. Thus, although FSC-containing lineages are not much more abundant for *ptc* than for *ptc yki; tub-yki*, the contribution of increased division rate, which appears to rely on *yki* induction, is manifest by the markedly different frequencies of EC/FSC lineages (Fig. 10G and 10J).

## Discussion

The replenishment of lost differentiated cells by adult stem cells is a vital physiological function for tissues with high cellular turnover, including human epidermal and intestinal cells. Adult stem cells must execute this function over a lifetime, overcoming natural variations and occasional extreme challenges, while their necessary continued replication and potential longevity make stem cells vulnerable to the accumulation of cancer-causing mutations. Adult stem cell studies are slowly revealing the cellular organizations and behaviors selected by these evolutionary pressures. In one large class of paradigms, which includes Drosophila FSCs and mammalian intestinal stem cells, stem cell division and differentiation are independent (Jones 2010; Reilein *et al*. 2017; Clevers and Watt 2018; Reilein *et al*. 2018; Beumer and Clevers 2021; Greulich *et al*. 2021; Kalderon 2022). Hence, maintenance of a specific stem cell requires that division competes successfully with differentiation; this balance can be altered by genetic mutation, to produce a functional stem cell deficit or a hyper- competitive lineage with oncogenic potential. Moreover, differentiation in these paradigms is governed by the spatial distribution of extracellular signals and is initiated by changes in cell location (along the AP axis for FSCs and the crypt bottom/top axis for intestinal stem cells).

Equally relevant, but less studied, are the developmental processes that create the requisite organization of stem cells and niche cells that produce key extracellular signals. Drosophila FSCs belong to a class of paradigm where stem cells are not set aside during development (Guiu *et al*. 2019; Obernier and Alvarez-Buylla 2019; Bond *et al*. 2021; Reilein *et al*. 2021). Instead, a set of precursors simultaneously produces stem cells and the adult differentiated cells they will support. If differentiated cells are continuously irreversibly lost from the precursor pool through migration away from the future stem cell domain, as is true for FCs in the pupal Drosophila ovary and likely holds also for mammalian intestine, the survival of a specific precursor to produce a stem cell may depend on sufficient division activity during development to counter differentiation losses (Fig. 11 and Fig. 12). Also, if stem cell division and migration (leading to differentiation) must be balanced during development and in adults, the use of a similar set of extracellular factors might be expected. Our studies of the responses of single cells to specific genetic manipulations during Drosophila ovary development support both of these hypotheses (Fig. 11). These conclusions, based on experimental evidence for FSCs, are likely to apply generally to tissues supported by stem cells that develop alongside the mature tissue and act as a community with independent regulation of division and differentiation (Fig. 12).

**Figure 11:**
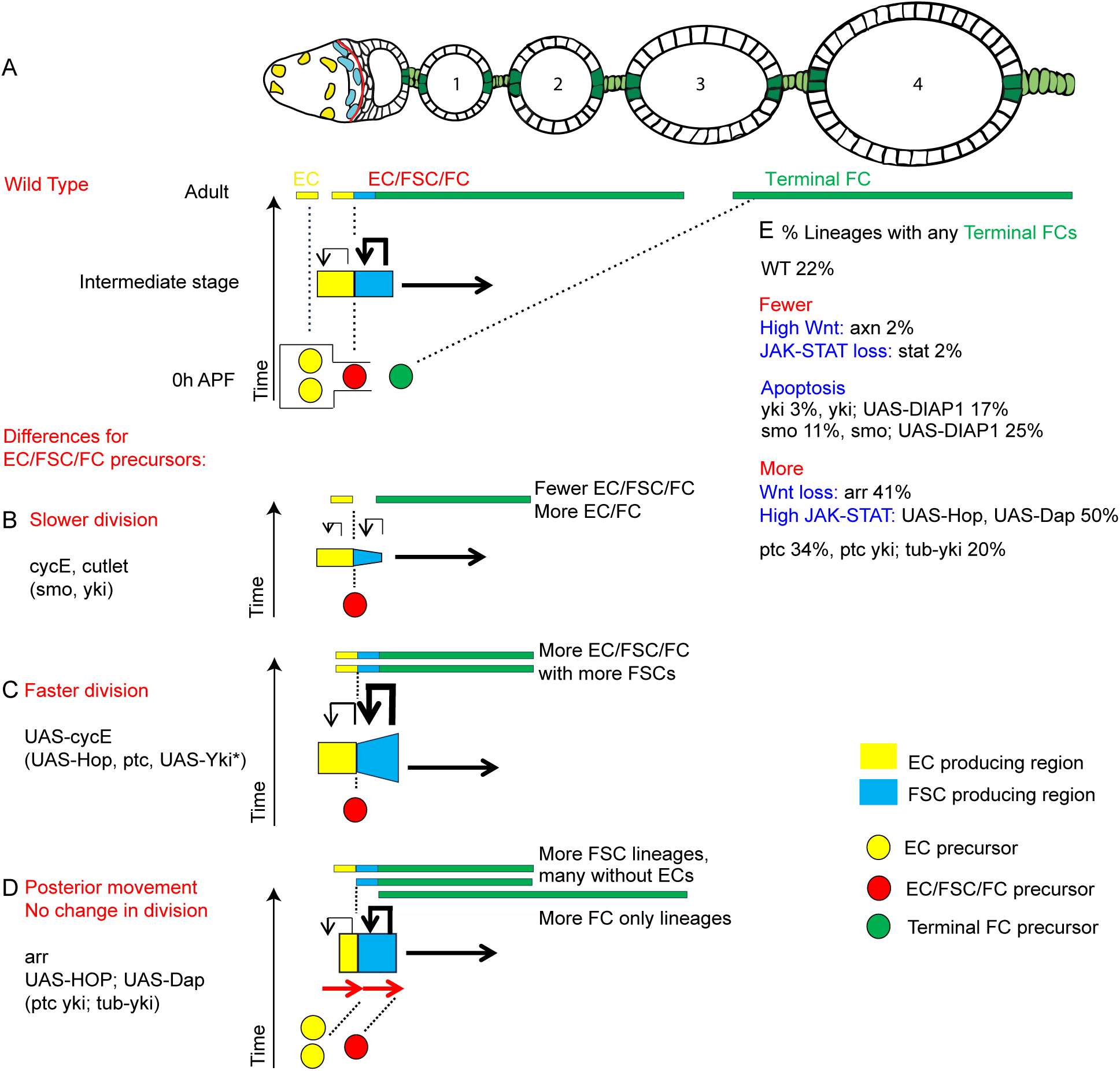
Summary of key influences on FSC formation and formation of the first FCs. **(A)** Ovariole diagram with ECs in yellow, FSCs in cyan, specialized stalk and polar cell FCs in green and other FCs in white. Representative precursors of EC-only (yellow), EC/FSC/FC (red) and terminal FC lineage (green) precursors are shown for oh APF. EC-only precursors are the most abundant, so two yellow cells are shown. An intermediate developmental time (such as 48h APF) is shown for EC/FSC/FC precursors to highlight the increased A/P range of derivates destined to become ECs (yellow) and FSCs (cyan), as well as the constant loss of EC/FSC precursors to become FCs to the posterior (horizontal arrow). The more posterior EC/FSC precursors divide faster than anterior EC/FSC precursors, especially later in pupation (larger renewal arrow). Our studies show that this renewal is important for FSC precursors to persist, compensating for losses from continued FC formation. **(B-D)** Differences from wild-type behavior that affect precursors becoming FSCs. Experimental results are summarized and the underlying rationalization is illustrated (showing only relevant precursors). **(B)** Cell-autonomous reduction of division rate of an EC/FSC/FC (red) precursor among other wild-type precursors leads to progressive loss of FSC precursors throughout pupation because FSC precursor division no longer matches losses from differentiation to FCs. Consequently, many potential EC/FSC/FC lineages become EC/FC lineages. This behavior was seen most clearly for reduced activity of *cycE* and *cutlet*, but also for *smo* and *yki* when apoptosis was inhibited. **(C)** Conversely, a genetic cell-autonomous increase in division rate relative to other precursors leads to a progressive increase of FSC precursors during pupation, ensuring survival of FSC precursors and giving rise to EC/FSC/FC lineages with more FSCs than usual. Experimentally, this condition increased the frequency of EC/FSC/FC lineages, implying that some potential FSC lineages are normally lost in wild-type ovarioles because of insufficient division. This behavior was seen most clearly for increased activity of *cycE.* Similar results were seen for excess activated Yki (but not for regulators of Hippo pathway activity expected to increase Yki activity) and lesser, similar changes were observed for *ptc*. Excess JAK-STAT (*UAS-Hop*) produced many FSC-containing lineages but there were too many marked cells altogether to quantify lineage results. (**D**) A genetic change deduced to cause cell-autonomous posterior movement of all precursors (as seen for *arr*) resulted in fewer EC-only lineages, more FSC-containing lineages and more FC-only lineages. The increase in FSC-producing is because there are initially more potential EC-only (yellow) precursors than EC/FSC/FC precursors. Hence, a posterior shift relative causes more precursors originally in EC-only locations to move into FSC-producing territory than the number of precursors moving out of EC/FSC/FC territory to become FCs. Many of the resulting FSC-containing lineages lacked ECs. That is presumably because precursors entering the EC/FSC/FC-producing domain can continue to move posteriorly through that domain (red arrows) and are sustained, in part, through normal rates of cell division. Division continues in the FSC-producing (cyan) region but not in the EC-producing (yellow) region in the second half of pupation, so that many lineages maintain FSCs but not ECs. A high proportion of FSC lineages without ECs was seen also for *UAS-Hop; UAS-Dap* and *ptc yki; tub-yki* genotypes, supporting the deduction that they cause posterior movement around the FSC precursor region without affecting cell division rate. **(E)** The most posterior ICs at pupariation (green in (**A**)) were deduced from lineage studies here to migrate out of the germarium over the first 48h of pupation and then become the first FCs, occupying the terminal egg chamber (and sometimes also the penultimate egg chamber). The frequency of this outcome (producing Terminal FCs) was greatly decreased by loss of JAK-STAT or increased Wnt signaling; it was increased by reciprocal changes in those pathways. Reduced terminal FC contributions in the absence of *yki* and *smo* activity appeared to be due to spatially selective apoptosis. Increased Hh signaling (*ptc*) increased terminal FC lineage frequency, apparently dependent on normal *yki* gene regulation.

**Figure 12.**
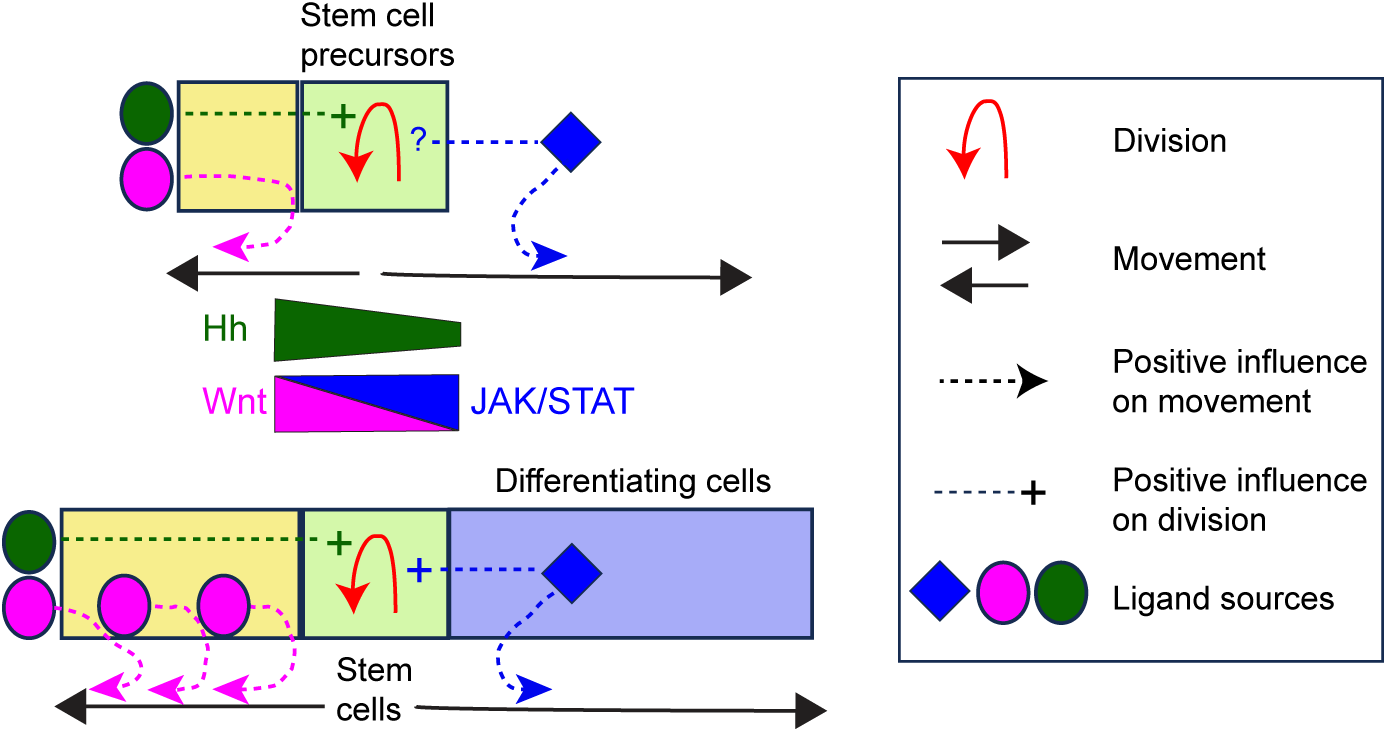
General principles illustrated by FSC development. The cartoon depicts a pool of precursors on top and the adult tissue they form at the bottom. If a set of precursors (in the pale green space) produce both stem cells and differentiated tissue extending away from the source during development, then the probability that a given precursor will remain in the stem cell zone and become an adult stem cell depends on its division rate competing successfully against movement out of the stem cell zone. In paradigms with independent division and differentiation of adult stem cells, the same competition applies to maintaining a given stem cell. Both division rate and cell movements are likely to be regulated by external signals. It would generally be economical to use largely similar signals to regulate those processes during both development and adulthood, as we have found for Drosophila ovarian FSCs. Most simply, this is achieved by signals from niche cells established early in development. In the Drosophila ovary, two such signals come from Cap cells, which form prior to pupation: Hh promotes division and Wnt promotes anterior movement. There are two other features of ovary development, which might also be used in other paradigms. First, the primitive Cap cell niche is elaborated or expanded during pupal development by the production of additional niche cells (Escort Cells; pale yellow region) that supplement signaling from the anterior by producing additional Wnt ligands. Second, there is an important signal in adults (activating the JAK-STAT pathway) that originates from differentiated cells posterior to the stem cells. This signal promotes both stem cell division and posterior exit from the stem cell zone. During development, the same pathway promotes exit from the stem cell zone and has the potential to stimulate division; our data do not show whether division is stimulated at the earliest times during pupation or what is the exact cellular source of JAK-STAT pathway ligand prior to production of the first differentiated cells. Some stem cell paradigms, like mouse gut crypts, may not have the feature of FSCs that differentiated cells are produced in two directions (the yellow territory may be absent altogether or not replenished by the stem cells). The general principles still remain that (i) division rate is important for both development and maintenance of stem cells because it competes with cells moving out of the stem cell zone, and (ii) niche cells may produce a set of signals that act in a graded fashion over distance to regulate division rate and movement of both stem cell precursors and their adult stem cell derivatives.

### Origin of the first FCs

It was previously shown that most FC-producing lineages induced at 36h APF contributed either no terminal FCs or only terminal FCs, suggesting segregation of terminal FC precursors by that stage (Reilein *et al*. 2021). This deduction was supported by live imaging, showing that only cells posterior to germline cells (in the Extra-Germarial Crown and Basal Stalk) from about 48h APF populated the terminal egg chamber (Reilein *et al*. 2021). Some of these cells accumulated anterior to the terminal egg chamber, destined to become FCs on the penultimate egg chamber, consistent with lineage studies. By contrast, FCs in more anterior egg chambers, together with ECs and FSCs, derived from germarial ICs. In this work, we found that lineages induced at pupariation often included marked FCs in the terminal egg chamber and also anterior to the penultimate egg chamber. The high frequency of those occurrences could not be explained by chance induction of two separate lineages; instead, most such occurrences report the behavior of single-cell lineages. Hence, we deduce that somatic cells intermingled with germline cells (ICs) in the developing germarium at pupariation can give rise to adult somatic ovary cells in every location, from the most anterior ECs to the most posterior FCs and the basal stalk protruding from the posterior of the terminal egg chamber (Fig. 11A). Accordingly, the terminal FC precursors seen accumulating over the next 48h must largely derive from migration of ICs out of the germarium, distancing themselves from germline cells, even though such movements have not been directly observed by live imaging. Our lineage studies cannot exclude the possibility that some of the terminal FC population derive from a separate source but there is no specific evidence supporting that possibility. It seems most likely that all terminal FCs derive from ICs at pupariation.

The alternative potential source of terminal FCs previously considered were “basal cells”, already present posterior to developing germaria at pupariation but without the characteristic Fas3 expression of cells in the subsequently enlarging Extra-Germarial Crown. Recent studies identified a population of cells, named Swarm cells, in larval ovaries, according to selective expression of Single-minded and Crossveinless-2 (Banisch *et al*. 2021). By using corresponding antibodies and GAL4 lines for lineage tracing, it was shown that these apical cells migrate basally from 72h to 120h after egg-laying to form the “basal cell” population at the larval/pupal transition. Swarm cell migration is initiated by an ecdysone pulse, while a second ecdysone pulse triggers the now-posterior Swarm cells to send a differentiation signal to posterior PGCs. Thereafter, Swarm cell lineages are lost during pupal development, leaving little residue in adults, consistent with an earlier surmise of basal cell death during pupation from electron micrographs (King *et al*. 1968). These deductions are consistent with our conclusion that ICs, and not basal cells, are the precursors of terminal FCs.

### Cell division rate strongly affects the likelihood of a precursor becoming a FSC

Division rate is a key parameter for adult stem cells. For FSCs and all adult stem cells maintained by population asymmetry with division-independent differentiation, a genetic change that alters the division rate of a single stem cell necessarily has a strong effect on the likelihood that the stem cell lineage will amplify, survive or be extinguished (Reilein *et al*. 2018). A stem cell with increased division rate will likely be maintained, with an elevated chance of amplifying to occupy an entire niche, immortalizing the genetic change and possibly seeding a cancer. Conversely, a stem cell with reduced division that cannot contribute efficiently to the physiological function of the stem cell community tends to be eliminated quite rapidly. We found experimentally that there is an analogous effect of division rate on the likelihood of becoming an FSC during pupal development. Thus, very few precursors with reduced division due to *cycE* or *cutlet* mutations became FSCs (Fig. 11B), while a greater proportion than normal became FSCs as a consequence of faster division due to excess CycE or increased JAK-STAT or Hh signaling (Fig. 11C). The likely rationale for this outcome is similar to the situation for adult FSCs. Precursors in the A/P territory of a developing germarium that may become FSCs are continually depleted during pupation by irreversibly becoming FCs (Fig. 11A). This loss is potentially balanced by influx from more anterior positions and by precursor division. In the adult germarium there is no net flux of ECs to become FSCs (Kalderon 2022). It appears that the major means of FSC precursor survival during pupal development is also through replication, even though there is net posterior expansion of the germarium early in pupation and hence, the potential for substantial replenishment of the FSC precursor domain from the anterior. Thus, the competitive advantage of a faster-dividing potential FSC actually initiates even prior to FSC formation, likely beginning at pupariation. The genetic factors that regulate division rate may differ between pupal and adult stages but it appears from our studies that several are in common (*cycE*, *cutlet*, JAK-STAT and Hh signaling) (Fig. 11B, C). These findings can explain the stronger effect on FSC lineage prevalence of mutations that increase or reduce division rates when induced in larvae rather than adults, as noted long ago for alterations to Hh signaling (Vied *et al*. 2012). An analogous principle may hold for some human stem cells with division-dependent differentiation, allowing the amplification of potential cancer-initiating mutations even before adulthood or full tissue maturation.

### Regulation of cell division rate during pupation

There is prominent spatial regulation of somatic ovary precursor cell division rate during pupal development, with EdU incorporation and FUCCI cell cycle reporters providing evidence of lower rates and early cessation of division anteriorly (Reilein *et al*. 2021). Indeed, this spatial regulation of division is potentially instrumental in setting the appropriate number of adult ECs. Unfortunately, we can only measure division rate in selected regions and with some limitations during pupation, counting ECs in EC-only lineages, the ECs and FSCs of EC/FSC/FC lineages, and FCs in terminal FC lineages. These measures sufficed to provide reliable evidence of reduced or increased division in response to genetic manipulations but cannot provide precise details of changes in the spatial distribution of division rates. Thus, we have evidence of decreased division for loss of function alterations to *cycE*, *cutlet*, *smo* and *yki*, together with increased division for excess CycE, activated Yki, *ptc* and increased JAK-STAT signaling, measured over the whole duration of pupation. Very little information was revealed for alterations, like loss of *stat* and excess Wnt pathway activity, which resulted in the production of almost exclusively ECs.

Since Hh pathway activity had a positive effect on precursor division and Hh pathway activity is known to decline from anterior to posterior in pupal ovaries (Lai *et al*. 2017), this pathway cannot contribute to the greater division rate of posterior cells. Excess JAK-STAT activity produced lineages with very large numbers of marked cells, partly obscuring the likely A/P position of the original precursor. Nevertheless, there were very few examples of the highly prevalent (>50%) typical wild- type EC-only clone with just 2-3 labeled cells. It seems clear, therefore, that excess JAK-STAT signaling increased division of even the most anterior precursors. Thus, it is surely important to limit JAK-STAT pathway activity in anterior precursors. Once an egg chamber has budded to produce polar cells, which are a key source of JAK-STAT pathway ligand, we observed strong pathway reporter signal in all egg chambers and the posterior half of the germarium, as in adults (Vied *et al*. 2012; Melamed and Kalderon 2020). It therefore seems inevitable that, as in the adult, JAK-STAT pathway activity promotes division of posterior FSC/FC precursors but not anterior EC precursor germarial cells during the second half of pupation.

A key question is whether JAK-STAT is an important stimulator of FSC or FC precursor division over the first 48h. There were few or no FSC- or FC-containing *stat* mutant lineages to measure cell numbers directly. Moreover, the severe loss of *stat* FSC lineages, which might be caused by reduced division rates, by anterior precursor movement or both, could have occurred during the second half of pupation. We did not see a strong JAK-STAT pathway reporter signal prior to 30h APF. Thereafter, however, reporter signal was stronger in posterior germarial cells and extra-germarial cells than in the anterior half of the germarium (Fig. 7), so it is plausible that graded JAK-STAT pathway activity may also contribute to faster division of more posterior cells over the first half of pupation (Fig. 12).

Inferred division rates of germarial pupal precursors are roughly inversely correlated to Wnt pathway activity reported by Fz3-RFP. Thus, EC precursors have low or undetectable pathway activity only early in pupation while anterior precursors are still dividing; more posterior precursors consistently have only low Wnt pathway activity and do not arrest. Loss of Wnt pathway activity (*arr*) did not, however, increase the average size of EC-only lineages or EC/FSC lineages. Almost all *axn* mutant precursors become ECs, so it is not known whether abnormally high (and potentially supra- physiological) Wnt pathway activity would impede posterior precursor cell division, as found for adult FSCs (Melamed and Kalderon 2020). Thus, despite the spatial correlation between higher Wnt signaling levels and lower division rates, we have no direct evidence of Wnt signaling reducing precursor division rates during pupation.

### Wnt signaling promotes anterior migration

We found strong and consistent alterations in precursor outcomes for genetic alterations of Wnt pathway activity. Loss of activity caused an extensive posterior shift in outcomes, with a large increase in lineages containing terminal FCs and only FCs, together with a large reduction in lineages containing only ECs or containing any r1 ECs. The frequency of lineages containing FSCs was substantially increased and an unusually high fraction of FSC lineages did not include any ECs. This is an interesting outcome: loss of Wnt signaling causes increased FSC loss in adults (because of greater conversion to FCs and apparently no replenishment from the anterior) but increased FSC production during pupal development. This can be rationalized on the basis that there are normally many more EC-only than FC-only precursors (63% vs 17%), so more cells have the potential to enter FSC- producing regions from the anterior than from the posterior (Fig. 11D). Moreover, continued posterior movement appears to convert many potential EC/FSC/FC lineages to FSC/FC lineages. The resulting high frequency of FSC/FC lineages plausibly constitutes a distinguishing characteristic of genotypes that promote posterior migration without altering division rates for a given location (Fig. 10J and 11D).

Increased Wnt pathway activity produced strong phenotypes, which were broadly the converse of those observed for loss of Wnt pathway activity (more EC-only lineages; fewer FC-only and terminal FC-containing lineages). Amongst EC outcomes, there was only a small shift in favor of r1 ECs (contrasting with the large converse effect of loss of pathway activity), suggesting that precursor migration in the most anterior regions of the developing germarium, where Wnt signaling appears to be roughly uniform, may not normally be driven by differential Wnt pathway activity.

There was also a reduction in *axn* FSC-containing lineages, all of which included ECs, from 15% to 6%. If anterior displacement caused all prospective FSC precursors to become ECs and all FC-precursors to form FSC-containing lineages, there would be a net gain in FSC lineages because the initial FC-only precursor pool is larger than the FSC precursor pool. Since FSC lineages are depleted we might infer that FC-only precursors migrated beyond the FSC domain to form EC-only lineages. An additional contribution to FSC loss could be a reduction in cell division rate. There were too few FC- and FSC-containing *axn* lineages to assess division rate in these posterior regions. However, adult FSC division is severely reduced by loss of *axn* (Reilein *et al*. 2017; Melamed and Kalderon 2020) and the properties of FSC precursors during the last 2 days of pupation are likely highly similar to adult FSCs because they appear to exist in a very similar environment. Hence, some contribution of reduced division rate to the observed paucity of FSC-containing lineages is likely.

We attempted to measure the temporal effects of Wnt pathway activity by initiating genetic alterations at different times during pupation. These results cannot easily be interpreted quantitatively but indicated directly that Wnt signaling influenced outcomes in the latter stages and indirectly (inferred from greater severity when initiated early) that Wnt signaling levels were also influential early in pupation. Also, since terminal FC fates were greatly reduced for *axn* lineages induced at pupariation but not when induced 36h later, we infer that the primary deficit is likely migration of precursors posteriorly out of the germarium into the EGC and basal stalk during the first 48h of pupation. Thus, terminal FC formation relies on limiting Wnt pathway activity (Fig. 11E), as found for FC production from adult FSCs (Reilein *et al*. 2017; Melamed and Kalderon 2020). Throughout pupal development, Wnt pathway activity, reported by Fz3-RFP, declined to undetectable levels roughly coincident with the anterior extent of strong Fas3 expression, which appears to mark FC precursors. Thus, very low or absent Wnt pathway activity appears to be a significant factor in allowing FC formation throughout pupation, as in adults.

### JAK-STAT signaling promotes posterior cell migration

The situation for JAK-STAT pathway signaling is roughly the mirror-image of that described for Wnt signaling, with a couple of significant differences (Fig. 11). First, increased JAK-STAT activity greatly stimulated precursor division. Second, although increased JAK-STAT activity (together with *UAS-Dap* to block changes in cell division rate) reduced the incidence of lineages with r1 ECs, loss of *stat* activity did not have the converse effect. Thus, even the most anterior precursors can be shifted more posterior in response to JAK-STAT but slightly more posterior EC precursors do not rely on JAK-STAT pathway activity to prevent anterior movement (unlike FC precursors and FSC precursors). The most likely explanation is that JAK-STAT activity is low, as observed directly, and functionally insignificant in the anterior half of the germarium throughout pupation. Interestingly, as for *arr* mutations, there was an increase in FSC-containing lineages lacking marked ECs for the *UAS-Hop UAS-Dap* genotype, consistent with the phenotype originating from posterior migration from EC-producing regions, rather than an increase in division rate (there was no indication of increased division for this genotype).

The large deficit of *stat* mutant lineages with marked terminal FCs (Fig. 11E) was a particularly important finding. The deficit was not observed for lineages induced 36h after pupariation and therefore appears to originate from a failure of precursors to migrate posteriorly out of the germarium over the first 48h of pupation. This is consistent with JAK-STAT pathway activity universally favoring FC formation in adults and pupae. The *stat* mutant phenotype suggests that low Wnt pathway activity, as seen in the most posterior pupal precursors, is not sufficient for normal terminal FC production, although the condition of eliminating both Wnt and JAK-STAT pathway simultaneously has not been directly tested.

In adults there is always a nascent or just-formed egg chamber budding from the germarium with anterior polar cells producing a JAK-STAT ligand that produces a posteriorly-biased gradient in the vicinity of FSCs to promote FC formation (Vied *et al*. 2012). Prior to formation of the first egg chamber there are no polar cells as a source of ligand. The genetic results show that STAT must have significant activity during this period because very few precursors lacking *stat* activity become terminal FCs. The small number of mutant terminal FCs formed were found in different locations in different ovarioles. Had all marked FCs been found, for example, in the basal stalk, that might have suggested that the most posterior precursors do not need to respond to a JAK-STAT signal to maintain their position and might therefore be the source of JAK-STAT pathway ligand.

We were able to detect pathway activity from 30h APF onwards. Activity was strongest in regions surrounding and posterior to the most posterior germline cyst, consistent with the possibility that this heightened activity promotes posterior migration of precursors out of the germarium prior to production of the first egg chamber. It will be important to determine which cells might be a source of ligand for the JAK-STAT pathway in the period prior to polar cell production. The pattern of pathway activity at 48h APF is consistent with the possibility of production of ligand by Fas3-positive cells (and/or efficient passage through Fas3-positive cells) and is similar, albeit weaker, to the pattern in adult germaria. That observation raises the question of whether the graded JAK-STAT pathway activity in adult germaria results entirely from ligand produced in polar cells or might also be contributed by early Fas3-positive FCs. Selective inhibition of Upd ligand production in adult polar cells using *neur-GAL4* strongly reduced STAT-GFP with partial penetrance (around 70%), while inhibition with *109-30-GAL4*, which is expressed in all early FCs and slightly higher in polar cells, was also effective (Vied *et al*. 2012). Those results argue strongly for a primary role of polar cells but also allow the possibility that early FCs may provide an additional source of JAK-STAT pathway ligand in adult germaria.

### Hedgehog signaling and Yki as an intermediate; division rate and some location effects

The contributions of Hh signaling and Yki activity to the behavior of adult germarial cells are relatively clear. Both Hh signaling and Yki promote FSC division and are required to limit apoptosis, while both are necessary in ECs to limit BMP production and hence allow normal germ cell maturation. In FSCs, Hh pathway activity can increase Yki activity through transcriptional regulation of *yki* (Huang and Kalderon 2014), while in ECs different reports suggested independent actions of Hh and Yki (Huang *et al*. 2017) or post-transcriptional regulation of Yki activity by Hh (Li *et al*. 2015).

Manipulation of Hh signaling and Yki activity during pupal development also produced some related phenotypes, reminiscent of effects on adult FSCs. Both increased Hh signaling (*ptc* mutation) and increased activated Yki (*UAS-YkiS168A*) expression increased measures of precursor division and increased FSC representation, as would be predicted as a consequence (Fig. 11C). However, Hpo pathway inactivation (via *hpo*, *wts* and *kibra*), which is expected to activate Yki post-transcriptionally, did not appear to increase division or FSC representation consistently. The increase in semi-quantitative measures of precursor division and of FSC representation observed for *ptc* were mostly reduced when *tub-yki* substituted for *yki*, suggesting that the increased division rate may, as in adult FSCs, depend on transcriptional induction of *yki*. However, there were exceptions, limiting the evidence that responses to *ptc* require normal transcriptional regulation of *yki*. Moreover, the inferred division rates and overall distribution of cell types for the *yki; tub-yki* genotype were quite similar to controls, suggesting that transcriptional regulation of *yki* activity can be substituted quite well by a constitutive promoter, as in adult FSCs.

There was a clear loss of *yki* mutant lineages that was suppressed by expressing DIAP1 to inhibit apoptosis but this appeared only to affect posterior precursors giving rise to FC lineages. The paucity of *yki; UAS-DIAP1* FSC-containing lineages was likely due primarily to reduced precursor division because it was suppressed by adding excess CycE together with DIAP1 (in lineages initiated 2d before pupariation to allow timely initiation of expression of the *UAS*-driven transgenes in MARCM lineages), strongly suggesting that Yki normally contributes to requisite precursor division rates.

A variety of other observations suggest some additional contributions of Hh signaling and Yki activity but these deductions are tempered by the limited magnitude or significant variability in some of the noted effects. First, there did appear to be some consequences for A/P movement of precursors. Loss of *yki* in the presence of *UAS-DIAP1* did not result in any notable changes of representation of r1 ECs but did reduce the representation of terminal FCs in lineages initiated 2d before pupariation. Increased Yki activity consistently (*kibra*, *wts*, *hpo*, *UAS-YkiS168A*) reduced r1 EC representation but only *UAS-YkiS168A* modestly (and not significantly) increased terminal FC production. Increased Hh signaling modestly decreased r1 EC representation, while increasing terminal FC representation. The effect on r1 ECs was retained when *yki* was substituted by *tub-yki*, while the effect on terminal FC representation was lost. This suggests that increased Hh signaling reduces r1 EC production independent of Yki, even though increased Yki activity has the same effect, while terminal FC representation, which is modestly increased by *UAS-YkiS168A* and decreased by loss of *yki*, is only increased by excess Hh signaling when transcriptional regulation of *yki* is allowed. These deductions align with earlier conclusions about adult ovaries: Hh signaling acts through Yki to influence more posterior cells (FSCs), while Hh and Yki act independently, albeit with similar consequences, for anterior cells (ECs) (Huang and Kalderon 2014; Huang *et al*. 2017).

The interpretation of *smo* phenotypes, representing loss of Hh pathway activity, has an additional caveat. Clone frequency was lower than for controls and was not restored by *UAS-DIAP1*, even though lineage phenotypes were altered by DIAP1. Thus, there is significant precursor loss or death (either non-apoptotic or apoptosis is not efficiently rescued by *UAS-DIAP1*) with unknown spatial selectivity. In *smo; UAS-DIAP1* lineages there was a loss of FSCs, plausibly due to limited cell division (Fig. 11B). This would be expected to increase the frequency of FC-only or EC/FC lineages. FC- only lineage frequency was increased but beyond that expected from FSC loss alone, and r1 EC representation was modestly reduced, with no change in terminal FC representation. Thus, loss of Hh signaling, at least among surviving cells, appears to cause a modest posterior shift over most of the precursor domain, excluding the most posterior regions. This deduction stands alongside a deduction of (stronger) posterior shifts also for increased Hh signaling. The latter deduction relates to potential consequences of Hh signaling, whereas the former reports the normal influence of Hh signaling. Thus, with significant caveats, it appears that Hh signaling might normally promote retention of anterior positions for most of the precursor domain. This is easily reconciled with the anterior bias of Hh pathway activity and the likelihood that posterior precursors soon fall out of the range of Hh signaling.

Another study examined the consequences of manipulating Hh pathway activity in all somatic cell ovary precursors (rather than clonally) and concentrated mainly on genetic changes initiating prior to pupariation, to conclude that Hh signaling promotes retention of ICs in the developing gonad (Lai *et al*. 2017). It is conceivable that the shortfall of *smo* lineages we observed is due to migration of some precursors away from the developing ovary. However, such losses might plausibly affect the most posterior precursors most strongly, and we found no selective shortage of terminal FC representation in *smo; UAS-DIAP1* lineages. Hence, we favor limited cell death as a more likely explanation. The effects of Hpo pathway inhibition (to increase Yki activity) have also been reported for early ovary development using manipulation of gene activity in all precursors, leading to a conclusion that the IC pool was increased in response, plausibly due to increased division (Sarikaya and Extavour 2015). This result is commensurate with our deductions from clonal analyses with UAS- YkiS168A and loss of *hpo*, though neither *kibra* nor *wts* mutations elicited a discernible indicator of increased division.

### Summary conclusions and open questions

In the adult ovary, FSCs are maintained within a narrow A/P domain. At the posterior edge of that domain, some FSCs move posteriorly into territory with lower Wnt signaling and higher JAK-STAT signaling to become FCs (Fig. 12). Genetic reduction of Wnt or increase of JAK-STAT signaling greatly increase the chance, cell autonomously, of an individual FSC becoming an FC, suggesting that the magnitudes of these pathways are major drivers of FSC movement (Vied *et al*. 2012; Reilein *et al*. 2017; Melamed and Kalderon 2020). The converse genetic argument holds for FSC conversion to ECs but the frequency of EC production is much lower than for FCs, perhaps because FSCs less frequently stray into more anterior territory, which is stably occupied by quiescent ECs, while space to the posterior of FSCs is constantly vacated by the process of egg chamber formation and budding. We previously found that the nature or behavior of somatic ovarian precursor cells, like FSCs, appears to be guided solely by location (Reilein *et al*. 2021). Thus, the precursors that become FSCs are simply those that end up in the location of FSCs at the end of pupation.

Here, we found that the external signals guiding the location of precursors during pupal development overlap extensively with those used in adults (Fig. 12). This is relatively easily rationalized for Wnt signaling because Cap cells, located at the anterior of the germarium, are already differentiated at pupariation and appear to be a source of Wnt signals throughout pupation and adulthood. It is likely that an additional source of Wnt arises progressively during pupation as EC precursors stop dividing and adopt mature cell characteristics of producing Wnt ligands (Sahai-Hernandez and Nystul 2013; Wang and Page-mccaw 2018) while the distance from Cap cells to FSC and FC precursors increases (Fig. 12). The net consequence is that from early pupation onwards there is graded Wnt signaling that declines to undetectable levels at the point of FC specification, which coincides with the onset of strong Fas3 expression (Fig. 5).

The involvement of JAK-STAT signaling was not predictable because a major source of ligand in adults are the anterior polar cells associated with an egg chamber budding from the germarium (Fig. 12). The first egg chamber produced during pupation has no antecedent and there is therefore no polar cell source of JAK-STAT ligand as a subset of somatic cells move posteriorly out of the developing germarium during the first 48h to acquire strong Fas3 expression and become the first FCs. We nevertheless detected JAK-STAT pathway activity around this precursor to FC transition zone and found that formation of the first FCs is strongly dependent on JAK-STAT activity, just as it depends on low Wnt activity. We do not know the source of the JAK-STAT ligand or whether pathway activity is graded over this transition zone, though it is clearly lower in more anterior regions. Hh pathway activity declines from anterior to posterior in both pupal and adult ovaries (Forbes *et al*. 1996; Vied *et al*. 2012; Lai *et al*. 2017). Although there were some effects of manipulating Hh signaling on A/P precursor outcomes (contrasting with adult studies, where only cell division rate appears to be affected), these were much weaker than observed for Wnt or JAK-STAT pathways, suggesting that the latter two pathways are the major determinants of developmental formation of ECs, FSCs and FCs, just as they have the greatest influence on FSCs becoming ECs or FCs or remaining as FSCs in adults.

The factors regulating proliferation of pupal precursors also were found to be very similar to those regulating adult FSC division rate, although the developmental data are much more limited and semi-quantitative at best. JAK-STAT and Hh signaling, via *yki* induction, are major stimulators of cell division, while Wnt signal normally has no significant influence (Fig. 12). The role of cell division rate in producing FSCs during pupal development was also found to be analogous to its role in maintaining FSCs. In both situations, it appears that the major loss of cells in a location with potential to act as an FSC is from posterior movement to become FCs. Although some cells may move from an EC-centered precursor domain to final FSC positions during pupal development, the maintenance of precursors that become FSCs and of adult FSCs themselves, despite posterior loss, both depend principally on a sufficient cell division rate. Hence, both the formation and the maintenance or expansion of FSC lineages is favored by genetic cell-autonomous changes that increase division rate (Fig. 11).

We suggest that two major findings of our study are likely to apply to other stem cell paradigms that share major characteristics with FSCs (Fig. 12). First, if stem cells of a specific tissue are specified during the entire period that the rest of the tissue is specified, rather than being set aside early during development, it is likely that the same signals contribute to stem cell development and stem cell maintenance in adults. Second, in such scenarios where, additionally, the division and differentiation of adult stem cells are separate processes, the probability of an appropriately located precursor becoming a stem cell and of an adult stem cell being maintained both depend on the division rate of that cell relative to competitors vying to become stem cells or remain as adult stem cells.

## Data Availability

Drosophila stocks used in this study are available upon request. The authors affirm that all data necessary for confirming the conclusions of the article are present within the article, figures, and supplementary spreadsheets deposited with the article.

## Acknowledgments

We thank the Bloomington Drosophila stock center, the Developmental Studies Hybridoma Bank (DSHB), and Laura Johnston (Columbia University) for reagents, FlyBase as an information resource, and the confocal microscope resource provided by the Department of Biological Sciences, Columbia University. We thank Aron Choi, Brian Heubel, Pegah Khosravi-Kamrani and Karen Sophia Park for experimental assistance, David Melamed, Hoyon Kim, Laura Johnston and Iva Greenwald for discussions.

## Study funding

This study was funded by NIH grant RO1 GM079351 to DK.

## Supplementary Materials

**Supplementary Figure 1: Wnt signaling influences over different periods of pupation.**

**Supplementary Figure 2: JAK/STAT signaling in pupal germaria after stalk and polar cell formation.**

**Supplementary spreadsheets with raw data and calculations underlying all presented data**

**Single-cell 0hAPF calculations**

All summary clone data for lineages induced at 0h APF together with the calculations converting data to estimations of single-cell lineage data and averaging of all control values and other genotypes tested in more than one experiment.

**Single-cell -2dAPF calculations**

All summary clone data for lineages induced at -2d APF together with the calculations converting data to estimations of single-cell lineage data and averaging of all control values and other genotypes tested in more than one experiment.

**0h APF Significance Fisher and -2d APF Significance Fisher**

Copies of the two previous spreadsheets (numbers only, no formulae exposing the calculation methods) with additional columns to show p values of significant differences to controls if p<0.05 using Fisher’s exact two-tailed test.

**0h APF Controls Variation**

Calculation of SD and SEM for single-cell lineage parameters among control experiments and comparison to expectations from probabilistic labeling of precursors in different A/P locations.

**0h APF Graphs Fig1,3**

First sheet is 0h APF master sheet (from the single-cell calculation sheet). Others show data for Figs. 1B-D and Fig. 3I.

**0h APF Graphs Fig4-10**

First sheet is 0h APF master sheet (from the single-cell calculation sheet). Others show data for Figs. 4, 6, 8H, 9D, 10G

**-2d APF Graphs**

First sheet is -2d APF master sheet (from the single-cell calculation sheet). Others show data for Figs. 3J, 8I, 9E.

**-3.5d axn August 2023 Numbers**

Control and *axn* lineages initiated 3.5d before eclosion (36h APF); raw data and deduced frequencies of clone types.

**-4d arr DIAP Numbers**

Control and *arr; UAS-DIAP1* lineages initiated 4d before eclosion (24h APF); raw data and deduced frequencies of clone types.

**Fig. S1 Graph Data**

Data for Supplementary Fig. 1.

**Fig5 Fz3RFP Graph**

Data for Fig. 5 graph of Fz3-RFP Wnt reporter intensities over the germarium at different times.

**-3.5d stat November 2021 Numbers**

*stat* lineages initiated 3.5d before eclosion (36h APF); raw data and deduced frequencies of clone types.

**FC occupancy aggregate**

Raw data and aggregation for fraction of terminal egg chamber occupied by marked FCs and frequency of additional FC labeling in penultimate or more anterior egg chambers for cycE and cutlet genotype lineages (initiated at 0h APF and -2d APF, leading to graph in Fig. 3F), and for selected Hedgehog and Yorkie genotypes (0h APF Hh Yki TermFC). Raw cell number data for EC-only and EC/FSC lineages of select Hh Yki genotypes to estimate SEM (Cell#). Summary data for fraction of available cysts with marked FCs in FSC-containing lineages (%cystsMarkedFCs 0h APF).

**Division Measures Hh/Yki**

Summary data for graphs in Fig. 10H-K.

Raw data lineage spreadsheets: “**Compilation of Numbers**” named by Figure and time of lineage initiation (“-5d” = 0h APF, “-7d” = -2d APF).

First sheet in a pair is the raw tabulation of marked cells for each sample of a given genotype. The second sheet compiles the raw data to give a variety of measures, including those transferred to the master spreadsheets used to estimate single-cell lineage parameters (“calculations” spreadsheets).

## Notes

### Competing Interest Statement

The authors have declared no competing interest.

### Summary of Updates

Title and Abstract changed. Additional author added. Several Figures and associated text revised. Two summary model figures added. General significance of the findings for this paradigm has been added or expanded in the Discussion, end of Abstract and end of Introduction. Some added data required additional supplementary spreadsheets with corresponding raw data and/or calculations.

