## Supplemental Figures for "Regulation of somatic stem cell precursor fates and proliferation during *Drosophila melanogaster* pupal ovary development resembles the signaling framework for adult stem cell behavior"

### Supplementary Figures

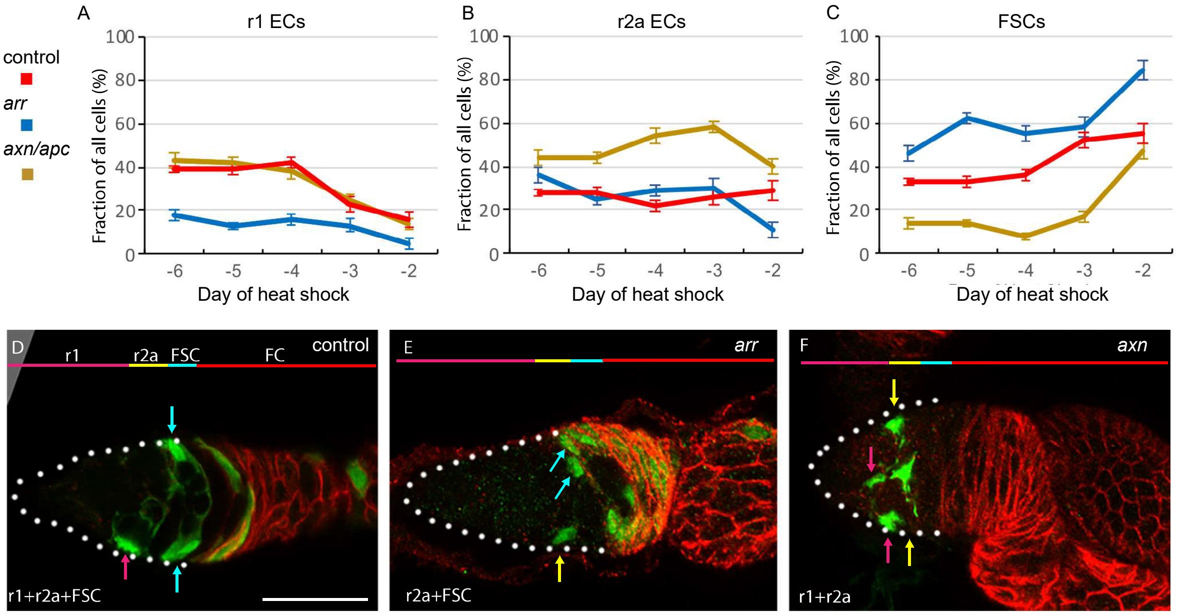


**Supplementary Figure 1: Wnt signaling influences over different periods of pupation.**
**(A-F)** MARCM lineages with genetically altered Wnt pathway activity were induced from 6d to 2d before eclosion and examined in 2d-old adults. (**A-C**) The total number of marked cells of each cell type (r1 or r2a ECs and FSCs) was counted over all samples and expressed as a percentage of all marked ECs and FSCs for control clones (red), *arr* mutant clones lacking Wnt pathway activity (blue) and *axn* or *apc* mutant clones (combined data, gold) with increased Wnt pathway activity. **(D-F)** Images of germaria stained for Fas3 (red) show examples of a **(D)** control lineage with all cell types labeled, **(E)** an *arr* lineage with marked r2a ECs, FSCs and FCs **(F)** an *axn* lineage with only marked r1 and r2a ECs. Scale Bar, 20 µm. White dotted line shows outline of germaria. For raw data please see supplementary spreadsheet titled Fig S1 Graph Data.

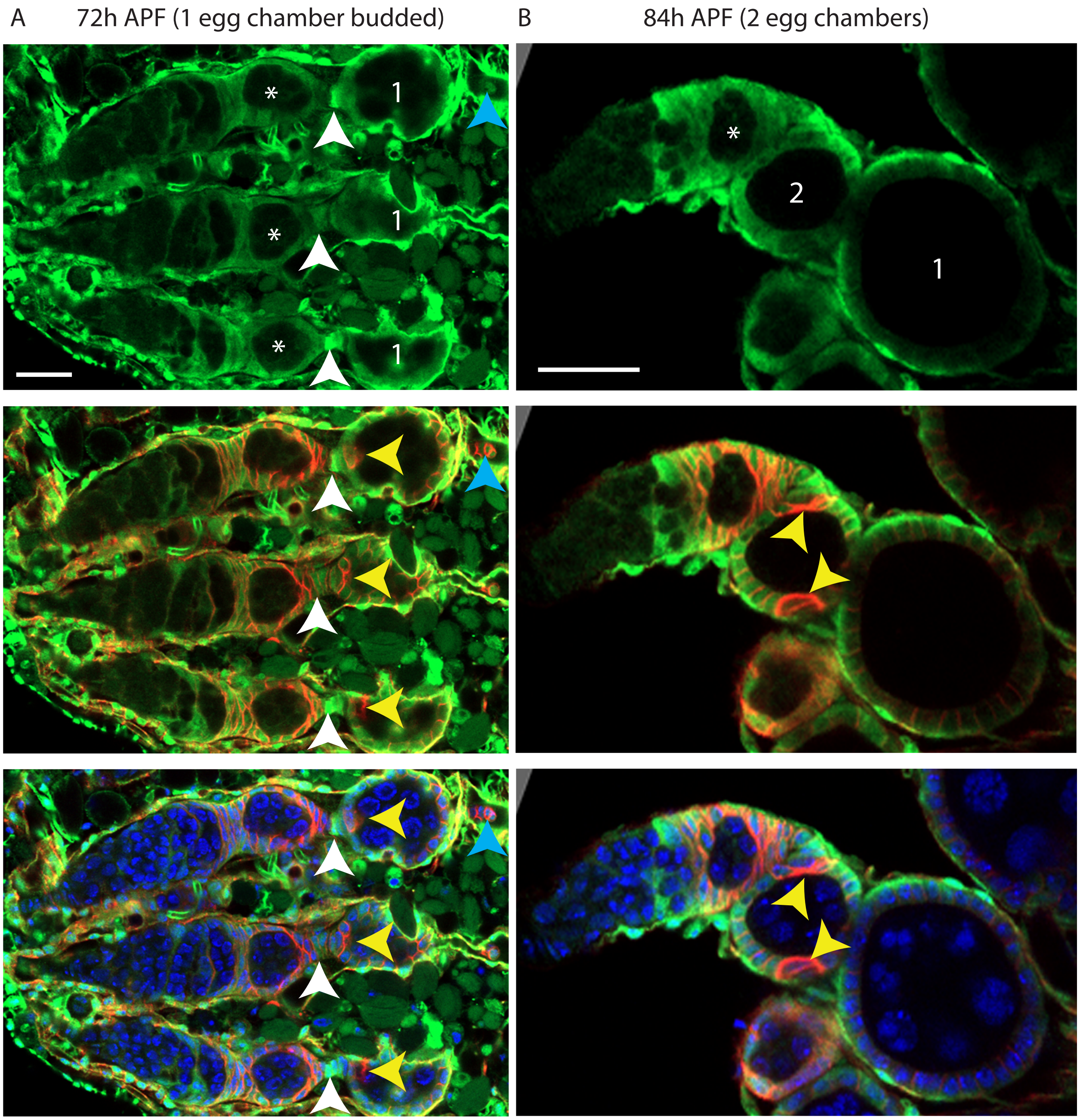


**Supplementary Figure 2: JAK/STAT signaling in pupal germaria after stalk and polar cell formation.** Pupal ovaries expressing STAT-GFP were stained for GFP (green), Fasciclin 3 (red), and with DAPI (blue). Bars, 20mm. **A)** Image from a pupal ovary at 72h APF, after the first egg chamber has budded (labeled with a 1), polar cells have formed on the first egg chamber (yellow arrowheads indicate the anterior polar cells), and a stalk has formed between the egg chamber and the germarium. Epithelial sheath cells that surround each developing ovariole and the entire ovary strongly express STAT-GFP. STAT-GFP is also expressed in follicle cells in the egg chamber and the germarium, and around the cysts just anterior to the Fas3 border. The most mature egg chamber in the germarium is marked by an asterisk. STAT-GFP is often prominent in stalk cells (white arrowheads) and is also detected in the basal stalk (blue arrowhead). **B)** By 84h APF, two egg chambers have budded (labeled with 1 and 2), and STAT-GFP expression resembles the adult pattern, where it is expressed in follicle cells of the egg chambers and the posterior of the germarium, and tapers off anterior to the Fasciclin 3 border. Yellow arrowheads indicate the polar cells in the $2^{nd}$ egg chamber and the asterisk marks the most mature egg chamber in the germarium.
